# Heterogeneity of functional cellular properties for neurons in mouse cerebral cortex

**DOI:** 10.1101/2025.03.25.645203

**Authors:** Julien Ballbé, Margaux Calice, Marta Gajowa, Lyle J. Graham

## Abstract

Biological systems are known to exhibit a high degree of heterogeneity in their constituent components and their organization, and neuronal systems are no exceptions. To understand the functional impact of this heterogeneity in the brain, network models need to consider how this is manifested at the level of cellular properties, and thus how to consider variability in reported experimental data. Many studies have pointed out the variability in neuronal density, structural organization or synaptic connectivity across different neuronal networks and populations. Similarly, neuronal physiological properties are known to greatly vary across neuronal populations. Yet, the characterization of electrophysiological diversity has mainly relied on descriptions of firing properties (e.g. bursting, spike frequency adaptation) with various quantitative definitions of the boundaries between neuronal classes (e.g. fast spiking, regular spiking). Furthermore, lab specific implementations of experimental design and data analysis are an obstacle for comparisons between studies. In this context, the quantitative consideration of neuronal variability across commonly accepted neuronal classes provides an objective approach to describe neuronal physiological heterogeneity.

We analyzed several publicly available databases to characterize the variability of linear and input/output properties of cortical neurons, according to multiple factors covering the entire cortical neuronal population. We assessed the variability of the main cortical neuron types (Excitatory, PValb, Sst, Htr3a, Vip), revealing their heterogeneity as function of cortical area (primary visual, motor and somato-sensory areas), including between layers within a given area. Our comparative database study revealed that different experimental conditions (e.g., *in-vitro* vs. *in-vivo*, recording temperature) can influence the properties of any given cell type, while preserving overall differences between types. We find that considering the input to a given neuron in terms of the effective voltage response of a linear model can account for some of the heterogeneity of I/O properties, and suggest that these properties are directly linked to cell input resistance, thus cell size. This works constitute a strong foundation for the consideration of detailed neuronal electrophysiological heterogeneity in future large-scale modeling works.

## 2 Introduction

Computational models of cortical networks are an essential tool for understanding their functional properties, establishing the causality between the biophysics of the constituent neurons and emergent network dynamics. For large scale network models which explictly consider action potentials, minimal representations of neurons instantiate low dimensional input/output (I/O) relationships, for example reducing complex dendritic morphologies to single point cells, retaining at least an explicit definition of the threshold input for action potential generation and the I/O gain for larger inputs. With respect to spikes, more elaborate cellular models may take into account simple modulation of spiking output including adaptation and saturation of spike frequency, or more complex dynamics such as bursting. With respect to different types of neurons, the simplest network models of cerebral cortex typically include two cell types (e.g. excitatory and inhibitory) for a specific circuit, thus a given cortical layer in a given cortical area. More elaborate network models can consider multiple cellular types, multiple layers with cortex, and, ultimately, multiple cortical areas.

Emergent dynamics of these highly non-linear systems are strongly dependent on the quantitative details of the elements, and this motivates careful consideration of relevant parameters obtained from the biology. An important consideration is that these details for any given neuron type are not precise, and in principle the variation of any given parameter, indeed the overall parameter heterogeneity within a population of a given type, may contribute quantitatively if not qualitatively to network behaviour.

Large-scale models of mouse cortex provide an important context for a mechanistic understanding of the mammalian brain’s functional properties, allowing the exploitation of extensive anatomical and neurophysiological data and a relatively small network size. In that context, we collected datasets from several public domain electrophysiological databases of mouse cortical neurons that provide measures of their firing properties. We then made a comparative analysis of I/O heterogeneity over selected subsets of the datasets corresponding to experimental protocol, cell types, cortical layers and cortical areas.

What precisedly consitutes the definition of different neuronal types is an ongoing debate, yet also essential for unveiling the complexity of neuronal systems (Zeng, 2022). Neurons have been classified according to different modalities, with the aim of establishing a classification scheme that can eventually account for all neurons. The most evident distinction is between excitatory and inhibitory neurons, based on the type of neuro-transmitter they release. Neurons have also been classified according to their morphology (e.g. dendritic arborization, synaptic projection) (Defelipe, 2002; Markram et al., 2004). Neurons have also been classified physiologically, at the most basic level according to their response to a depolarizing current step sufficient to elicit trains of action potentials. This resulted in the consideration of different distinct type of cells. The first classification of firing patterns identified three main classes of neurons: regular spiking (RS), intrinsically bursting (IB), and fast-spiking (FS) neurons (Connors and Gutnick, 1990). Since then, other electrophysiological types have been identified (Contreras, 2004). Yet, these morphological and physiological cell type definitions are mostly qualitative, with no clear numerical boundaries to delimit cell classes. A more objective classification of neurons have been used over the past decades, relying on the cellular expression of diverse markers, like calcium-binding proteins like Paravalbumin (PValb) (Hendry and Jones, 1991; Hendry et al., 1989), the somatostatin peptide (Sst) (Rudy et al., 2011; Tremblay et al., 2016), or serotonine-receptor (Htr3a) (Morales and Bloom, 1997; Férézou et al., 2002). Together, these markers have notably been account to report, in a non-overlapping manner, for the entire GABAergic neuronal population of the cerebral cortex (Xu et al., 2010; Lee et al., 2010; Rudy et al., 2011; Tremblay et al., 2016; Rodarie et al., 2022), therefore representing a more objective cell classification prisms than electrophysiological or morphological classification. Over the last decade, this molecular taxonomy has been extended with the implementation of single-cell RNA sequencing (Tang et al., 2009), enabling to class cells according to their genetic profile. these studies provided more in-depth classification of cell types, and gained ever more attention as it also allowed to linked cell types to spatial location (Zeisel et al., 2015; Tasic et al., 2016, 2018; Gouwens et al., 2020; Williams et al., 2022). Overall the study of cortical neurons through these different modalities has brought up multiple insights. First, excitatory and inhibitory neurons are quite different in shape, electrophysiological properties and molecular identities. Where glutamatergic neurons constitute a relatively homogeneous group, recognizable by their large soma, and long-range projections (Defelipe and Fariñas, 1992; Mao and Staiger, 2024), interneurons present much more variability according to their molecular markers, electrophysiological profiles, and morphological features (Markram et al., 2004; Ascoli et al., 2008; Gouwens et al., 2019, 2020).

Interneurons types have been studied according to their molecular markers (i.e.: PValb, Sst, Vip, Htr3a), highlighting correspondences notably between molecular markers and electrophysiological properties. Thus, PValb neurons have been found to largely correlate with FS firing types. Furthermore, RS were not expressing PValb, but the other classes of molecular markers like VIP and Sst (Contreras, 2004). This dual analysis between molecular markers and electrophysiological characteristics, has been confirmed by recent work reveling that a set of properties measured from extensive recording protocols enabled to identify different electrophysiological types (e-types) which correspond to common molecular markers (Gouwens et al., 2019). Even though these associations are globally correct, the terms Fast-Spiking or Regular-Spiking encompass a variety of electrophysiological features expressed at different degree, and which highly depends on the way these are quantified. For example, FS cells are, among other, characterized by a little or no spike frequency adaptation (SFA), whereas it is highly expressed in RS neurons (Ascoli et al., 2008; Tremblay et al., 2016). If we consider the SFA to be measured similarly, even though there are no consensus, as the measure may highly be dependent on the data sampling (Benda and Herz, 2003; Peron and Gabbiani, 2009; Shinomoto et al., 2003, 2009; Mitriĉ et al., 2019), there are no biological limit as to what makes a SFA to be considered as small or big, other than purely qualitative and subjective observations. This example could be applied to other aspects of neuronal electrophysiological properties, like neuronal gain, rheobase, input resistance… It therefore appears that a more convenient and objective manner to electrophysiologically characterize neuronal classes (for example defined by there molecular markers), would be to compare the distribution of electrophysiological features

Furthermore, even molecularly defined classes of neurons have been observed to display some degree of intra-class heterogeneity. Notably, Vip expressing interneurons have been reported to express morphological and electrophysiological variability according to the cortical layer they belong to (Prönneke et al., 2015). They notably observed that layer II/III Vip neurons tend to be more depolarized at rest than Vip neurons from subgranular layers, as well as smaller after-spike hyperpolarization potential (AHP). Another study compared Sst expressing neurons’ electrophysiological, morphological and transcriptomic features in different cortical layers, and cortical areas (Scala et al., 2019). Their work notably pointed that these cell may differ in different electrophysiological phenotypes between layer IV and layer V in Visual and somato-sensory cortices. These layer-dependent electrophysiological properties within a same neuron type have been notably further confirmed by electrophysiological (Gouwens et al., 2019) and transcriptomic (Gouwens et al., 2020) clustering, which observed that e-types and t-types of both excitatory and inhibitory cells may be restricted to particular cortical layers. Yet, a more in depth and exhaustive review of the physiological properties of the different cortical neuron classes is still lacking and could be of great importance, notably in the context of large scale neuronal networks.

To better characterize and understand electrophysiological variability, we relied on the recently developed TACO pipeline (Ballbé et al in preparation) to gather different publicly available databases of current-clamp recordings. By analyzing cellular properties, according to different biological factors like neuron type, cortical layer, and cortical area, but also experimental like *in vivo/in vitro* or recording temperature, we aim to characterize the variability in neuronal properties by examining differences in commonly used electrophysiological measures and assessing how these features are distributed across different factors.

### 2.1 Gathering of multiple databases

The reusability of scientific databases is a major aspect of open sciences, notably to ensure reproducibility, and honest review of results (Wilkinson et al., 2016; Lamprecht et al., 2020). Despite the extensive use of intracellular electrophysiological recording since the works of Hodgkin and Huxley in the 30s, only recently have some laboratories started to implement solutions to promote inter-database interoperability, like common data format with the Neurodata Without Border (NWB) (Teeters et al., 2015; Rübel et al., 2022) or common data repositories as the DANDI repository (as part of the BRAIN Initiative) to facilitate exchange and availability of database. Yet, there are still a lot of intracellular current-clamp database not referenced in the DANDI repository or not relying on the NWB format. To enable the valorization of these databases, regardless of their structure, the TACO (Trace-based Analysis for Coherent and Organized database integration) pipeline (Ballbé et al, in preparation) has been designed, and proposes a set of methods to 1) extract raw electrophysiological data from database, 2) account for common electrophysiological recording artifacts like the bridge error, 3) automatically apply common set of user-defined quality criteria and 4) measure different input/output (I/O) relationship properties along with commonly defined linear properties, using data-based methods.

## 3 Methods

### 3.1 Databases

For our study, we gathered datasets of intracellular current-clamp patch recordings using long-step current inputs (≥ 500ms) from the following databases: Allen Cell Type Database (CTD, celltypes.brain-map.org), Scala 2019 ((Scala et al., 2019)), Scala 2020, and NVC (data collected by our laboratory) databases.

#### 3..1.1 Allen Cell Type Database

The Allen Institute for Neurosciences published over the last years multiple database of different modalities like transcriptomics, morphology and electrophysiology (Gouwens et al., 2019). This database of electrophysiological recording, present multiple advantages, as the use of detailed and varied current-clamp protocols (short-square, long-quare, ramp stimulus), and the detailed cell-level metadata information, like mice line, reporter status, and cortical layer in which it has been recorded. Mainly composed of visual cortex cells, it has been extensively used in different context notably for machine learning (Ghaderi et al., 2018; Rodríguez-Collado and Rueda, 2021; Ophir et al., 2024), transcriptomic analysis (Gouwens et al., 2020), and model generation (Teeter et al., 2018; Nandi et al., 2022). To select the cells of interest from this database, we targeted the cells recorded from transgenic mice line for which we could find references in the literature about the cells type they targeted (Excitatory, PValb, Sst, Htr3a or Vip) (See Table 1 and 2), and reported by the Allen Cell Type Database as expressing the cell reporter.

**Table 1:** Transgenic cell lines targeting excitatory neurons in the Allen Cell Type Database.

| Cell Type | Line Name | References |
| --- | --- | --- |
| Excitatory | Ctgf-T2A<br>-dgCre | (Tasic et al., 2016) |
| Excitatory | Cux2-CreERT2 | (Huang et al., 2021) |
| Excitatory | Glt25d2-Cre<br>_NF107 | (Kim et al., 2015)<br>(Tasic et al., 2016) |
| Excitatory | Nr5a1-Cre | (Nandi et al., 2022) |
| Excitatory | Rbp4-Cre<br>_KL100 | (Lee et al.)<br>(Nandi et al., 2022)<br>(Munz et al., 2023) |
| Excitatory | Rorb-IRES2<br>-Cre | (Tasic et al., 2016)<br>(Nandi et al., 2022) |
| Excitatory | Scnn1a-Tg2<br>-Cre | (Tasic et al., 2016)<br>(Zeng and Sanes, 2017)<br>(Nandi et al., 2022) |
| Excitatory | Scnn1a-Tg3<br>-Cre | (Tasic et al., 2016)<br>(Zeng and Sanes, 2017)<br>(Frantz et al., 2020)<br>(Nandi et al., 2022) |
| Excitatory | Tlx3-Cre<br>_PL56 | (Kim et al., 2015) |

**Table 2:** Transgenic cell lines targeting inhibitory neurons in the Allen Cell Type Database.

| Cell Type | Line Name | References |
| --- | --- | --- |
| Vip | Chat-IRES-Cre-neo | (Tasic et al., 2016; Zeng and Sanes, 2017) |
| Vip | Vip-IRES-Cre | (Tasic et al., 2016)<br>(Zeng and Sanes, 2017) |
| Vip | Vip2-IRES2-Cre | (Tasic et al., 2016)<br>(Zeng and Sanes, 2017) |
| Sst | Chrna2-Cre-OE25 | (Tasic et al., 2016) |
| Sst | Nos1-CreERT2 | (Tasic et al., 2016) |
| Sst | Sst-IRES-Cre | (Tasic et al., 2016) |
| PValb | Nkx2-1-CreERT2 | (Tasic et al., 2016) |
| PValb | PValb-IRES-Cre | (Tasic et al., 2016) |
| Htr3a | Htr3a-Cre<br>_NO152 | (Gerfen et al., 2013)<br>(Zeng and Sanes, 2017) |

#### 3.1.2 Scala 2019

This database (Scala et al., 2019), published on the Zenodo repository using Matlab file format, has originally been used to address morphological, electrophysiological and transcriptomic differences of Sst cells in different layers and cortices. This database is therefore composed of Patch-clamp recordings in visual and somato-sensory cortices, in either Layer IV or layer V. The neuronal classes was inferred from morphological attributes. More precisely, PValb cells were composed of large, small, double-bouquet and horizontally basket cells, while Vip cells were composed of bipolar cells with small soma extending to L1 and L5. Finally, non-Martinotti cells from somato-sensory cortex were considered as Sst cells (while in the visual cortex Sst interneurons are reported be predominantly Martinotti cells).

#### 3.1.3 Scala 2020

Scala et al. (2020), as part of the BRAIN Initiative Cell Census Network, introduced a comprehensive database to explore neuronal transcriptomic diversity in the motor cortex. Their study utilized the patchseq technique to record cells across various cortical layers from multiple transgenic mouse lines (PValb, Sst, Vip, and excitatory), under both room temperature, and physiological temperature. This *in vitro* approach revealed a correlation between morpho-electrical and transcriptomic variability, indicating that transcriptomic types may form a continuous spectrum rather than distinct, discrete families. The different cell types were documented directly in the cell files provided in the database. If the cell type contained ‘+’ (e.g., “SST+”) then we considered it as a cell expressing the cell reporter. Otherwise, we would not consider it belonging to any cell type.

#### 3.1.4 NVC database

In recent years, the host laboratory has compiled a database of *in vivo* “blind-patch” whole-cell recordings from multiple species, including cat, rat, and mouse. These recordings, in both current and conductance (dynamic) clamp configurations, do not provide cellular identification apart from physiological classification. This database is therefore a unique opportunity to study the impact of *in vitro/in vivo* configuration, as well as making the link between current clamp and conductance clamp measurements.

#### 3.1.5 Database summary

From these databases, we considered cells recorded from visual (VIS), somatosensory (SS) and motor (MO) cortices, for which the cell type and cortical layer were known (apart from the NVC cells, Figure 1), including a total of 2735 cells. When using the TACO pipeline to measure electrophysiological properties, the analysis was successful for 2730 cells (for 5 cells, the analysis failed due to untracked internal pipeline error), representing 112 208 traces. The TACO pipeline enables the user to apply user-defined quality criteria to every trace. To prevent any aberrant results, we applied in the TACO pipeline the following trace-based quality criteria: 1) The amplitude of the current trace should be lower than 5000pA, 2) the mean value of the current trace should be lower than 5000pA, 3) the amplitude of the voltage trace should be lower than 190mV, 4) the mean value of the voltage trace should be lower than 10mV and 5) the trace-based input resistance (i.e.: the voltage deflection divided by the current step) should be positive. After the application of the quality criteria, 1990 traces coming from 119 different cells were rejected, mainly because of a mean voltage trace higher than 10mV (Table 3).

**Figure 1:**
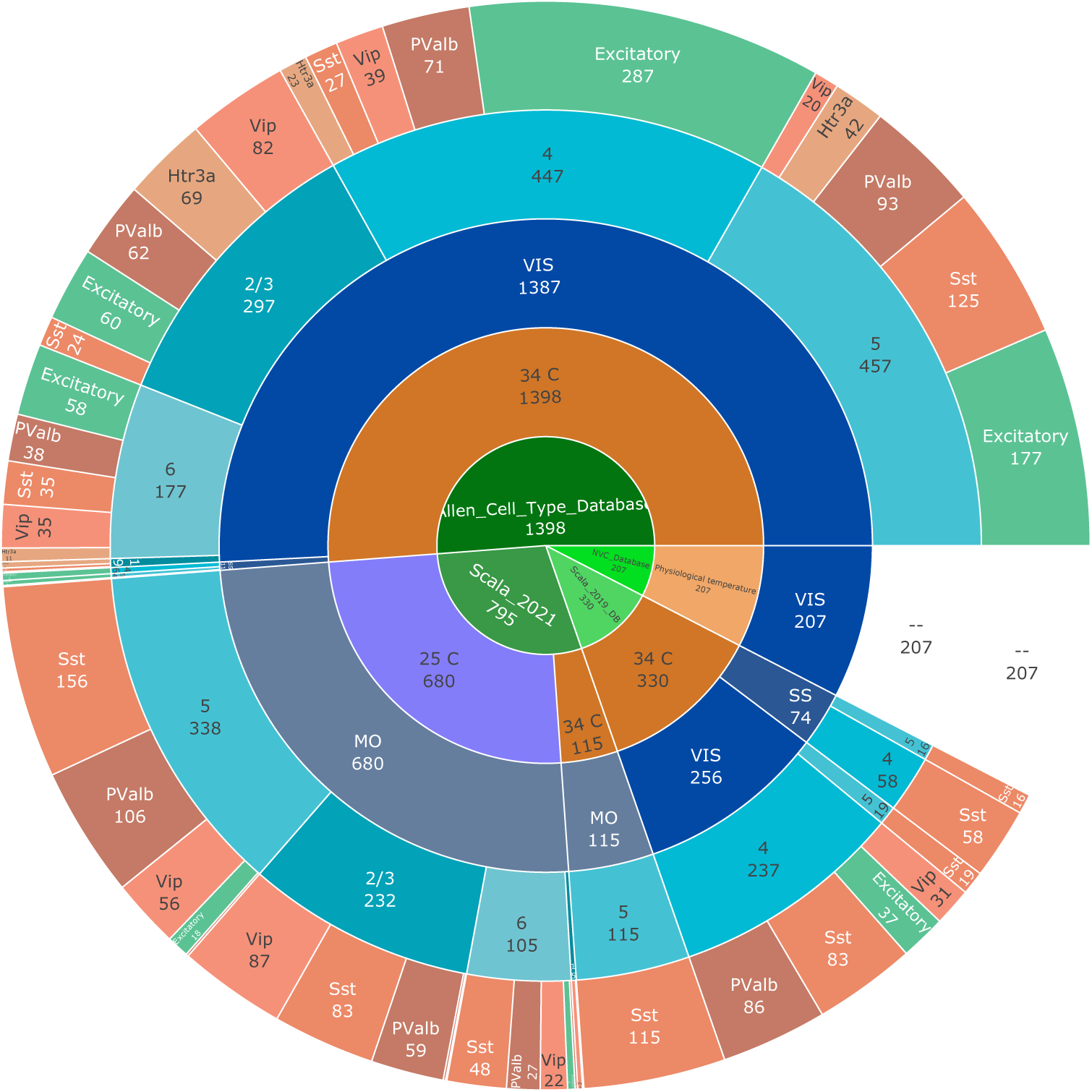
Number of cells gathered from different databases. We gathered four different databases. From which we kept only cells for which both the cortical layer and the cell type were known, except for the NVC database, whose interest relies on the *in vivo* protocol used.)

**Table 3:** Number of cells analyzed and traces rejected using the TACO pipeline.

| # of cells collected | # of cell analysis failure | # of cells analyzed | # of traces analyzed | # of traces rejected for analysis (from # cells) |
| --- | --- | --- | --- | --- |
| 2735 | 5 | 2730 | 122,208 | 1990 (119) |
| Current trace amplitude $\geq 5000$ pA | Current trace mean value $\geq 5000$ mV | Voltage trace amplitude $\geq 190$ mV | Voltage trace mean value $\geq 10$ mV | Trace-based input resistance $\leq 0$ GΩ |
| 0 | 0 | 47 | 1947 | 6 |

For these cells, we then relied on the TACO pipeline to perform the I/O analysis, considering the spikes produced during the first 500ms of the response. During this analysis, several conditions must be met to ensure reliable estimate of the I/O properties of the neurons. In total, out of the 2730 cells, 2301 underwent the full analysis (Table 4).

**Table 4:** I/O fit results. Successful fit procedure are indicated by either Hill of Hill-Sigmoid. The other lines present all the possible reasons for the fit to fail, in which case the fit is not performed

| Obs | Allen Cell Type Database | NVC Database | Scala 2019 | Scala 2020 |
| --- | --- | --- | --- | --- |
| **Total** | **1398** | **207** | **330** | **795** |
| Less than 3 different frequencies to fit | 16 | 1 | 2 | - |
| Less than 4 different Non zero response | 64 | 5 | 1 | - |
| Minimum frequency step higher than 30Hz | 130 | 4 | 12 | 39 |
| 3rd order polynomial fit did not catch ascending portion | - | - | 21 | 131 |
| Both fitting methods (Hill and Hill sigmoid) stuck at initial conditions | 1 | - | - | - |
| Not enough traces to perform 3rd order polynomial fit | 1 | - | - | - |
| Stimulus not enough spread | 1 | - | - | - |
| Hill | 995 | 101 | 69 | 38 |
| Hill-Sigmoid | 190 | 96 | 225 | 587 |

### 3.2 Statistical analysis

The statistical analysis were performed in R programming language (v 4.3.1), using rstatix package. Before conducting the various statistical analyses to determine mean differences, we first tested the parametric hypotheses to determine the appropriate test. To assess whether the data within each group were normally distributed, we employed the Shapiro-Wilk test. Additionally, we used the Levene’s test to evaluate the homogeneity of the variance among the groups. The observations were independent of one another. If either of the two preliminary tests was statistically significant (*p <* 0.05), then the variance test applied was the non-parametric Kruskal-Wallis to assess if there was any statistically significant difference between means of the groups of interest. If so (i.e.: *p <* 0.05) then a group-wise comparison test was performed using the Dunn’s test, with Bonferroni adjusted p-values. In the cases where both Shapiro-Wilk and Levene’s tests were not statistically significant (i.e.: *p* ≥ 0.05), then a one-way ANOVA test was used to assess any statistically significant difference between means of the groups of interest. If so, then post-hoc Tukey test was used for pair-wise comparisons, with p-values corrected using Bonferroni methods. The p-values reported on the plots represent the following levels of statistical differences : Not significant : 1 ≥ *p >* 0.05, ∗ : 0.05 ≥ *p >* 0.01, ∗∗ : 0.01 ≥ *p >* 0.001, ∗ ∗ ∗ : 0.001 ≥ *p >* 0.0001, ∗ ∗ ∗∗ : *p* ≤ 0.0001. When the variance test (Kruskal-Wallis or ANOVA) was statistically significant (*p <* 0.05), the effect size (*η*^2^) was computed on pairs of levels by performing variance test on pairs of levels (when the factor of analysis had more than two levels).

To compare the shape of two distributions, we relied on the Kolmogorov-Smirnov (KS) test. This test compares the cumulative distributions of the two samples and relies on this maximum difference for test statistic. The statistical significance levels are as described previously. This test was performed using the kstest function from the scipy.stats library in Python.

Correlation between features was computed by using the Kendall’s tau correlation coefficient, to account for non-normal distributions. The correlation was considered as statistically significant or not based on the p-value of the correlation test (”stats” library, R). When significant, the correlation was considered as : negligible : |*τ* | ≤ 0.1; weak : |*τ* | ≤ 0.29; moderate : |*τ* | ≤ 0.59; strong : |*τ* | ≤ 0.89, very strong : |*τ* | ≥ 0.9 (Akoglu, 2018).

### 3.3 Distribution fit

The distribution of each feature was fitted using Maximum Likelihood Estimation, utilizing the “fit” method from the “scipy.stats” package in Python. For each population considered, the feature’s distribution was fitted to three different probability density functions: a Gaussian (Equation 1), a Skewed Gaussian (Equation 2) and a Log-Normal (Equation 3) :

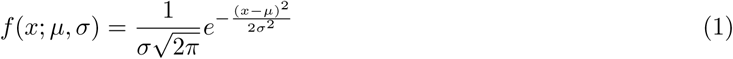

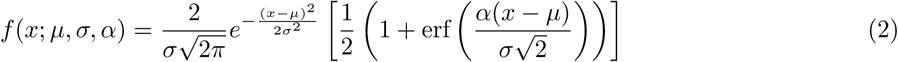

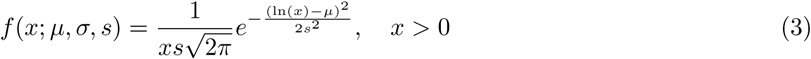

To determine the best-fitting distribution, a Kolmogorov-Smirnov test was performed to analyze the divergence between the experimental distribution and each of the tested probability density functions. The theoretical distribution that resulted in the highest p-value was considered the best fit to describe the experimental distribution.

## 4 Results

### 4.1 Analysis description

Our analysis aimed at characterizing the electrophysiological diversity of neuronal functional properties, obtained from comparable current-clamp protocols measuring the response to long duration (e.g. 500msec) current inputs. We analyzed subthreshold membrane properties of the neuronal membrane as the resting potential, the time constant and the input resistance measured at resting potential. We also analyzed the functional properties of cortical neurons describing the neuron’s I/O relationship, including gain, rheobase, in some cases, saturation frequency and spike frequency adaptation. These features were analyzed in different neuronal populations, according to either different aspects of the experimental protocol, or according to cellular or anatomical classification. The classification of neurons based on the type of neurotransmitter they release enables to decipher between excitatory and inhibitory neurons, while interneurons can be further described according to their molecular profile (i.e.: PValb, Sst, Vip, Htr3a). Such classification of cortical neurons, along with distinctions based on cortical areas and layers within those areas, offers the benefit of being non-overlapping, therefore accounting for the entire neuronal population of cerebral cortex. Table 5 summarizes the different comparisons and their associated datasets. For each figure, all descriptive statistics and best-distribution fitting parameters are presented in supplementary materials.

**Table 5:** Dataset Comparisons. Temp = Temperature. RT = Room temperature. PT = Physiological temperature (34°C for *in-vitro* protocols). Block = Synaptic blocker. Type = Transgenetic classification of cell types. State: Synaptic state (*in-vitro* and *in-vivo*). A = Section 4.2.1. B = Section 4.2.2. C = Section 4.3.1. D, E = 4.3.2. F = Section 4.4.1. G = Section 4.4.2. H = Section 4.5.1. I = Section 4.5.2.

| Comparison |  |  |  |  |  |  |  |  | Dataset |  |  |  |  |  |
| --- | --- | --- | --- | --- | --- | --- | --- | --- | --- | --- | --- | --- | --- | --- |
| State | Temp | Type |  |  | Area |  | Layer |  | Source | Area | Layer | Temp | Protocol | Block |
| A |  | C |  |  | F |  | H |  | Allen | V1 | 2/3, 4, 5, 6 | PT | In-vitro | Yes |
|  |  |  | D |  |  | G |  |  | Scala 2019 | V1 | 4, 5 | PT | In-vitro | No |
|  |  |  |  |  |  | G |  |  | Scala 2019 | SS | 4, 5 | PT | In-vitro | No |
|  | B |  |  | E |  |  |  | I | Scala 2020 | M1 | 2/3, 5, 6 | RT | In-vitro | No |
|  | B |  |  |  | F |  |  |  | Scala 2020 | M1 | 2/3, 5, 6 | PT | In-vitro | Yes |
| A |  |  |  |  |  |  |  |  | NVC | V1 | All | PT | In-vivo | No |

### 4.2 Impact of experimental conditions on electrophysiological properties

When addressing neuronal electrophysiological properties, each aspect of the protocol can play an important role on the quantitative measurements made during the recordings. All datasets considered here used whole-cell patch recordings to study biophysical properties of different neuronal populations in mouse cerebral cortex, yet each employed specific methodologies that could influence neuronal response to current inputs. The first distinction was whether the protocol was in-vitro (brain slice, Allen CTD, Scala 2019 and Scala 2020) or in-vivo (anesthetized, NVC). The major part of the Scala 2020 database was recorded at a decreased temperature (25°C, to extend recording times), while other databases recorded neurons at 34°C, thus near physiological temperature, as was the case for the NVC database. The final pertinent distinction between protocols was the pharmacological alteration of cellular properties: the Allen CTD, as well as part of the Scala 2020 database used kynurenic acid and picrotoxin in their recording solutions, to block fast glutamatergic and GABAergic synaptic transmissions, while the Scala 2019 and NVC databases did not. To better understand how these experimental parameters could influence cellular properties, we compared neuronal electrophysiological properties across databases differing in experimental protocols.

#### 4.2.1 Synaptic State: *In-vitro* vs. *in-vivo*

One consequence of different experimental protocols is the level of spontaneous synaptic activity impinging on a recorded cell, with a subsequent impact on the I/O relation. Spontaneous firing is usually very low for *in vitro* protocols, nevertheless spontaneous synaptic events are typical, and may be explicitly controlled by pharmacolgical synaptic blockers. Spontaneous firing during *in vivo* protocols depends on the level of anesthesia, but for standard protocols are typically between 1 and 10Hz (Destexhe et al., 2003). If we consider that there are typically thousands of afferent inputs to a single cortical neuron, the effective rate of spontaneous activity to a cell can easily be several kHz. To address this question we compared recordings from visual cortex taken from the Allen CTD (*in vitro*) and NVC (*in vivo*) databases. For simplicity we considered only regular spiking cells, thus all cell types except PValb neurons from the Allen CTD, and all cells from NVC database whose maximum firing frequency was lower than 100Hz.

Corresponding with the hypothesis of increased spontaneous synaptic activity *in vivo* (Destexhe et al., 2003), the input resistance for cells recorded *in vivo* is significantly lower compared to cells recorded *in vitro* (Figure 3A). Furthermore, cells recorded *in vivo* have lower gains with respect to the input current (Figure 3B) and higher rheobases (Figure 3C), as compared to *in vitro* cells. These differences are confirmed by a Kruskal-Wallis test on the distributions (Figure 3B and C, middle) and by comparing the CDFs with a Kolmogorov-Smirnov test (Figure 3B and C, bottom).

To explain these difference in I/O properties between the two conditions, we considered that since biophysical properties at the cellular level arise from various voltage-dependent ion channels particular for each type, the voltage induced by a given current input is a more relevant input variable for comparing I/O characteristics across cells. Indeed, we found significant correlations between input resistance and both rheobase and current-based gain in both datasets (Figure 2). Correspondingly, this suggests that a higher background synaptic activity *in vivo* can account for many differences in functional properties as compared to *in vitro* neurons (Destexhe et al., 2003).

**Figure 2:**
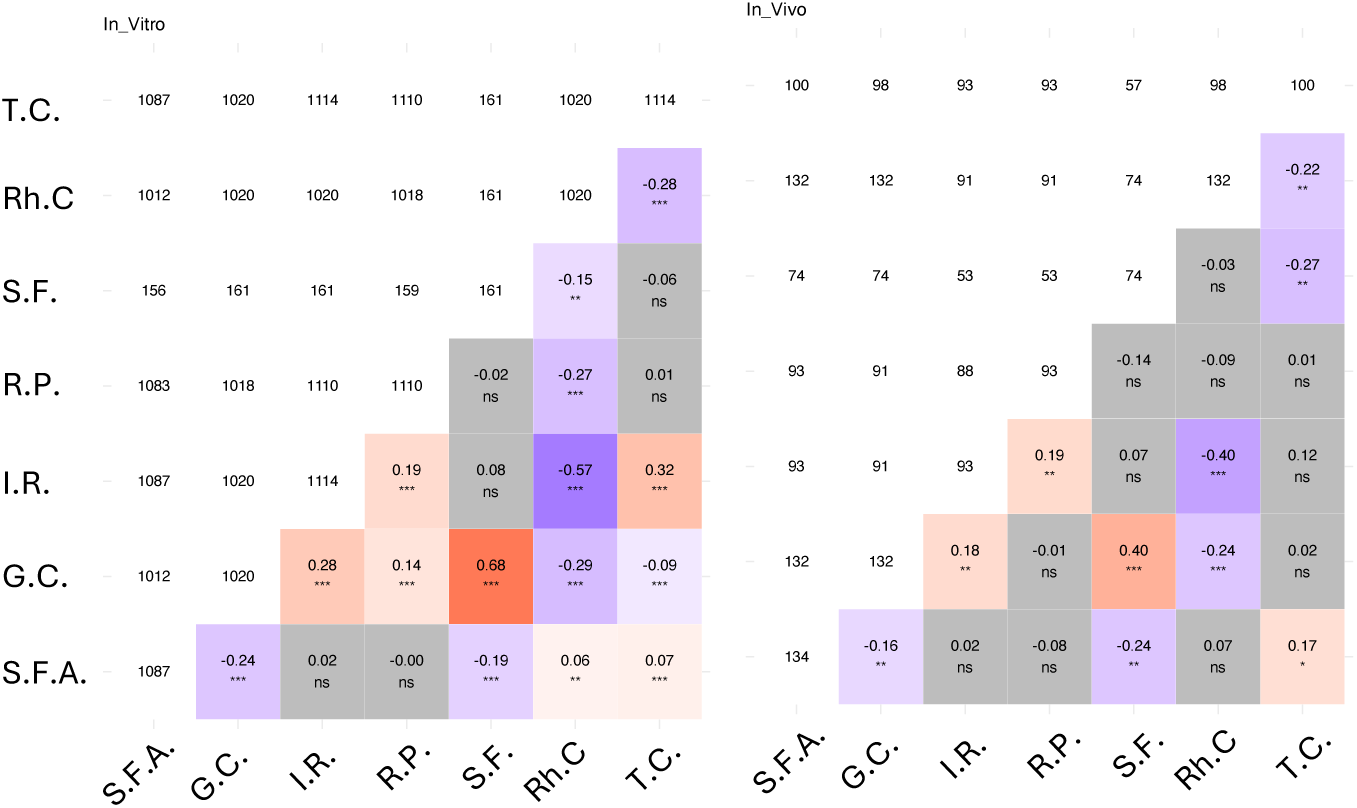
Correlation of linear and current-based firing properties in cells recorded in *in vitro* or *in vivo* experiments. For each matrix, the diagonal represent the number of observation per features, the upper triangular matrix represents the number of joint observation between two features, and the lower triangular matrix represents the Kendall’s tau correlation coefficient, with the statistical significance. Only correlation analysis which were statistically significant (p≤0.05) were colored according to the correlation sign. Cells used for this analysis were recorded at physiological temperature in mice’s visual cortex during either *in vitro* (Allen CTD) or *in vivo* (NVC) recordings

To test this hypothesis we rescaled the current input for a given recording by the input resistance of the cell, obtaining the estimated voltage response of a linear cell model. This procedure translates the rheobase input (pA) to an input voltage threshold input (mV) (Figure 3E), and the current-based gain (Hz/pA) to a voltage-based gain (Hz/mV) (Figure 3C). This transformation of both properties resulted in statistically non-distinguishable *in vivo* and *in vitro* populations, as suggested by the non statistically significant Kruskal-Wallis tests (Figure 3E and C, middle), and non significant Kolmogorov-Smirnov tests on the CDFs (Figure 3E and C, bottom).

**Figure 3:**
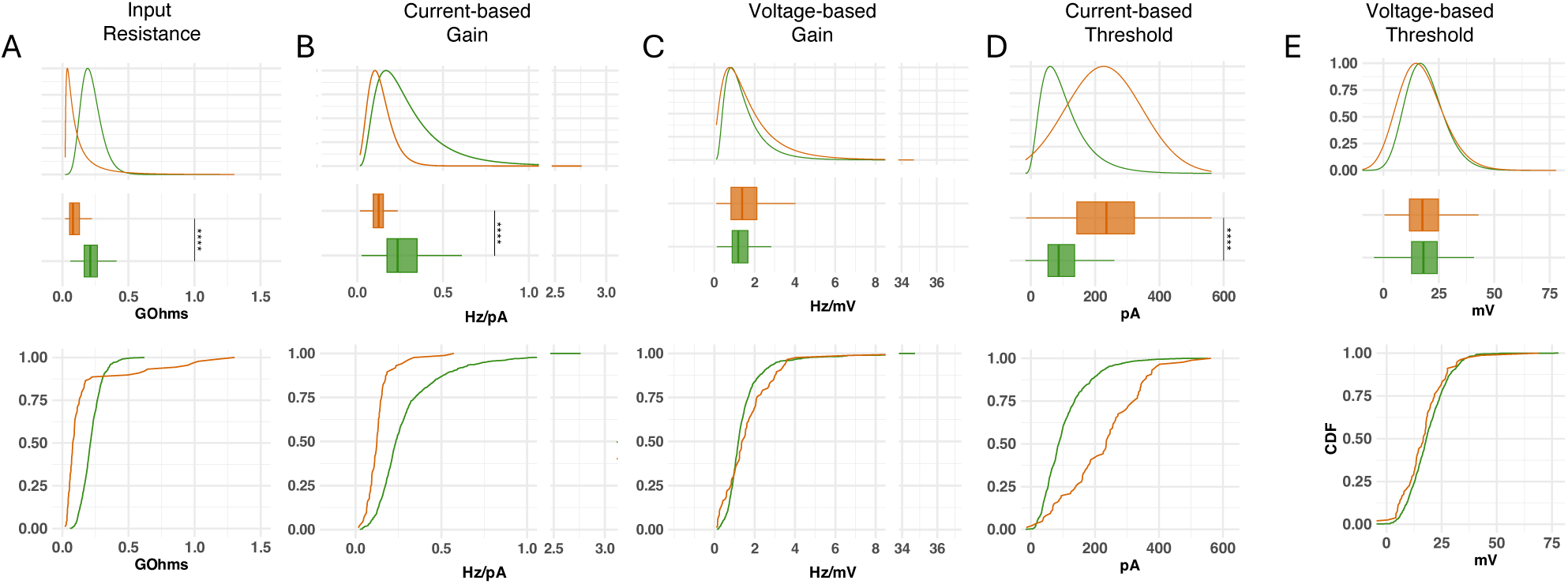
Differences of electrophysiological properties of visual cortex cells *in vivo* and *in vitro*. Cells from the Allen CTD and NVC databases. **A** Input resistance of cells recorded *in vivo* is significantly reduced compared to *in vitro* cells. Firing properties such as current-based gain (**B**) and rheobase (**D**) also show significantly different distributions, which were markedly reduced when adjusted for the input resistance according to a linear circuit model, thus for the gain expressed in Hz/mV (**C**) or for the threshold expressed in mV (**E**). Top: Fitted distributions. Middle: Box plots with significant differences. Bottom: CDFs from the data. Summary statistics and distribution fit parameters are presented in Supplementary Table 6

Though a good match for the main ionic consitutents is to be expected, undoubtably the extracellular millieu *in vivo* is much more complex than the artificial bath solution of the *in vitro* preparation. Nevertheless, these results support the hypothesis that differences between cells in I/O measures based on input current recorded *in vivo* or *in vitro* can be partly explained by higher network activity in the former case lowering the cell’s input resistance. Finally, the overall correlation between the current-based I/O parameters and input resistance motivated considering the these measures in terms of the linear voltage input for the subsequent analyses comparing different *in vitro* datasets.

#### 4.2.2 Recording temperature

To examine the impact the temperature on *in vitro* recordings, we compared linear and firing properties of L5 Sst neurons recorded in motor cortex from the Scala 2021 database, recorded at 25°C or 34°C. The lower temperature gave a higher input resistance (Figure 4A) and, accordingly, a higher membrane time constant (Figure 4B), as well as a more depolarized resting potential (Figure 4C). The differences for all three parameters, while statistically significant, were relatively small. As described above, an increased input resistance would normally be associated with an increase in the response to current input, and indeed a lower temperature resulted in lower rheobase (Figure 4E). Paradoxically, though, a lower temperature reduced current gain (4D). Considering firing parameters adjusted for voltage-based input, a lower temperature reduced voltage-based threshold (Figure 4H) and voltage-based input gain (Figure 4G), but did not align the two conditions as described in the previous section. Interestingly, no statistically significant difference was observed when comparing the voltage threshold relative to the resting potential (Figure 4I). Decreased temperature also impacted other properties of the spike train evoked by long current step, thus reducing the saturation frequency (Figure 4F), and accelerating spike frequency adaptation (Figure 4J). Globally, these diverse and complex effects on the general I/O properties complicate a comparative analysis of datasets obtained at different temperatures, and consequently the remainder of the comparative analyses are constrained to the same recording temperature.

**Figure 4:**
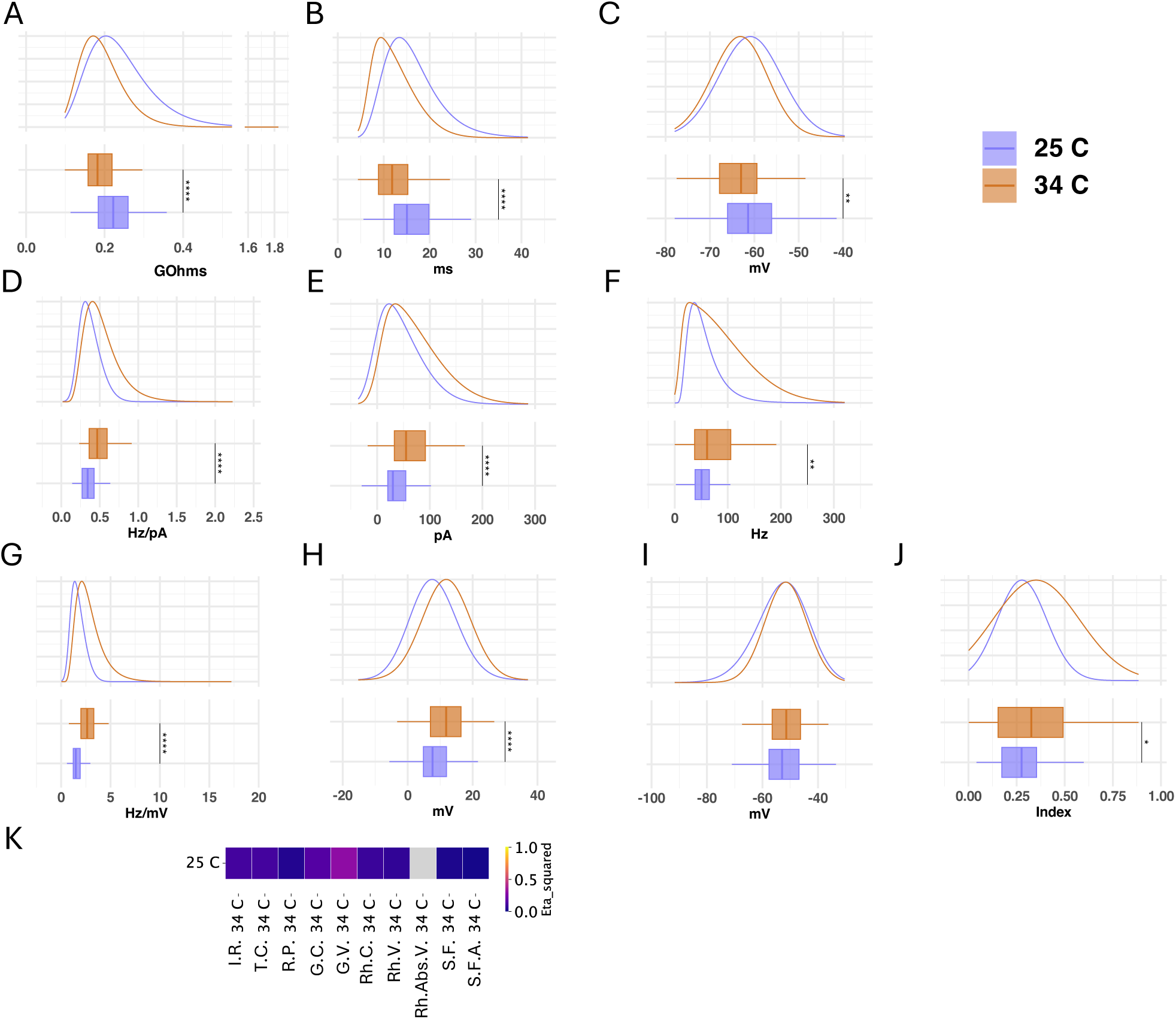
Influence of recording temperature on electrophysiological properties in Sst neurons from the layer 5 of Motor cortex. Relying on cells from the Scala 2021 database, we compared between cells recorded at either 25°C (136 cells) or 34°C (108 cells)**A**) Input Resistance (I.R.), **B**) Time constant (T.C.), **C**) Resting potential (R.P.), **D**) Gain expressed in current-based input (G.C.), **E**) Rheobase expressed in current-based input (Rh.C.) **F**) Saturation frequency (S.F.), **G**) Gain expressed in voltage-based input (G.V.), **H**) Rheobase expressed in voltage-based input (Rh.V.), **I**) Rheobase expressed in absolute voltage-based input (Rh.Abs.V.) and **J**) Spike Frequency Adaptation (S.F.A.). **K**: Heat Map representing the effect size of variance test for the different features. Features for which the difference was non-statistically significant are in grey. Summary statistics and distribution fit parameters are presented in Supplementary Table 7

### 4.3 Heterogeneity of electrophysiological properties between cell types

#### 4.3.1 Visual cortex with synaptic block

We first considered the cell types of visual cortex from the Allen database, recorded at physiological temperature with synaptic blockers.

Cortical neurons were found to display large cell type related heterogeneity in their linear properties. For example, PValb expressing interneurons recorded in Visual cortex at 34°C (Allen CTD) had statistically significant lower input resistance compared to other cell types (mean 129*M* Ω) as revealed by a KW test (Figure 5 A). The input resistance also differ between other cell types (mean 202, 218, 247, 248 *M* Ω for Excitatory, Htr3a, Sst, and Vip cells respectively), even though the effect size corresponding to these neuron groups are lower than with PValb cells (up to 0.0862 between Excitatory and Vip cells) (Figure 5 J). To compare the differences between the distribution, we followed our analysis by performing pairwise Kolmogorov-Smirnov tests. Except for the distributions of Sst and Vip cells (p=0.073), all other pairs of distribution significantly differ (Sst-Htr3a p≤0.01; Excitatory-Htr3a ≤0.001, all other combinations had p≤0.0001). This analysis therefore confirmed that based on input resistance, cortical neurons highly differ between classically defined classes. PValb neurons have the lowest input resistance, followed by excitatory and then by Htr3a, Sst and Vip cells. However, the effect size were limited between these categories, suggesting the input resistance between these cell types only slightly differ.

**Figure 5:**
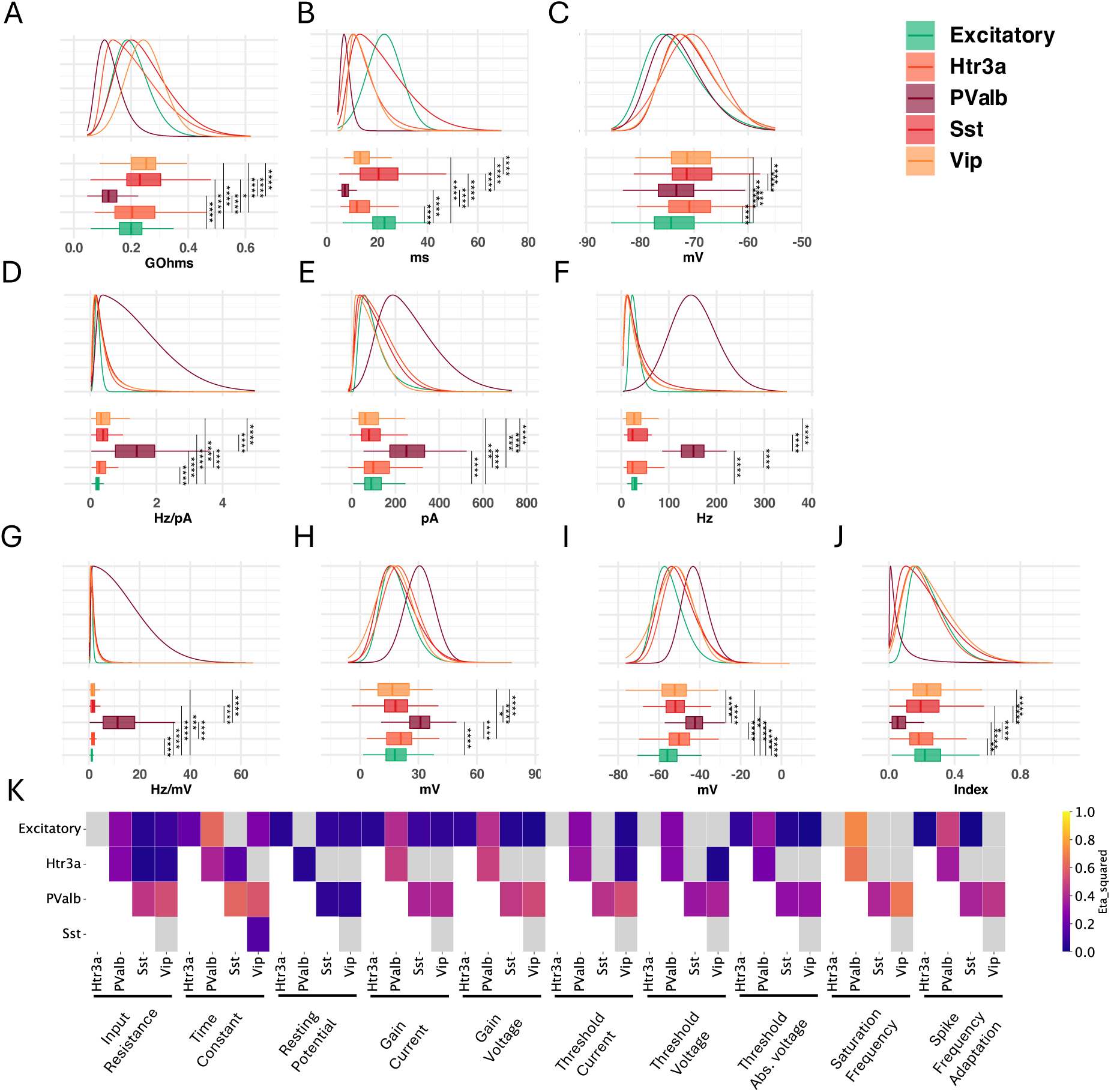
Heterogeneity across cell types: Visual cortex (Allen CTD). Comparison of **A**: Input resistance, **B**: Time constant, **c**: Resting potential, **D**: Current-based gain, **E**: Current-based threshold, **F**: Saturation frequency, **G**: Voltage-based gain, **H**: Voltage-based threshold relative to *V_rest_*and **I**: Voltage-based threshold, in different neuronal types. **J**: Heat map of effect size from KW test performed on features pairs. Only KW test which were statistically significant are colored, grey tiles represent non statistically significant differences. The analysis highlights the increased excitability of PValb neurons compared to other neuronal types. Summary statistics and distribution fit parameters are presented in Supplementary Table 8

Similar observation could be made regarding the differences in the membrane time constant. PValb cells had the lowest time constants (mean 7.54 msec), followed by Htr3a and Vip cells (14.5ms for both cell types). Finally, Sst (22.1 msec) and Excitatory (22.8 msec) cells had the longest time constant (Figure 5 B). For all pairs of cell types, the variance analysis revealed statistically significant differences (except between Vip and Htr3a, p = 1, and between Excitatory and Sst cells, p = 0.07), with large effect size (*η*^2^ between 0.148 between Sst and Vip cells, up to 0.617 between PValb and excitatory cells) (Figure 5 J). This observation was then confirmed by the KS test, which reported that for all pairs of cell types, the differences of distribution were statistically different, even between Sst and Vip cells, and between Vip and Htr3a. For the later pair, we had p≤0.01, while for all other pairs, p≤0.0001. The differences of results between the KW and Dunn test, and the KS test can be explained by the fact that the two test do not measure the same aspects of the data. While KW test measure the central tendency between different populations, KS test relies on the underlying distributions shape. Therefore two distributions may have similar mean values, but differ in their distribution, resulting in apparently different results.

The distributions of the cell resting potential *V_rest_* also differed between neuronal types. Specifically, excitatory and PValb neurons were observed to be more hyperpolarized (mean -73.5 mV and -73.0 mV, respectively) compared to Htr3a (-70.6 mV), Vip (-70.3 mV), and Sst (-70.3 mV). Statistical differences were identified across most pairs, except between excitatory and PValb cells, and among Htr3a, Vip, and Sst cells (Figure 5 C). Correspondingly, effect sizes were minimal between excitatory and PValb cells, and among Htr3a, Vip, and Sst cells. These findings aligned with results from KS tests, indicating that the resting potential of excitatory and PValb cells differed from those of Sst, Htr3a, and Vip cells.

Our analysis continued with the comparison of functional properties, emphasizing input threshold and firing gain. The largest differences in firing properties were observed between PValb cells and other neuron types. Indeed, PValb cells had higher gain, threshold and saturation frequency values compared to other cell types. We notably observed in the cells recorded in the Allen CTD that the main differences were observed between PValb cells and other cell types, and between excitatory and other cell types (Figure 5 D). Indeed, excitatory cells had the lowest gain among these neuron categories (0.22 Hz/pA) while PValb had the highest values (1.68 Hz/pA). In between, Htr3a (0.35 Hz/pA), Vip (0.42 Hz/pA) and Sst (0.45 Hz/pA) cells’ mean gain values did not statistically differed (Dunn test adjusted p = 1). These results were confirmed by similar results obtained from KS test, except between Sst and Htr3a neurons (p ≤ 0.01), suggesting that mainly excitatory and PValb cells’ gain distribution differed from the others. Furthermore, expressing gain in term of linear voltage (as explained earlier) (Figure 5 G) yielded similar results with both KW/Dunn test and KS tests, except between Sst and Htr3a neurons for which the KS test was not statistically significant anymore (p = 0.24).

Similarly, PValb cells had the higher rheobase among the different neuron types (mean 271.7 pA), followed by Htr3a, excitatory, Sst and Vip neurons (124.2, 107.5, 103.9 and 82.7 pA respectively) (Figure 5E). Statistically, PValb cells differed from all other cell types, as did Vip cells (except when compared to Sst cells). Excitatory, Htr3a and Sst cells did not significantly differ between each other. On the contrary, based on the distribution shapes, there were statistically significant differences between all pairs neuron types (except between excitatory and Htr3a cells, KS p = 0.059). Re-scaling the rheobase to the input resistance of the cell (therefore expressing the input threshold in mV), slightly altered the level of statistical differences, (Figure 5H), but yielded similar effect size (Figure 5J). Therefore, no difference was detected between Vip and excitatory cells anymore(KW test, p=1). Also, based on the distribution shapes, all pair of neurons types statistically differed, except Excitatory and Sst cells (KS test, p=0.31) and Sst and Htr3a (KS, p=0.13). Interestingly, reporting the voltage-based threshold to increased the differences between the neuronal types, as PValb neurons and Excitatory cells were both statistically different from all other cell types (Figure 5 I). The analyses of both the voltage-based threshold and absolute voltage-based threshold suggest that PValb neurons require a proportionally stronger input relative to their resting membrane potential to elicit neuronal firing.

Finally, the saturation frequency varied among cell types (Figure 5 F). PValb cells had the highest saturation frequency (mean 158.8 Hz), followed by Sst (50.8 Hz), Htr3a (33.2 Hz), Vip (33.1 Hz), and excitatory cells (28.3 Hz). The KW/Dunn test revealed that only PValb cells differed from other cell types, which was further confirmed by the KS test, showing that PValb cells differed from all other neuron categories (p≤0.0001). Additionally, the distribution shape of Vip and excitatory cells was different (p≤0.05). This distinct faster firing for PValb cells supported the previous segregation of these cells from the other, assumed regular spiking, cell types in the comparison between the *in vitro* and *in vivo* datasets.

#### 4.3.2 Visual cortex and motor cortex with no synaptic block

These observations based on the Allen CTD were coherent with an analyis on other datasets recorded without synaptic blockers, notably for motor cortex from Scala 2020 (Figure 6 and for visual cortex from Scala 2019 (Figure 7)). Quantitative differences between the datasets, particularly for the visual cortical neurons, may be attributed to synaptic blockers, as well as different prootocol temperatures (Allen and Scala 2019, physiological temperature; Scala 2020, room temperature). Nevertheless the qualitative similarities between the datasets support the hypothesis that the classification of cortical interneurons according to their molecular categories has a correspondence to their electrophysiological profile.

**Figure 6:**
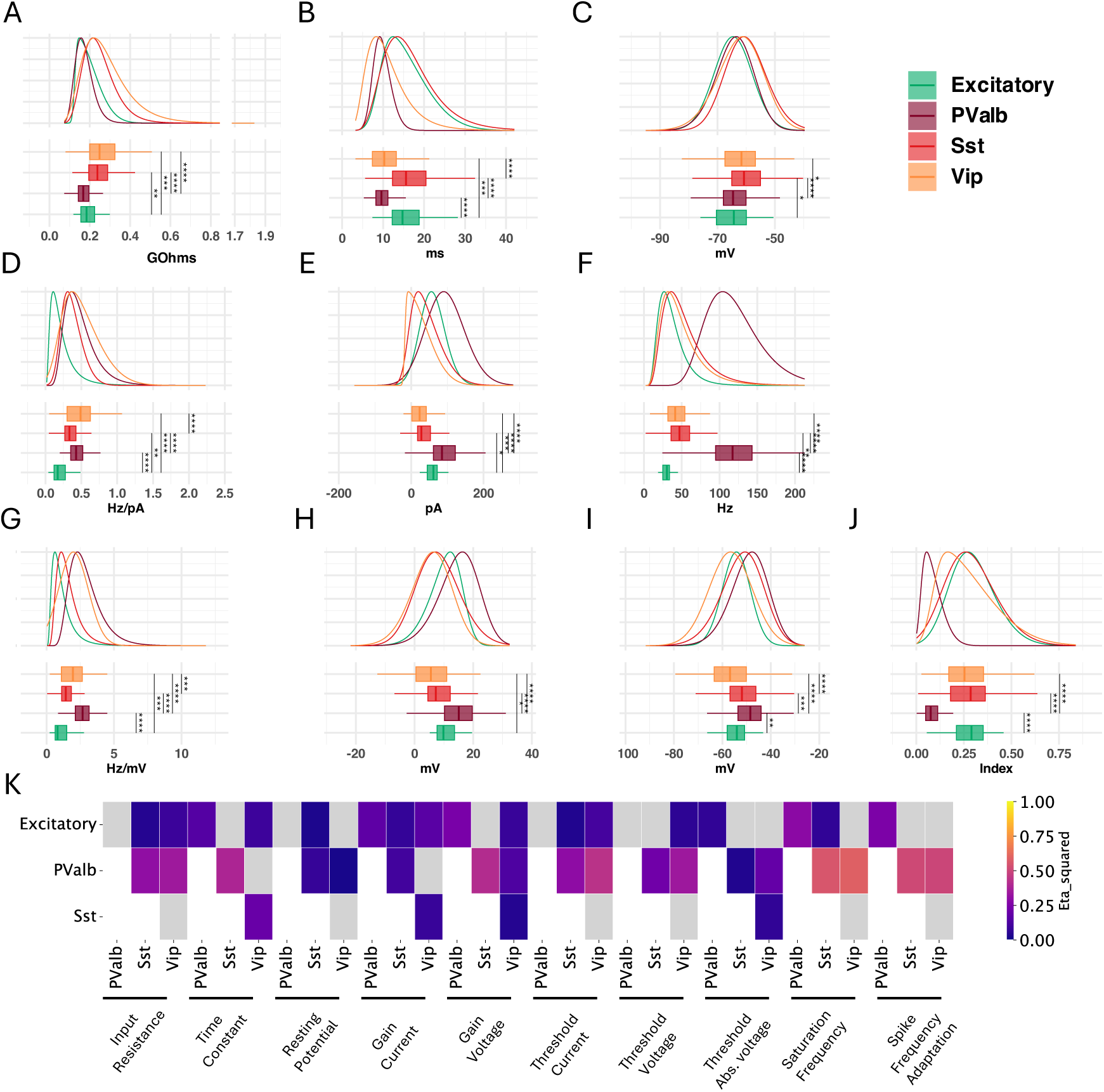
Heterogeneity across cell types: Motor cortex (Scala 2020). Comparison of **A**: Input resistance, **B**: Time constant, **c**: Resting potential, **D**: Current-based gain, **E**: Current-based threshold, **F**: Saturation frequency, **G**: Voltage-based gain, **H**: Voltage-based threshold relative to *V_rest_*and **I**: Voltagebased threshold, in different neuronal types. **J**: Heat map of effect size from KW test performed on features pairs. Only KW test which were statistically significant are colored, grey tiles represent non statistically significant differences. The analysis, made on cells recorded at 25°C, without the use of synaptic blockers (Scala 2020 database) highlights the increased excitability of PValb neurons compared to other neuronal types. Summary statistics and distribution fit parameters are presented in Supplementary Table 9

**Figure 7:**
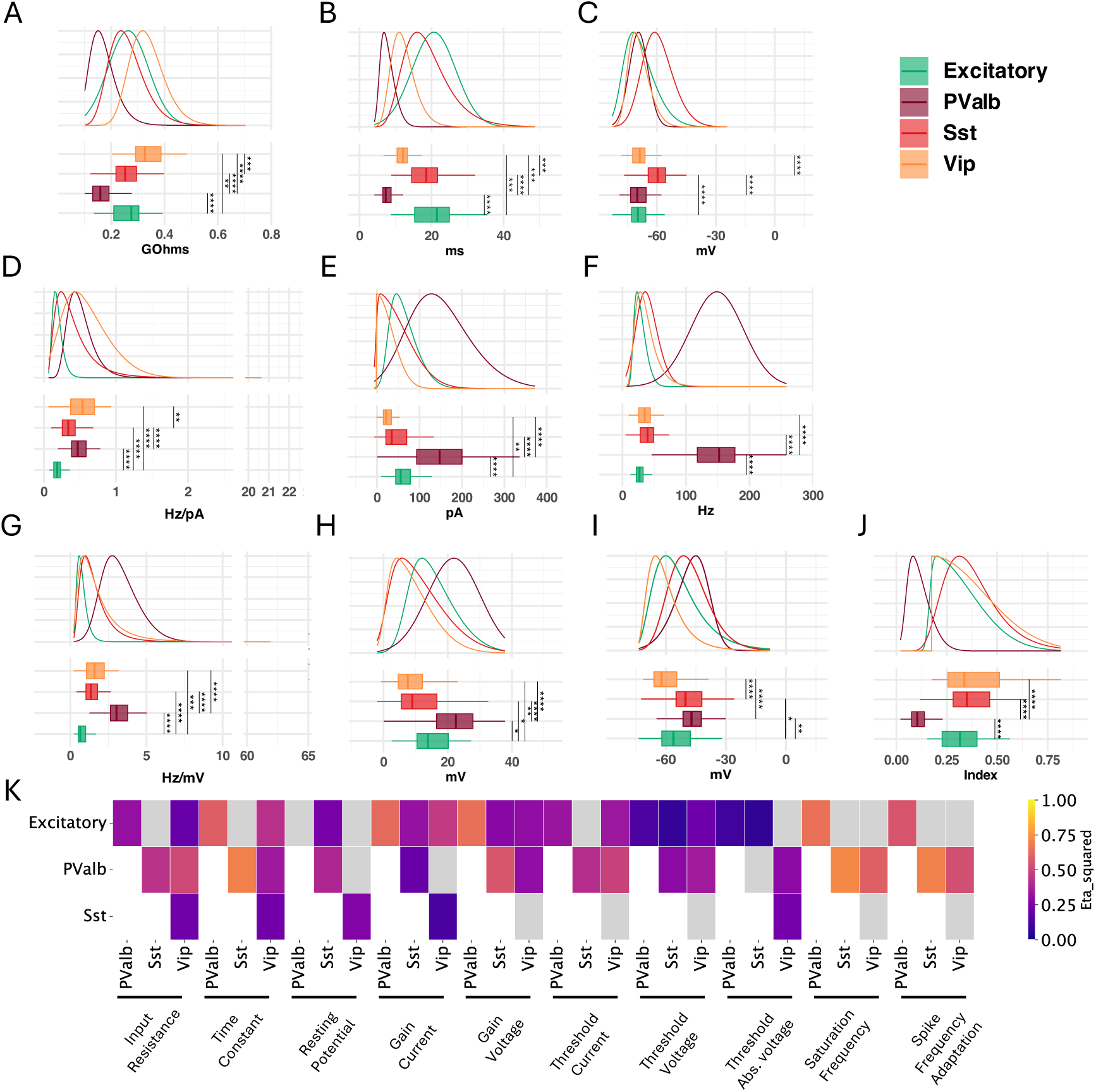
Heterogeneity across cell types: Visual cortex (Scala 2019). Comparison of **A**: Input resistance, **B**: Time constant, **c**: Resting potential, **D**: Current-based gain, **E**: Current-based threshold, **F**: Saturation frequency, **G**: Voltage-based gain, **H**: Voltage-based threshold relative to *V_rest_*, **I**: Voltage-based threshold and **J** Spike Frequency Adaptation index, in different neuronal types. **K**: Heat map of effect size from KW test performed on features pairs. Only KW test which were statistically significant are colored, grey tiles represent non statistically significant differences. The analysis, made on cells recorded at physiological temperature, without the use of synaptic blockers (Scala 2019 database) highlights the increased excitability of PValb neurons compared to other neuronal types. Summary statistics and distribution fit parameters are presented in Supplementary Table 10

#### 4.3.3 Cell types have similar correlation relationships

We followed up by analyzing the correlation of electrophysiological properties among different cell types. For visual cortex we observed similar relationships between features across the different cell types (Figure 8), such as input resistance being strongly anti-correlated with current-based threshold, strongly correlated with time constant or correlation between gain and saturation frequency or strongly correlation between input resistance and time constant. These examples illustrated the typical relationships between linear and I/O properties describing neuronal excitability. Interestingly, PValb neurons showed weakly anti-correlated input resistance and current-based gain whereas other cell types had weakly correlation between these features. Similarly, PValb neurons differed from other neuronal types by having moderately correlated current-based gain and thresholds, whereas these properties in other cell types were moderately anti-correlated. This noticeable differences of PValb cells seem to correspond to steeper I/O relationships (i.e.: higher thresholds, and higher gains), potentially evoking Type II like excitability.

**Figure 8:**
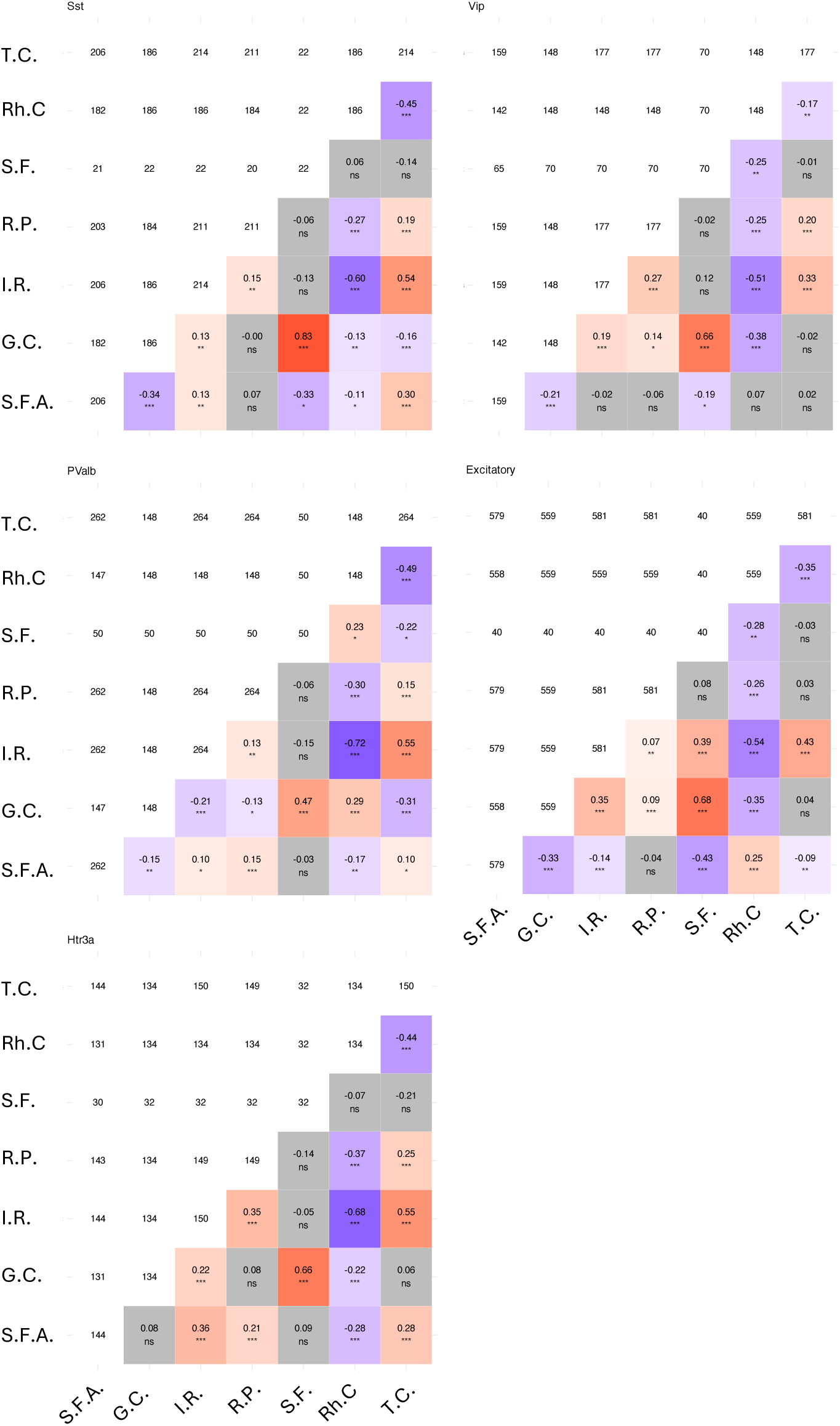
Correlation of linear and current-based firing properties in cells recorded in visual cortex from Allen CTD. For each matrix, the diagonal represent the number of observation per features, the upper triangular matrix represents the number of joint observation between two features, and the lower triangular matrix represents the Kendall’s tau correlation coefficient, with the statistical significance. Only correlation analysis which were statistically significant (p 0.05) were colored according to the correlation sign. Cells used for this analysis were recorded at physiological temperature with synaptic blockers (Allen CTD).

Interestingly, this specificity of PValb neurons was not observed in cells recorded in visual cortex without synaptic blockers (Figure 9) or in motor cortex recorded at 25°C with synaptic blockers (Figure 10). Overall, these cells displayed more cell-type related homogeneity and moderate correlation and anti-correlation between features compared to cells recorded in the visual cortex, the main differences being that excitatory cells displayed strong correlation of input resistance and saturation frequency, whereas in Sst and PValb neurons this relationship was moderately anti-correlated.

**Figure 9:**
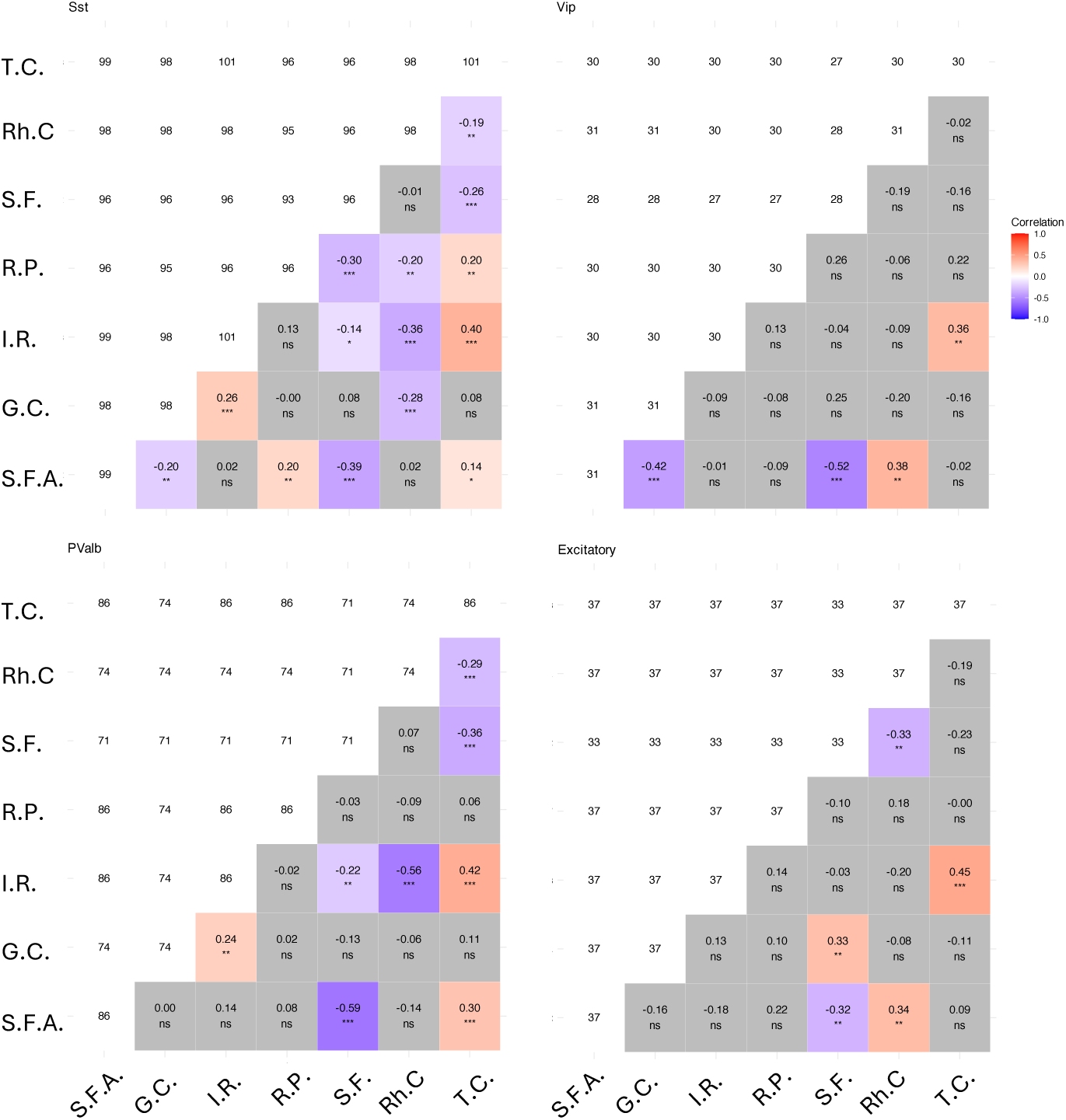
Correlation of linear and current-based firing properties in cells recorded in visual cortex from Scala 2019 Database. For each matrix, the diagonal represent the number of observation per features, the upper triangular matrix represents the number of joint observation between two features, and the lower triangular matrix represents the Kendall’s tau correlation coefficient, with the statistical significance. Only correlation analysis which were statistically significant (p 0.05) were colored according to the correlation sign. Cells used for this analysis were recorded at physiological temperature without synaptic blockers (Scala 2019 Database).

**Figure 10:**
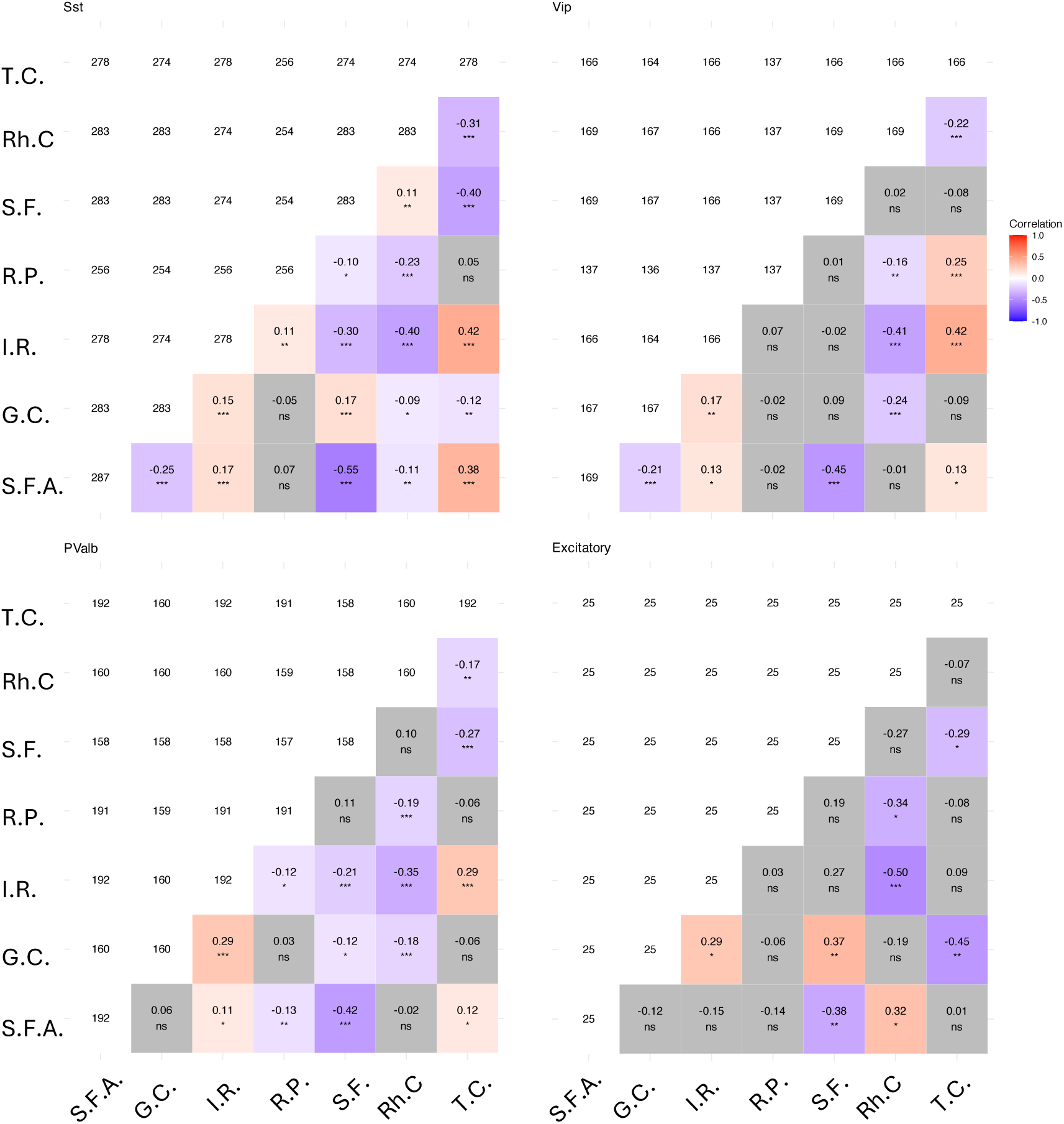
Correlation of linear and current)based firing properties in cells recorded in motor cortex. For each matrix, the diagonal represent the number of observation per features, the upper triangular matrix represents the number of joint observation between two features, and the lower triangular matrix represents the Kendall’s tau correlation coefficient, with the statistical significance. Only correlation analysis which were statistically significant (p 0.05) were colored according to the correlation sign. Cells used for this analysis were recorded at room temperature without synaptic blockers (Scala 2021).

These differences of correlation analysis between visual and motor cortex cells illustrates the fact that similar cell types can have different properties, depending on the cortical area these are recorded from, although we should not exclude the impact of synaptic blocker and temperature on neuronal properties. A more in depth investigation of neuronal variability according to anatomical considerations could help better characterize neuronal electrophysiological heterogeneity.

### 4.4 Cell type specific heterogeneity across cortical areas

#### 4.4.1 Sst L5 cells in visual and motor cortex

We next studied if the electrophysiological properties of cortical neurons varied according to their cortical area. To do so, we compared Sst cells recorded in layer 5 of visual and motor cortex at 34°C, using synaptic blockers (Visual cortex : Allen CTD, Motor cortex : Scala 2021) (Figure 11). These cells appeared to primarily differ according to their linear properties (KW test p≤0.0001 for input resistance, time constant and resting potential). Specifically, visual Sst cells had higher input resistances (mean 260 *M* Ω, 194 *M* Ω respectively, *η*^2^=0.168) and time constants than motor cortex Sst neurons (24.1 ms, 12.4 ms, *η*^2^=0.400), and were also more hyperpolarized (-70.5 mV, -63.5 mV *η*^2^ = 0.263). Similarly, firing properties varied according to the cortical area (KW, p≤0.0001, p≤0.05, p≤0.01 for gain, threshold and saturation). However, these differences were more moderate compared to linear properties, based on the effect size. Sst cells recorded in visual cortex had lower gain than motor cortex cells (mean : 0.39 Hz/pA, 0.53 Hz/pA respectively, *η*^2^=0.095), higher threshold (89.4 pA, 68.03 pA, *η*^2^=0.025) and lower saturation (37.8 Hz, 81.6 Hz, *η*^2^ = 0.056). Expressing gain and threshold in linear voltage input also reflected the decreased excitability in firing properties of visual Sst cells compared to motor cortex cells, with lower gain (mean : 1.69 Hz/mV, 2.92 Hz/mV, *η*^2^ = 0.1400), and higher threshold (mean : 19.2 mV, 6.7 mV, *η*^2^ = 0.283). However, by reporting the voltage-based threshold to the cells’ resting potential, no difference was observed between visual and motor cortex Sst cells (KW test, p=0.355, mean -51.8 mV, -56.6mV).

**Figure 11:**
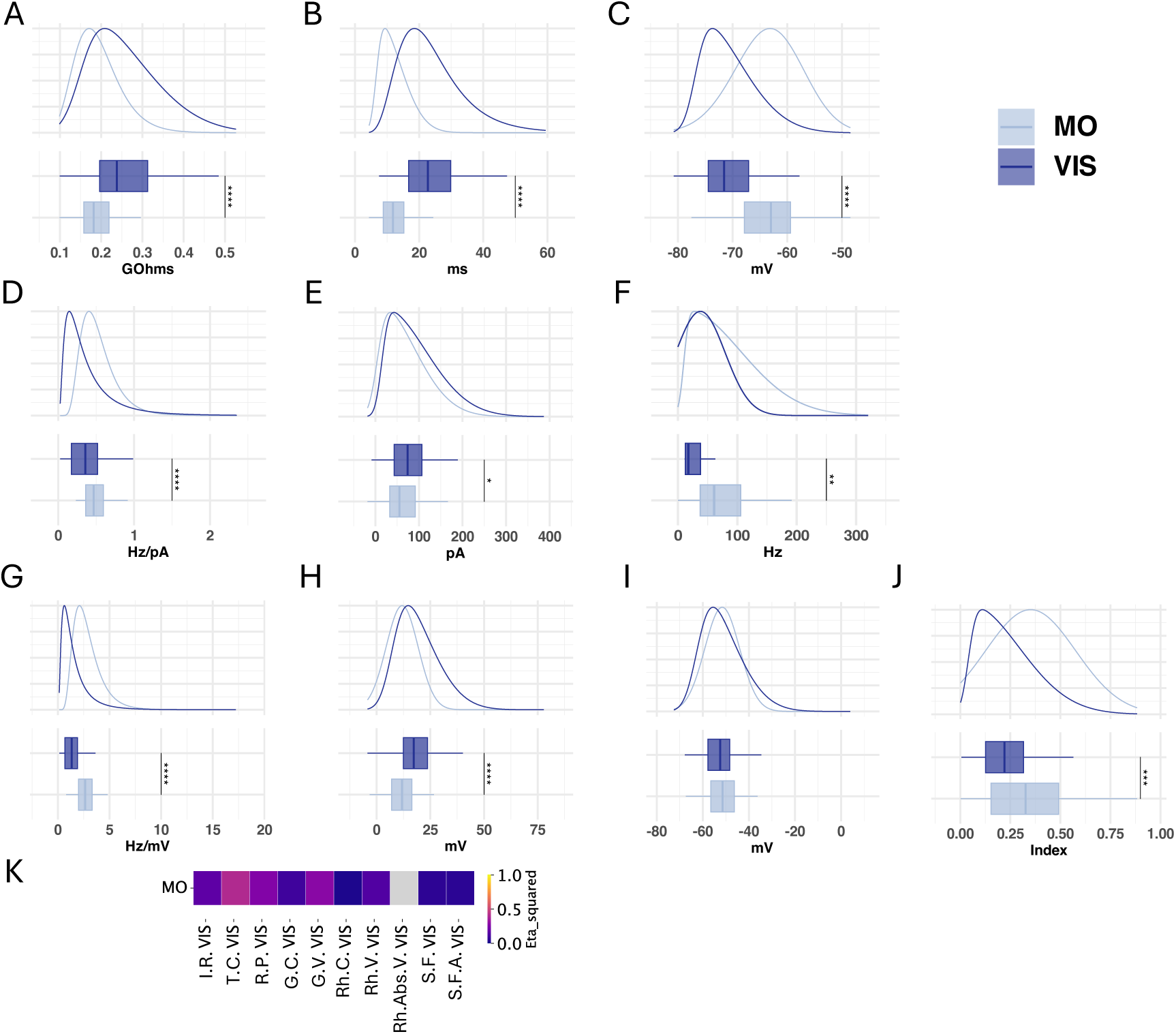
Comparison of electrophysiological properties in Layer 5 Sst cells recorded in visual or motor cortex.. Comparison of **A**: Input resistance, **B**: Time constant, **c**: Resting potential, **D**: Current-based gain, **E**: Current-based threshold, **F**: Saturation frequency, **G**: Voltage-based gain, **H**: Voltage-based threshold, **I**: Voltage-based threshold(ff) and **J** Spike Frequency Adaptation index, in different neuronal types. **K**: Heat map of effect size from KW test performed on features pairs. Only KW test which were statistically significant are colored, grey tiles represent non statistically significant differences. The analysis, made on cells recorded at physiological temperature, with the use of synaptic blockers, from Visual (Allen Cell Type Database) and Motor (Scala 2021 database) cortex. Summary statistics and distribution fit parameters are presented in Supplementary Table 11

#### 4.4.2 Sst L4 cells in visual and somatosensory cortex

To address the question of whether these intercortical area differences in linear and firing properties were observed only between visual and motor cortex neurons, we next compared the properties of Sst recorded in layer 4 of either visual or somatosensory cortex, without the use of synaptic blockers (Scala 2019 database) (Figure 12). Similarly to the comparison between visual and motor cortex, Sst cells recorded in visual cortex appeared to be less excitable compared to somatosensory Sst cells. Indeed, visual Sst cells had higher input resistance compared to cells recorded in somatosensory cortex (mean 266 *M* Ω, 190*M* Ω, respectively, *η*^2^ = 0.321), and time constant (19.3ms, 11.1ms, *η*^2^ = 0.431). Also, visual Sst cells had lower saturation frequency (39 Hz, 79 Hz, *η*^2^ = 0.472), and higher index of spike frequency adaptation (0.40, 0.27,*η*^2^ = 0.251). However, Sst cells recorded in either visual or somatosensory cortex did not differed in resting potential (KW test, p = 0.338, mean : -59.7 mV, -59.2 mV), nor in gain (KW test: p = 0.078, median 0.32 Hz/pA, 0.36 Hz/pA) and rheobase (KW test, p = 0.731, mean : 52.3 pA, 53.3 pA). Yet, when expressing gain (median 1.27 Hz/mV, 1.86 Hz/mV, *η*^2^ = 0.308) and rheobase (mean 12.01 mV, 8.69 mV, *η*^2^ = 0.037) in voltage input, the difference between Sst cells recorded in visual and somatosensory cortex was statistically significant, suggesting a reduced excitability for Sst cells recorded in Visual cortex compared to cells recorded in somatosensory cortex. However, no difference was observed when expressing threshold in absolute voltage input (KW test, p = 0.121, mean: -47.6 mV, -50.5 mV).

**Figure 12:**
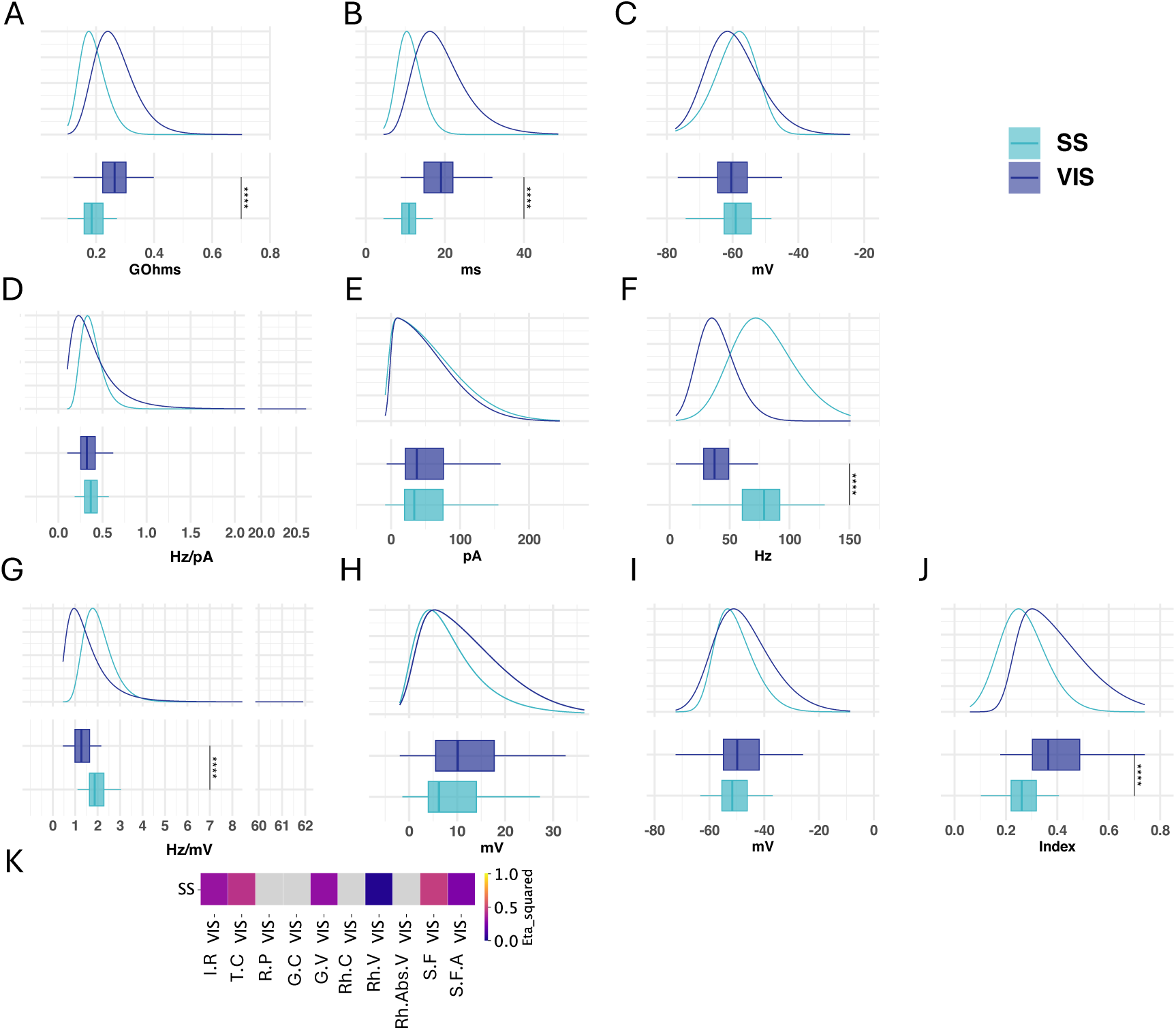
Comparison of electrophysiological properties in Layer 4 Sst cells recorded in visual or somatosensory cortex.. Comparison of **A**: Input resistance, **B**: Time constant, **c**: Resting potential, **D**: Current-based gain, **E**: Current-based threshold, **F**: Saturation frequency, **G**: Voltage-based gain, **H**: Voltage-based threshold, **I**: Voltage-based threshold(ff) and **J** Spike Frequency Adaptation index, in different neuronal types. **K**: Heat map of effect size from KW test performed on features pairs. Only KW test which were statistically significant are colored, grey tiles represent non statistically significant differences. The analysis, made on cells recorded at physiological temperature, without the use of synaptic blockers, from Visual and Somatosensory cortex (Scala 2019 database). Summary statistics and distribution fit parameters are presented in Supplementary Table 12

These results suggest that electrophysiological properties of cortical interneurons actually vary according to the cortical areas they originate from. Notably, the differences of input resistance in these different situations suggest that visual Sst cells have smaller cell bodies compared to somatosensory and motor cortex cells. Yet, the fact that firing features expressed in linear voltage input, as well as time constant vary between cortical areas suggests that Sst cells recorded in different areas have different membrane properties, leading to differences in excitability that cannot be explained solely by difference in cell size. Specifically, Sst neurons appeared to be less excitable than their counterparts recorded in motor cortex or somatosensory.

### 4.5 Cell type specific heterogeneity across cortical layers

#### 4.5.1 Inter-layer variability in visual cortex

To pursue our characterization of electrophysiological properties, we compared the variation of these properties according to anatomical factors like cortical layers. We started by comparing how linear and firing properties varied for a single cellular type according to the cortical layer it was recorded from, by considering neurons recorded in visual cortex from the Allen CTD (Supplementary Tables 13, 14, 15, 16,17, Figures 13, 14, 15, 16, 17).

**Figure 13:**
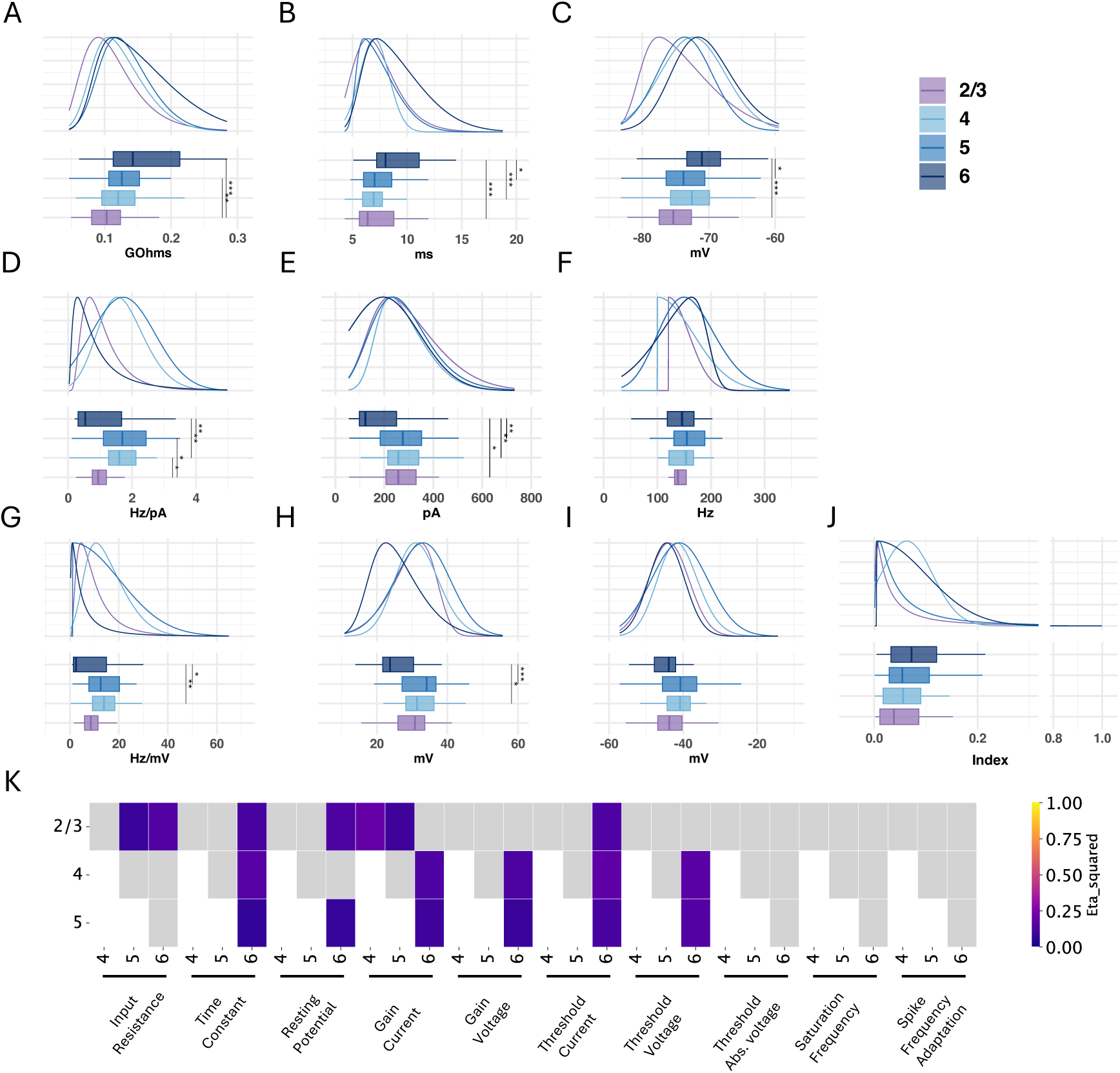
Comparison of electrophysiological properties of PValb neurons in Visual cortex between layers. Comparison of **A**: Input resistance, **B**: Time constant, **c**: Resting potential, **D**: Currentbased gain, **E**: Current-based threshold, **F**: Saturation frequency, **G**: Voltage-based gain, **H**: Voltage-based threshold, **I**: Voltage-based threshold reported to cell’s resting potential and **J** Spike Frequency Adaptation index, in different neuronal types. **K**: Heat map of effect size from KW test performed on features pairs. Only KW test which were statistically significant are colored, grey tiles represent non statistically significant differences. The analysis, made on PValb cells recorded at physiological temperature, with the use of synaptic blockers, from Visual cortex (Allen Cell Type Database). Summary statistics and distribution fit parameters are presented in Supplementary Table 13

**Figure 14:**
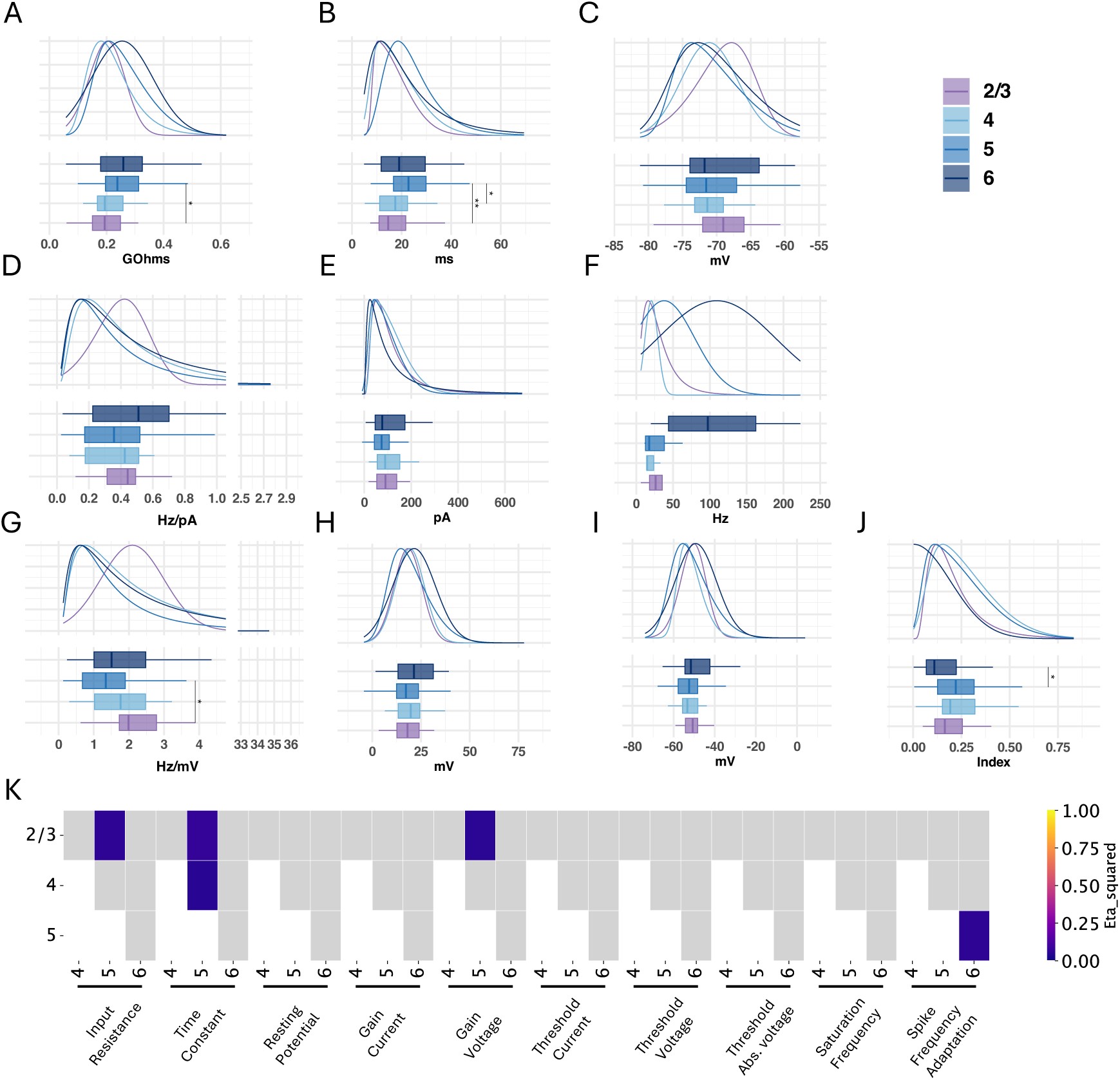
Comparison of electrophysiological properties of Sst neurons in Visual cortex between layers. Comparison of **A**: Input resistance, **B**: Time constant, **c**: Resting potential, **D**: Current-based gain, **E**: Current-based threshold, **F**: Saturation frequency, **G**: Voltage-based gain, **H**: Voltage-based threshold, **I**: Voltage-based threshold reported to cell’s resting potential and **J** Spike Frequency Adaptation index, in different neuronal types. **K**: Heat map of effect size from KW test performed on features pairs. Only KW test which were statistically significant are colored, grey tiles represent non statistically significant differences. The analysis, made on Sst cells recorded at physiological temperature, with the use of synaptic blockers, from Visual cortex (Allen Cell Type Database). Summary statistics and distribution fit parameters are presented in Supplementary Table 14

**Figure 15:**
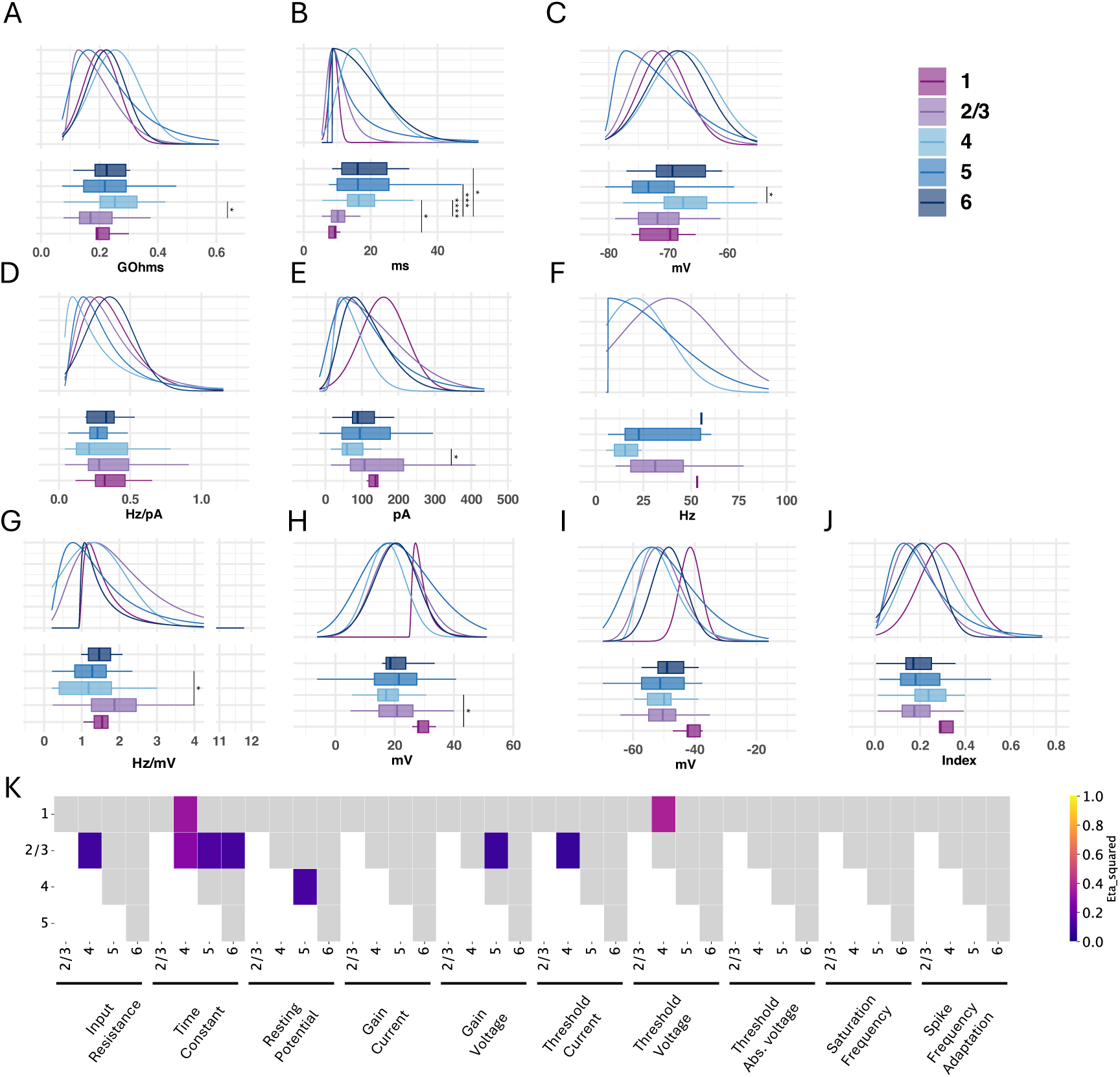
Comparison of electrophysiological properties of Htr3a neurons in Visual cortex between layers. Comparison of **A**: Input resistance, **B**: Time constant, **c**: Resting potential, **D**: Currentbased gain, **E**: Current-based threshold, **F**: Saturation frequency, **G**: Voltage-based gain, **H**: Voltage-based threshold, **I**: Voltage-based threshold reported to cell’s resting potential and **J** Spike Frequency Adaptation index, in different neuronal types. **K**: Heat map of effect size from KW test performed on features pairs. Only KW test which were statistically significant are colored, grey tiles represent non statistically significant differences. The analysis, made on Htr3a cells recorded at physiological temperature, with the use of synaptic blockers, from Visual cortex (Allen Cell Type Database). Summary statistics and distribution fit parameters are presented in Supplementary Table 15

**Figure 16:**
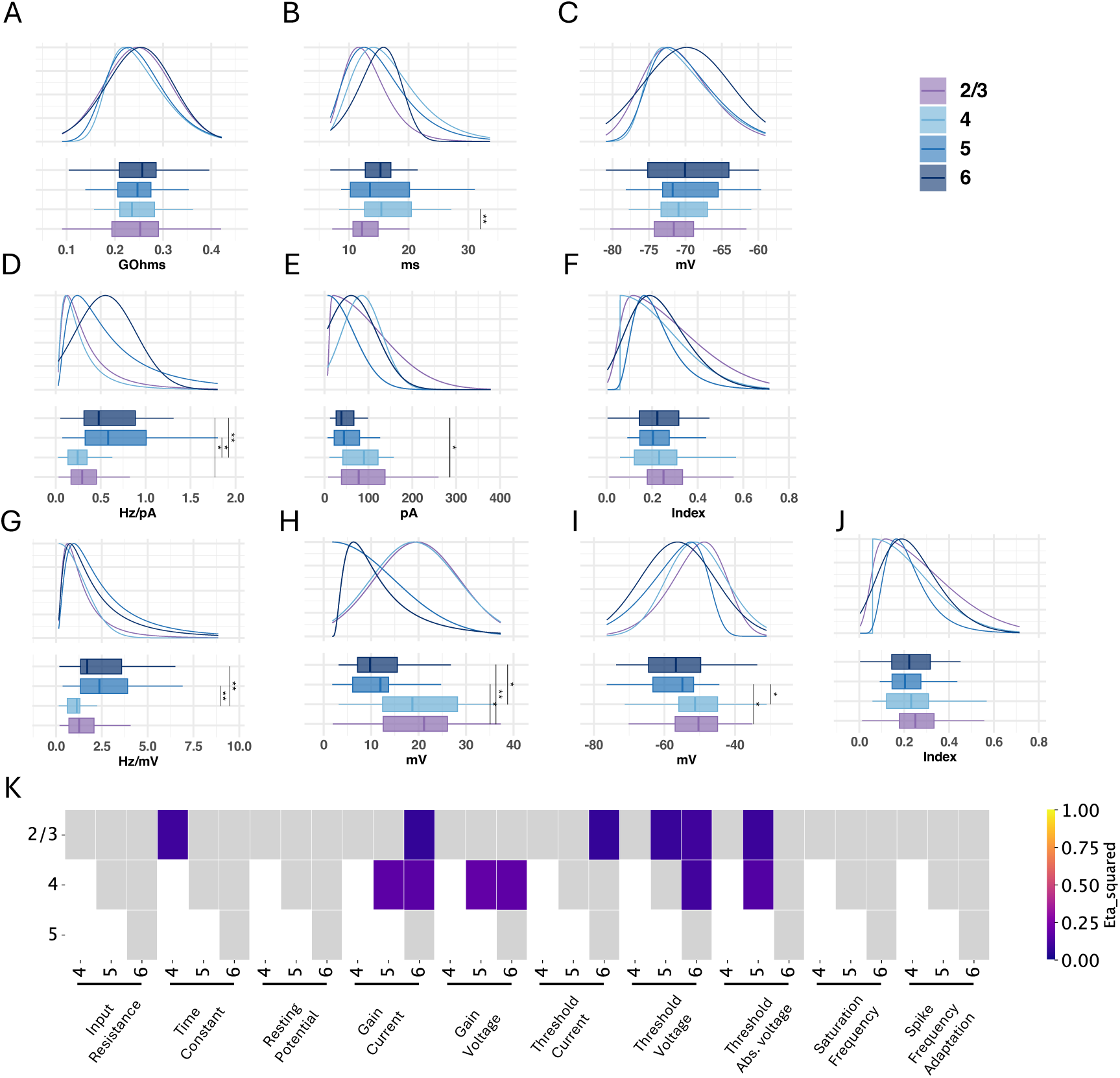
Comparison of electrophysiological properties of Vip neurons in Visual cortex between layers. Comparison of **A**: Input resistance, **B**: Time constant, **c**: Resting potential, **D**: Current-based gain, **E**: Current-based threshold, **F**: Saturation frequency, **G**: Voltage-based gain, **H**: Voltage-based threshold, **I**: Voltage-based threshold reported to cell’s resting potential and **J** Spike Frequency Adaptation index, in different neuronal types. **K**: Heat map of effect size from KW test performed on features pairs. Only KW test which were statistically significant are colored, grey tiles represent non statistically significant differences. The analysis, made on Vip cells recorded at physiological temperature, with the use of synaptic blockers, from Visual cortex (Allen Cell Type Database). Summary statistics and distribution fit parameters are presented in Supplementary Table 16.

**Figure 17:**
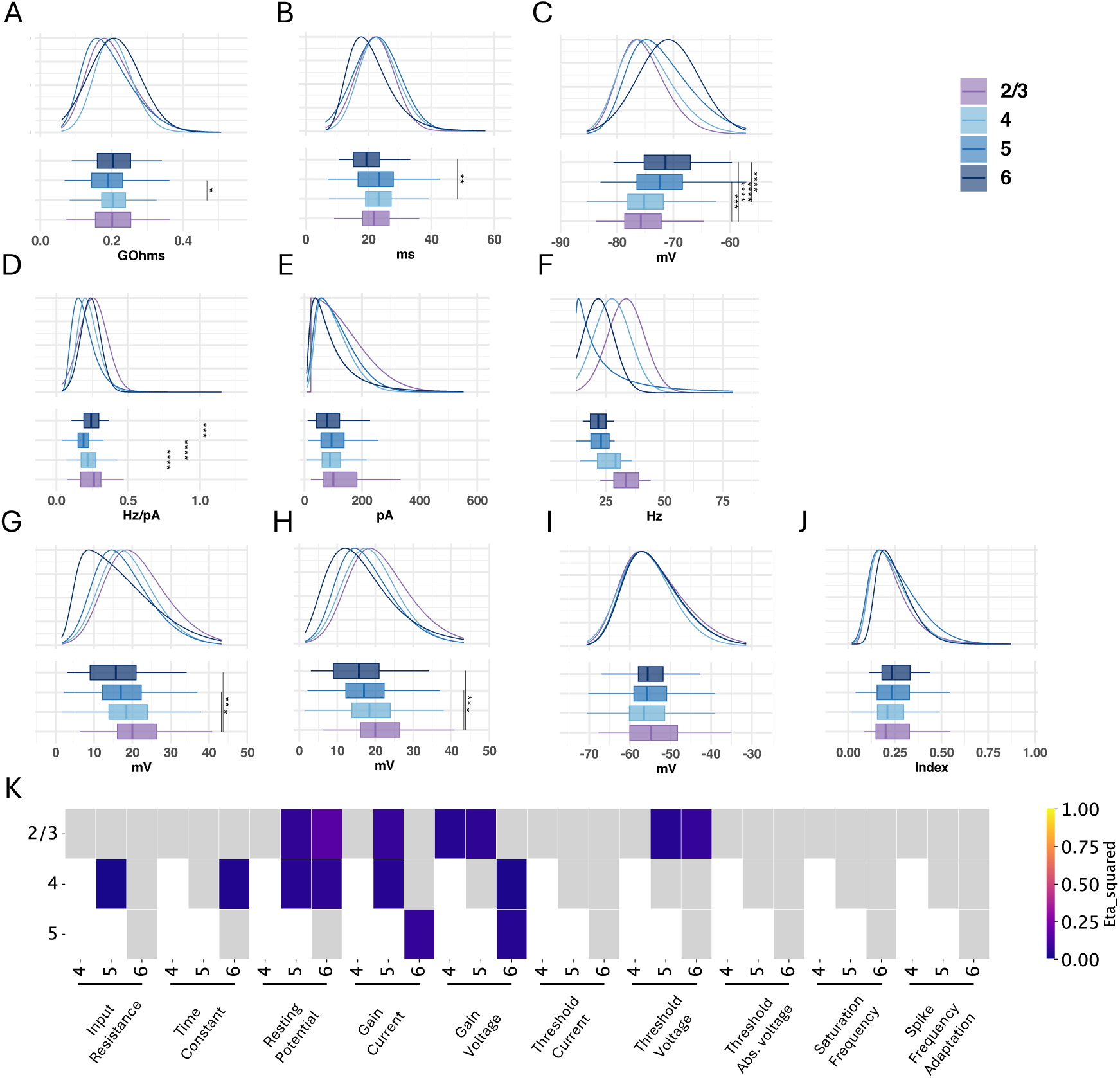
Comparison of electrophysiological properties of excitatory neurons in Visual cortex between layers. Comparison of **A**: Input resistance, **B**: Time constant, **c**: Resting potential, **D**: Currentbased gain, **E**: Current-based threshold, **F**: Saturation frequency, **G**: Voltage-based gain, **H**: Voltage-based threshold, **I**: Voltage-based threshold reported to cell’s resting potential and **J** Spike Frequency Adaptation index, in different neuronal types. **K**: Heat map of effect size from KW test performed on features pairs. Only KW test which were statistically significant are colored, grey tiles represent non statistically significant differences. The analysis, made on excitatory cells recorded at physiological temperature, with the use of synaptic blockers, from Visual cortex (Allen Cell Type Database). Summary statistics and distribution fit parameters are presented in Supplementary Table 17

This study revealed that linear properties of PValb neurons varied according to the layer they originated (KW test p≤0.0001 for Input resistance, p≤0.001 time constant and p≤0.001 for resting potential) (Figure 13 A, B, C respectively). Specifically, the input resistance of cells increased with cortical depth (Figure 13 A), even though the difference between neighboring layers was not significant, while it was between layer 2/3 and layer 5 (Dunn test, p adjusted ≤ 0.01, *η*^2^ = 0.083) and between layer 2/3 and layer 6 (Dunn test, p adjusted ≤ 0.0001, *η* = 0.142). Similarly, the time constant was longer for deeper layers (Figure 13 B). Yet, only neurons recorded in layer 6 statistically differed from L2/3 (Dunn test, p≤0.001, *η* = 0.115), L4 (Dunn test, p≤0.001, *η* = 0.151) and L5 (Dunn test, p≤0.05, *η* = 0.069). In the same way, neurons recorded in L6 were statistically more depolarized than neurons recorded in layer 2/3 (Dunn test, p≤0.001, *η* = 0.133) and in layer 4 (Dunn test, p≤0.05, *η* = 0.023) (Figure 13 C). PValb expressing interneurons also differed in their firing properties according to the cortical layers, especially regarding their gain ((Figure 13 D), and threshold (Figure 13 E) (KW test, p≤0.001, and p≤0.01 respectively). Specifically, neurons recorded in layer 2/3 and 6 had reduced gain compared to neuron in layer 4 and 5. Neurons recorded in layer 2/3 therefore statistically differed from neurons recorded in layer 4 (Dunn test, p≤0.05, *η*^2^ = 0.190), and 5 (Dunn test, p≤0.05, *η* = 0.094). Similarly, gain of layer 6 PValb neurons statistically differed from layer 4 (Dunn test, p≤0.01, *η* = 0.132) and layer 5 (Dunn test, p≤0.01, *η* = 0.094). Also, neurons recorded in layer 6 had lower threshold compared to neurons recorded in other layers (Supplementary Table 13), and only neurons from layer 6 statistically differed from other layers (L2/3-L6:p≤0.05, *η*^2^ = 0.134, L4-L6: p≤0.01, *η*^2^ = 0.175, L5-L6: p≤0.01, *η* = 0.117). These differences between layers were further confirmed when expressing gain (Figure 13 G) and threshold (Figure 13 H) in voltage input. However, the gain of layer 2/3 was not statistically different from layer 4 (Dunn test, p = 0.141) not from layer 5 (Dunn test, p = 0.266). Also, no difference was observed anymore between threshold of PValb neurons from layer 2/3 and 6 (Tukey test : p=0.175). However, by reporting the voltage-based threshold to the cells’ resting potential, no difference was observed between cells recorded in different layers (Figure 13 I)(ANOVA test, all pair-wise comparison Tukey test had adjusted p values ≥ 0.05). Similarly, no statistical differences was detected in the saturation frequency between layers (ANOVA, p=0.287) (Figure 13 F), nor in the spike frequency adaptation index (KW test: p=0.189) (Figure 13 J).

Interestingly, the layer-specific variability was not similar in other cell types (recorded in visual cortex, with the use of synaptic blockers, from the Allen Cell Type Database). For example, Sst neurons in visual cortex only displayed few inter-layer variability in their electrophysiological properties (Figure 14). As for PValb neurons, input resistance increased with cortical depth (Supplementary Table 13), yet only Sst neurons from layer 2/3 and 5 statistically differed (Dunn test, *η*^2^ = 0.055). Regarding the time constant, only the time constant of neurons recorded in layer 5 was different from those of upper layers (L2/3-L5 Dunn test, *η*^2^ = 0.070; L4-L5 Dunn test, *η*^2^ = 0.048). Also, an inter-layer variability was observed in the index of spike frequency adaptation between Sst neurons recorded in layer 5 and 6 (Dunn test, *η*^2^ = 0.057). However, no differences between Sst neurons recorded in different layers were observed for the resting potential (KW test, p=0.454), the gain when the input is expressed in current (KW test, p=0.2), the saturation frequency (KW test, p=0.246), the rheobase with current-based input (KW test, p=0.305) or voltage-based input (KW test, p=0.508 Table) or when reporting it to the cell’s resting potential (KW test, p=0.301). Yet, a slight difference was observed when expressing gain in voltage-based input between cells recorded in layer 2/3 and 5 (Dunn test, p≤0.05, *η*^2^ = 0.053).

Similarly, Htr3a neurons recorded in visual cortex in the presence of synaptic blockers (from the Allen Cell Type Database) displayed different layer-specific variability in their electrophysiological properties (Figure 15). Therefore, the input resistance of Htr3a neurons was statistically different only between layer 2/3 and 4 (Dunn test, p≤0.05, *η*^2^=0.101, Figure 15 A), while cells recorded in layer layer 4 were significantly more depolarized than cells recorded in layer 5 (Dunn test, p≤0.05, *η* =0.114, Figure 15 C), whereas cells’ resting membrane potential of other layers did not differed significantly. Htr3a neurons also had significantly different time constant according to the layer they originated from. More specifically, neurons recorded in upper layers had faster time constants compared to deeper layers, as neurons recorded from layer 2/3 statistically differed from neurons from layer 4 (Dunn test, p≤0.0001, *η*^2^=0.279,Figure 15 B), layer 5 (Dunn test, p≤0.001, *η*^2^=0.134) and layer 6 (Dunn test, p≤0.05, *η*^2^=0.112). Also, neurons from layer 1 had faster time constant than neurons from layer 4 (Dunn test, p≤0.05, *η*^2^=0.328). Similarly, current-based threshold of Htr3a neurons from layer 2/3 significantly differed with layer 4 neurons’ threshold (Dunn test, p≤0.05, *η*^2^=0.078,Figure 15 E). However, no differences were observed between layer in gain when the input is expressed in current (KW test, p=0.72, Figure 15 D), saturation frequency (KW test, p=0.205, Figure 15 F), spike frequency adaptation index (KW test, p=0.074, Figure 15 J), or when voltage-based threshold is reported to cell’s resting membrane potential (KW test, p=0.077, Figure 15 I). Yet, when expressing input in voltage, gain of Htr3a neurons from layer 2/3 was statistically different from neurons recorded in layer 5 (Dunn test, p≤0.05, *η*^2^=0.067, Figure 15 G). Similarly, statistical difference was observed between the threshold of Htr3a cells recorded in layer 1 and 4 (Dunn test, p≤0.05, *η*^2^=0.373, Figure 15 H), whereas no more difference was observed between layer 2/3 and layer 4 neurons.

We pursue our investigation by comparing electrophysiological feature of Vip expressing neurons recorded in visual cortex (from Allen Cell Type Database). Similarly to Htr3a cells, linear properties of Vip neurons only slightly varied according to the cortical layer, as no differences were observed in input resistance (ANOVA test, p=0.988, Figure 16 A), nor in resting membrane potential (ANOVA test, p=0.529, Figure 16 C). Yet, the time constant of neurons originating from layer 2/3 were significantly faster than the time constant of layer 4 neurons (Dunn test, p≤0.01, *η*^2^=0.094, Figure 16 B). The firing properties of Vip neurons were observed to be layer-dependent, as current-based gain was different between Layer 2/3 and 6 (Dunn test, p≤0.05, *η*^2^=0.069), between layer 4 and layer 5 (Dunn test, p≤0.05, *η*^2^=0.164), and between layer 4 and 6 (Dunn test, p≤0.01, *η* =0.157) (Figure 16 D). Similarly, when expressing threshold in current-based input, the only significant difference was observed between neurons from layer 2/3 and layer 6 (Dunn test, p≤0.05, *η* =0.071) (Figure 16 E). These differences in firing properties were further confirmed when reporting the input to the cell’s input resistance, therefore expressing gain in Hz/mV and the threshold in mV. Expressing the gain in voltage-based input, resulted in similar differences between layer 4 and 5 (Dunn test, p≤0.05, *η*^2^=0.181) and between layer 4 and 6 (Dunn test, p≤0.05, *η* =0.170) (Figure 16 G). However, the difference between layer 2/3 and 6 was not statistically significant anymore (Dunn test, p=0.093). Moreover, expressing threshold in voltage-based input increased the difference between layer 2/3 and 6 (Dunn test, p≤0.01, *η*^2^=0.098) (Figure 16 H). Furthermore, statistical differences were now observed between layer 2/3 and 5 (Dunn test, p≤0.05, *η*^2^=0.081), as well as between layers 4 and 6 (Dunn test, p≤0.05, *η*^2^=0.104). Also, reporting voltage-based threshold to cellular resting membrane potential reduced differences between layers as no more differences were observed between layer 4 and 6 (Tukey test, p=0.059). Yet, differences were still statistically significant between layer 2/3 and 5 (Tukey test, p≤0.05, *η*^2^=0.088), and between layer 4 and 5 (Tukey test, p≤0.05, *η*^2^=0.143) (Figure 16 I). However, no differences were observed between layers for saturation frequency (KW test, p=0.623, Figure 16 F) nor for spike frequency adaptation (KW test, p=0.423, Figure 16 J).

Finally, excitatory neurons from visual cortex had a different degree of layer-based variability in linear properties. Indeed, the input resistance of excitatory neurons only varied between layer 4 and 5 (Dunn test, p≤0.05, *η*^2^=0.016, Figure 17 A), and time constant only differed significantly between layers 4 and 6 (Dunn test, p≤0.01, *η* =0.032, Figure 17 B). However, excitatory neurons were observed to highly vary in their resting membrane potential between layer, as deeper layers cells had more depolarized resting potential. More specifically, statistical differences were observed as follow: L2/3-L5 (Dunn test, p≤0.001, *η*^2^=0.063), L2/3-L6 (Dunn test, p≤0.0001, *η*^2^=0.156), L4-L5 (Dunn test, p≤0.0001, *η*^2^=0.046) and L4-L6 (Dunn test, p≤0.0001, *η*^2^=0.056) (Figure 17 C). Excitatory neurons also varied in their current-based gain between layer 2/3 and 5 (Dunn test, p≤0.0001, *η*^2^=0.077), between layer 4 and 5 (Dunn test, p≤0.0001, *η*^2^=0.044) and between layer 5 and 6 (Dunn test, p≤0.001, *η*^2^=0.071) (Figure 17 D). These differences slightly differed when expressing gain in term of voltage-based input, as layer 2/3 cells differed from layer 4 (Dunn test, p≤0.01, *η*^2^=0.039) and from layer 5 (Dunn test, p≤0.001, *η*^2^=0.057), whereas cells from layer 6 significantly differed from layer 4 (Dunn test, p≤0.05, *η*^2^=0.022) and from layer 5 (Dunn test, p≤0.05, *η*^2^=0.039) (Figure 17 G). However, no differences were observed in saturation frequency (KW test, p=0.088, Figure 17 F), spike frequency adaptation (KW test, p=0.186, Figure 17 J) or current-based threshold (KW test, p=0.108, Figure 17 E). When expressing threshold in voltage-based input, excitatory cells were observed to have lower firing threshold with cortical depth, as neurons from layer 2/3 differed from layer 5 (Dunn test, p≤0.05, *η*^2^=0.039) and from layer 6 (Dunn test, p≤0.01, *η*^2^=0.072) (Figure 17 H). Yet, no more differences were observed between layers when reporting voltage-based threshold to the cell’s resting membrane potential (KW test, p=0.315, Figure 17 I).

These different examples highlighted the fact that for a given neuron type, the different linear and firing properties may vary according to the cortical layer in which neurons originate from. Furthermore, these layer-dependent variations appeared to be specific to each cell type, as some categories like PValb expressing neurons highly vary according to the cortical area, whereas Sst expressing neurons were highly homogeneous according to the layer (see panel K in Figures 13, 14, 15, 16 and 17).

#### 4.5.2 Inter-layer variability in motor cortex

To understand if this layer-based variability was specific to visual cortex, we also compared linear and firing properties of PValb, Sst, Vip and Excitatory cells in the motor cortex, recorded at room temperature without the use of synaptic blockers (Scala 2021 database, Supplementary Tables 18, 19, 20, 21, Figures 18, 19, 20, 21).

**Figure 18:**
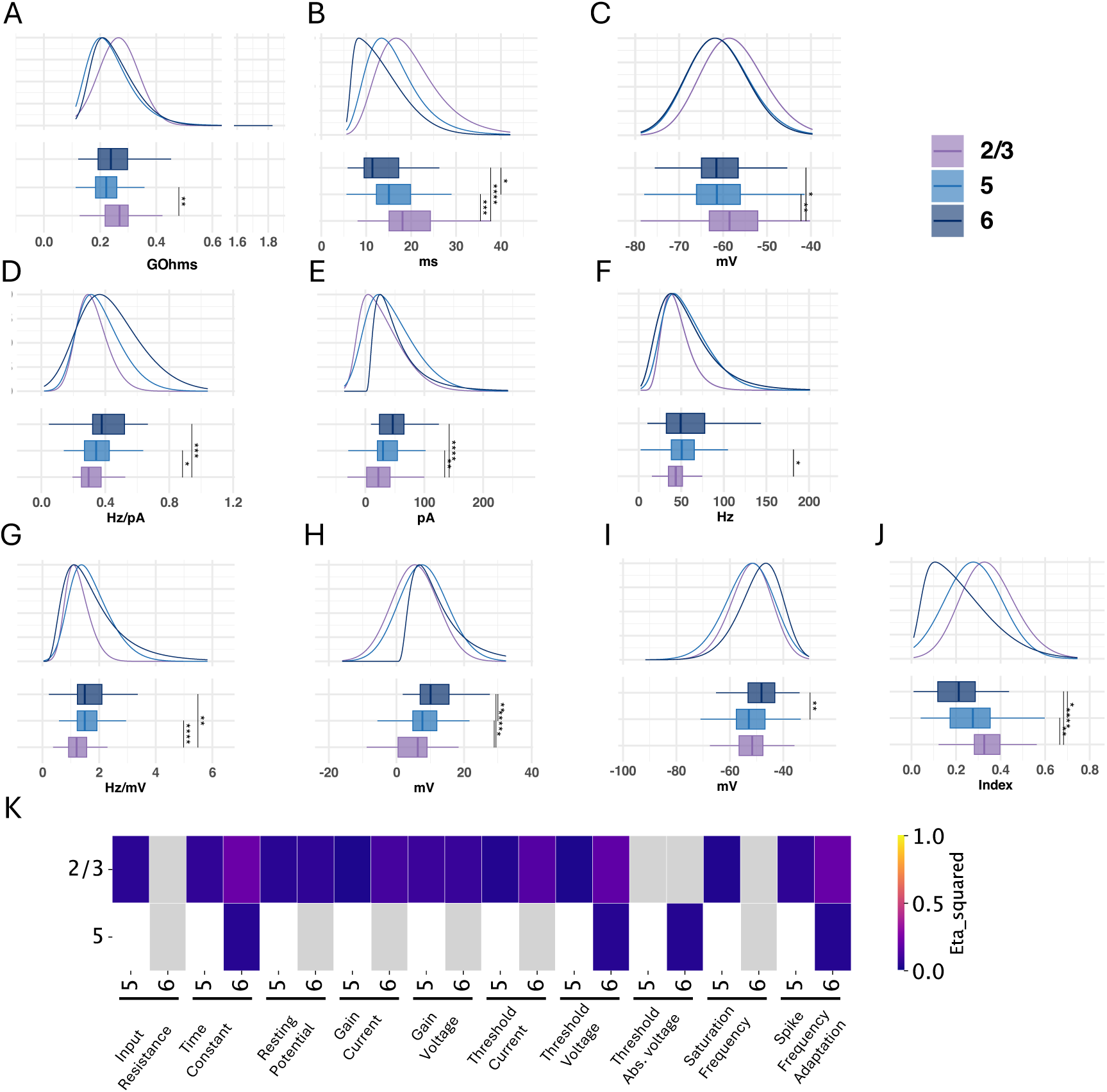
Comparison of electrophysiological properties of Sst neurons in Motor cortex between layers at room temperature. Comparison of **A**: Input resistance, **B**: Time constant, **c**: Resting potential, **D**: Current-based gain, **E**: Current-based threshold, **F**: Saturation frequency, **G**: Voltage-based gain, **H**: Voltage-based threshold, **I**: Voltage-based threshold reported to cell’s resting potential and **J** Spike Frequency Adaptation index, in different neuronal types. **K**: Heat map of effect size from KW test performed on features pairs. Only KW test which were statistically significant are colored, grey tiles represent non statistically significant differences. The analysis was made on Sst cells recorded at room temperature, without the use of synaptic blockers, from Motor cortex (Scala 2021 database). Summary statistics and distribution fit parameters are presented in Supplementary Table 18

**Figure 19:**
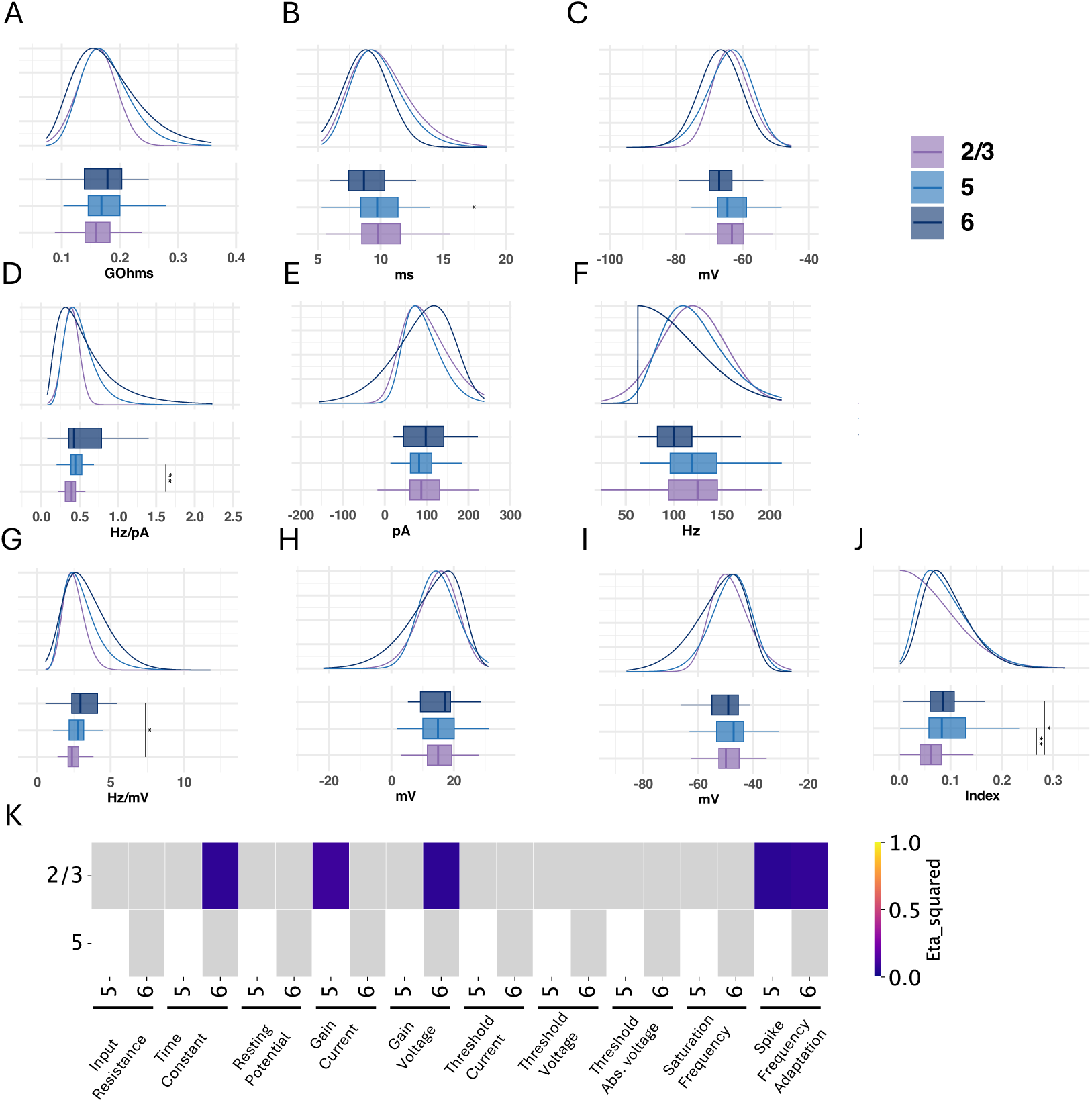
Comparison of electrophysiological properties of PValb neurons in Visual cortex between layers at room temperature. Comparison of **A**: Input resistance, **B**: Time constant, **c**: Resting potential, **D**: Current-based gain, **E**: Current-based threshold, **F**: Saturation frequency, **G**: Voltage-based gain, **H**: Voltage-based threshold, **I**: Voltage-based threshold reported to cell’s resting potential and **J** Spike Frequency Adaptation index, in different neuronal types. **K**: Heat map of effect size from KW test performed on features pairs. Only KW test which were statistically significant are colored, grey tiles represent non statistically significant differences. The analysis was made on PValb cells recorded at room temperature, without the use of synaptic blockers, from Motor cortex (Scala 2021 database). Summary statistics and distribution fit parameters are presented in Supplementary Table 19

**Figure 20:**
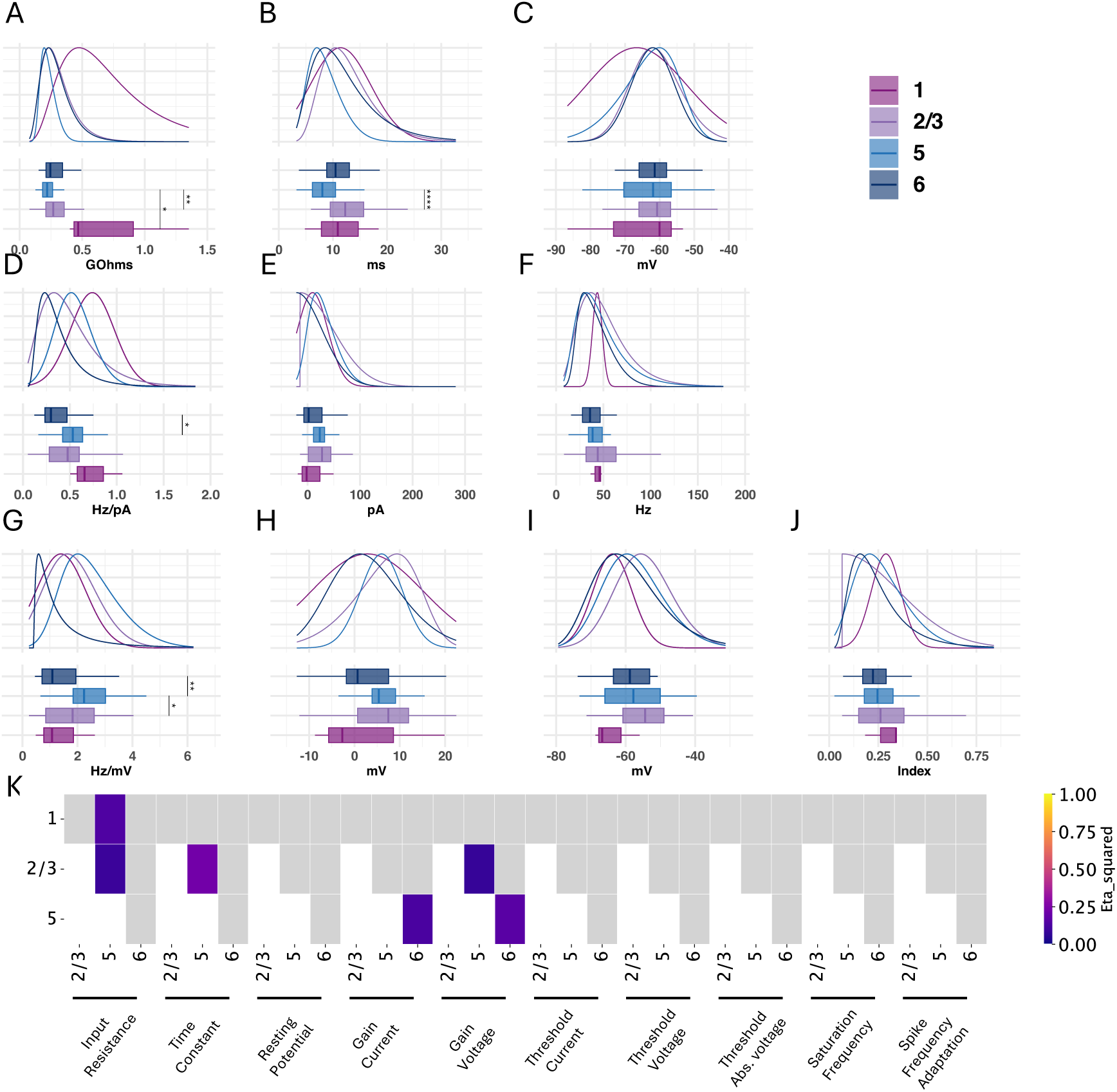
Comparison of electrophysiological properties of Vip neurons in Motor cortex between layers at room temperature. Comparison of **A**: Input resistance, **B**: Time constant, **c**: Resting potential, **D**: Current-based gain, **E**: Current-based threshold, **F**: Saturation frequency, **G**: Voltage-based gain, **H**: Voltage-based threshold, **I**: Voltage-based threshold reported to cell’s resting potential and **J** Spike Frequency Adaptation index, in different neuronal types. **K**: Heat map of effect size from KW test performed on features pairs. Only KW test which were statistically significant are colored, grey tiles represent non statistically significant differences. The analysis was made on Vip cells recorded at room temperature, without the use of synaptic blockers, from motor cortex (Scala 2021 database). Summary statistics and distribution fit parameters are presented in Supplementary Table 20

**Figure 21:**
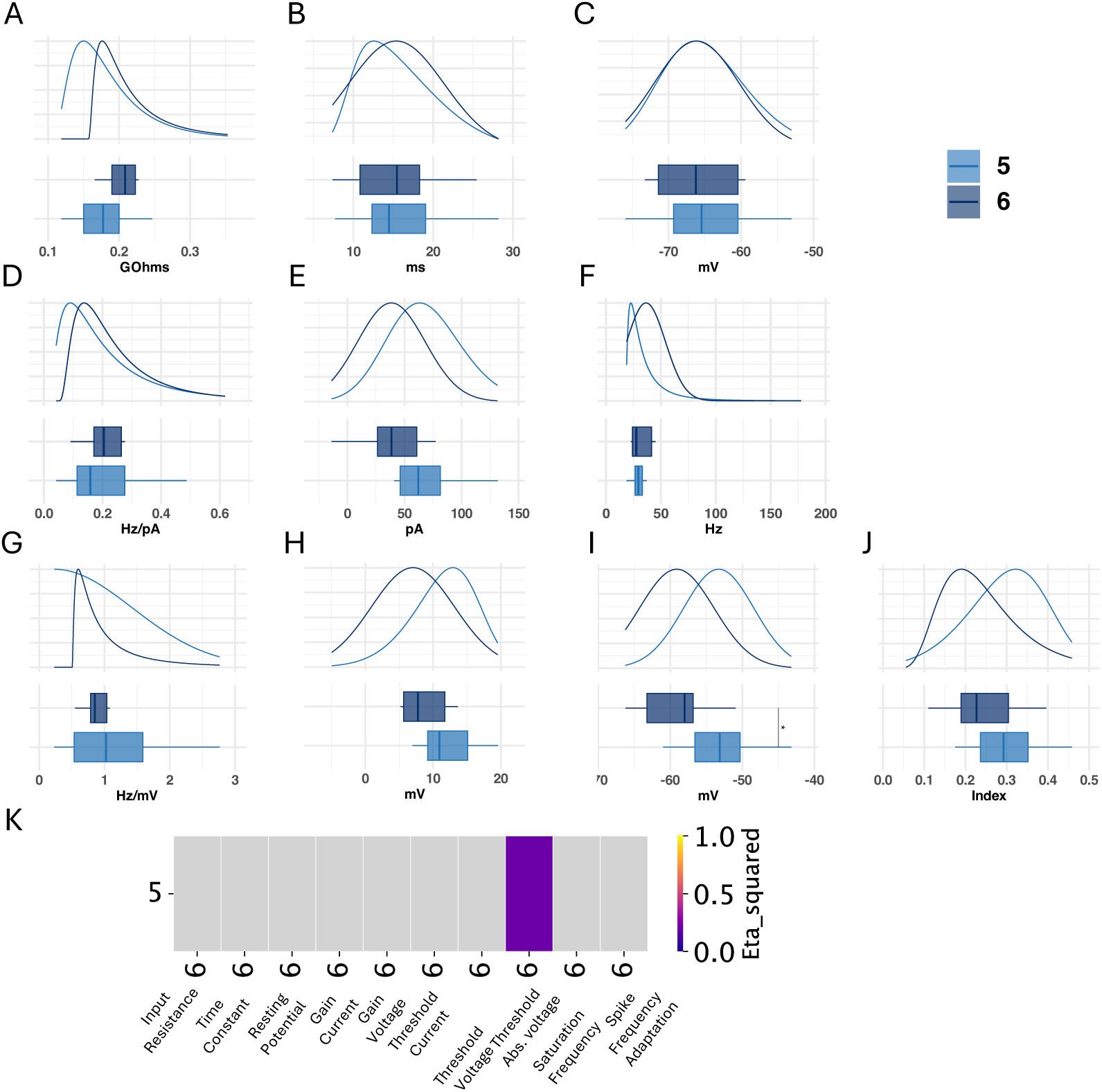
Comparison of electrophysiological properties of excitatory neurons in Motor cortex between layers at room temperature. Comparison of **A**: Input resistance, **B**: Time constant, **c**: Resting potential, **D**: Current-based gain, **E**: Current-based threshold, **F**: Saturation frequency, **G**: Voltage-based gain, **H**: Voltage-based threshold, **I**: Voltage-based threshold reported to cell’s resting potential and **J** Spike Frequency Adaptation index, in different neuronal types. **K**: Heat map of effect size from KW test performed on features pairs. Only KW test which were statistically significant are colored, grey tiles represent non statistically significant differences. The analysis was made on excitatory cells recorded at room temperature, without the use of synaptic blockers, from motor cortex (Scala 2021 database). Summary statistics and distribution fit parameters are presented in Supplementary Table 21

Especially, Sst cells recorded in the motor cortex displayed layer-dependent variability in their linear properties, with significantly different input resistance between cells recorded in layer 2/3 and 5 (Dunn test, p≤0.01, *η*^2^=0.056, Figure 18 A), faster time constants in deeper layer and significant differences as follow L2/3-L5 (Dunn test, p≤0.001, *η* =0.065), L2/3-L6 (Dunn test, p≤0.0001, *η* =0.201) and L5-L6 (Dunn test, p≤0.05, *η* =0.046) (Figure 18 B). Also, neurons from layer 2/3 were more depolarized than neurons from layer 5 (Tukey test, p≤0.01, *η* =0.048) and from layer 6 (Tukey test, p≤0.05, *η* =0.274) (Figure 18 C). Sst neurons also differed in their firing properties, as Sst neurons recorded in layer 2/3 were observed to have a reduced excitability compared to neurons recorded in deeper layers. Notably statistical differences were observed in current-based gain between neurons from layer 2/3 and layer 5 (Dunn test, p≤0.05, *η*^2^=0.023) and between layer 2/3 and layer 6 (Dunn test, p≤0.001, *η* =0.110) (Figure 18 D), as well as in current-based threshold between neurons from layer 2/3 and layer 5 (Dunn test, p≤0.01, *η* =0.036) and between neurons from layer 2/3 and layer 6 (Dunn test, p≤0.01, *η* =0.148) (Figure 18 E). Similar differences were observed when expressing input in linear voltage for gain (L2/3-L5: Dunn test, p≤0.0001, *η* =0.088; L2/3-L6: Dunn test, p≤0.01, *η* =0.081) (Figure 18 G) and for threshold (L2/3-L5: Dunn test, p≤0.05, *η* =0.026; L2/3-L6: Dunn test, p≤0.0001, *η* =0.176; L5-L6: Dunn test, p≤0.01, *η* =0.044) (Figure 18 H). However, reporting voltage input to the cell’s resting potential reduced the differences between the layers, as the only significant difference was observed between layer 5 and layer 6 (Dunn test, p≤0.01, *η*^2^=0.040) (Figure 18 I). Additionally, neurons from layer 2/3 had lower saturation frequency compared to neurons from layer 5 (Dunn test, p≤0.0001, *η*^2^=0.034)(Figure 18 F). Finally, Sst neurons in these different layers had significantly different spike frequency adaptation index. the differences observed were as follow: L2/3-L5 (Dunn test, p≤0.01, *η*^2^=0.060), L2/3-L6 (Dunn test, p≤0.0001, *η*^2^=0.195) and L5-L6 (Dunn test, p≤0.05, *η*^2^=0.041) (Figure 18 J).

On the other hand, properties of PValb cells were observed to moderately vary according to the cortical layer, as only neurons from layer 2/3 significantly differed from neurons in layer 6 in their time constant (Tukey test, p≤0.05, *η*^2^=0.062, Figure 19 B), and from neurons from layer 5 in their current-based gain (Dunn test, p≤0.01, *η* =0.091) (Figure 19 D). The later difference, however, was not observed when expressing input in linear voltage, but difference was observed between layer 2/3 and 6 (Dunn test, p≤0.0001, *η*^2^=0.056) (Figure 19 G). Also, PValb neurons from layer 2/3 had lower spike frequency adaptation index than neurons recorded in layer 5 (Dunn test, p≤0.01, *η*^2^=0.053) and layer 6 (Dunn test, p≤0.05, *η*^2^=0.068) (Figure 19 J). However, no significant differences were observed between PValb neurons recorded in different layer in input resistance (KW test, p=0.12) (Figure 19 A),resting potential (KW test p≤0.05, Dunn test ≥0.05 for all pairs) (Figure 19 C), current-based threshold (KW test, p=0.808) (Figure 19 E), voltage-based threshold (KW test, p=0.959) (Figure 19 H), absolute-voltage-based threshold (KW test, p=0.326) (Figure 19 I) nor saturation frequency (KW test, p=0.123) (Figure 19 F).

Similarly, Vip cells recorded in motor cortex showed only few layer-dependent differences. Indeed, Vip cells were observed to have a layer-dependent input resistance, with higher input resistance in upper layer as layer 5 cells differed significantly from layer 1 (Dunn test, p≤0.05, *η*^2^=0.134) and from layer 2/3 (Dunn test, p≤0.01, *η* =0.087) cells (Figure 20 A). Similarly, Vip neurons from layer 5 had faster time constants than cells from layer 2/3 (Dunn test, p≤0.0001, *η* =0.220) (Figure 20 B. Finally, Vip neurons in layer 6 had significantly lower current-based gain than neurons from layer 5 (Dunn test, p≤0.05, *η* =0.017) (Figure 20 D). This difference was still observable when expressing gain in term of voltage-based input, as neurons from layer 5 had higher voltage-based gain than neurons in layer 6 (Dunn test, p≤0.01, *η*^2^=0.159), and in layer 2/3 (Dunn test, p≤0.05, *η* =0.077) (Figure 20 G). No other differences were observed between Vip neurons recorded in different layers being for resting potential (ANOVA test, p=0.304), current-based threshold (KW test, p =0.202), saturation frequency (KW test, p=0.394), voltage-based threshold (KW test, p=0.302), absolute-voltage-based threshold (ANOVA test, p=0.088) or spike frequency adaptation (KW test, p=0.695) (Figure 20 C, E, F, H, I and J respectively).

Finally, electrophysiological properties of excitatory neurons recorded from layer 5 and 6 were highly homogeneous, as the only statistically significant difference was observed when expressing rheobase in absolute-voltage based input (ANOVA test, p≤0.05, *η*^2^=0.205) (Figure 21 I). All other properties were similar when comparing between these layers: input resistance (KW test, p=0.095), time constant (ANOVA test, p=0.845), resting potential (ANOVA test, p=0.671), current-based gain (KW test, p=0.505), current-based threshold (ANOVA test, p=0.075), saturation frequency (KW test, p=0.739), voltage-based gain (KW test, p=0.947), voltage-based threshold (ANOVA test, p=0.096) and spike frequency adaptation (ANOVA test, p=0.303) (Figure 21 A, B, C, D, E, F, G, H, and J respectively). However, the low number of observation (18 in layer 5 and 6 in layer 6) and the restricted layer sampling suggest that a more in depth review of this cell type could lead to more robust conclusions about the heterogeneity of electrophysiological properties of excitatory neurons in motor cortex.

Therefore, it appeared that membrane properties and excitability of Sst cells in motor cortex was layer dependent, with varying linear and firing properties across layers, whereas PValb and Vip neurons only slightly vary across layers in some properties.

#### 4.5.3 Summary of cell type inter-layer heterogeneity

Overall we find that cell type specific inter-layer variability in motor cortex is not the same in visual cortex (see panel K in Figures 13, 14, 15, 16 and 17) and motor cortex (see panel K in Figures 18, 19, 20 and 21). Whereas the properties of PValb neurons are the most variable per layer in visual cortex, this type among the less variable cell type in motor cortex. On the contrary, whereas Sst neurons were the most homogeneous cell type across layers of visual cortex, it was the most variable cell type in motor cortex.

## 5 Conclusion

Since the pioneer work of Hodgkin and Huxley (Hodgkin and Huxley, 1939; Hodgkin, 1948; Hodgkin et al., 1949; Hodgkin and Huxley, 1952), the study of neuronal electrophysiological properties has been an extensive field of study for decades. Many laboratories have addressed the electrophysiological properties of various neuronal types in various brain regions resulting in an increasing number of publicly available databases (Staff et al., 2000; Milior et al., 2016; Harrison et al., 2015; da Silva Lantyer et al., 2018; Gouwens et al., 2019; Scala et al., 2019, 2021; Vitale et al., 2021; Petersen et al., 2022; Valero et al., 2022). The publication of raw data alongside articles, is a growing habit in the scientific field, and is one of the Open Science aspects aiming at improving research sustainability (Janssens et al., 2023; Richter, 2024). To support this initiative, the establishment of common data files (Teeters et al., 2015; Rübel et al., 2022) and common online repositories format constitutes major steps toward more integrated scientific cooperation and reproducibility. Yet, even though two studies rely on similar experimental protocol (e.g.: current-clamp), the measurements can be difficult to compare. Indeed, discrepancies in experimental procedures (i.e.: stimulus range, fitting procedure, …) can greatly influence the results (Ballbé et al, in preparation).

The observations presented above indicate that neurons exhibit a multi-dimensional variation. For example when comparing across layers, all neuronal types displayed varying degrees of layer-dependent heterogeneity in their linear and firing properties. Some, like PValb in the visual cortex, were highly variable, while others were more homogeneous. Moreover, these cell-type layer-dependent variations in neuronal properties also seemed to be related to the cortical area from which the neurons originated. Consequently, neuronal heterogeneity appeared to be a highly dimensional aspect of neuronal biophysics.

### 5.1 Limit of inter-database comparisons

By gathering databases from different laboratories, we also addressed the differences in electrophysiological properties due to experimental parameters, like recording at lower than physiological temperature (Figure 4) or the use of synaptic blockers, which in both cases influenced membrane linear properties, inducing an increase in neuronal excitability, which was in agreement with previous reports (Hook, 2020; Scala et al., 2021). Synaptic blockers had comparable effect on linear and firing properties than when we compared neurons recorded *in vitro* and *in vivo*, decreasing input resistance, and increasing gain and threshold. Under the assumption that these differences can be attributed mainly to the difference of synaptic context (Destexhe et al., 2003), reporting current input to cell’s input resistance had different effects in both situations. While it entirely compensated differences between *in vitro* and *in vivo* cells firing properties distributions, the firing properties still differed between neurons recorded with or without synaptic blockers. This suggests that the use of synaptic blockers, mimicking the decreased level of synaptic inputs arriving on a neuron compared to *in vivo* conditions, alters membrane properties and increasing neuronal excitability differently. The comparison of cells recorded *in vitro* and *in vivo*, was made on untargeted population of visual cortex cells, contrary to the investigation of synaptic blockers which was made on specific cell types (Sst or PValb neurons). However, electrophysiological properties can greatly vary according to cell type, and layer. A more precise, cell-type specific investigation of the difference between *in vitro* and *in vivo* condition could help understand these differences. However, it should be noted that the role of background synaptic activity on the differences in excitability between neurons recorded *in vitro* and *in vivo* has been challenged, as the modulation of synaptic activity in slice recording during dynamic-clamp recording did not compensate for difference with *in vivo* recordings, suggesting that properties of cells in slices do not only differ because of different background activity but potentially from different neuromodulatory mechanisms (Fernandez et al., 2018). Other aspects of experimental design, such as animal age (Pan et al., 2016; Halfmann et al., 2023), can influence electrophysiological properties and limit the comparability of databases. In summary, our findings demonstrate the significant impact that variations in experimental design can have on neuronal properties, which can hinder the interoperability and comparison of results across databases.

### 5.2 Multifaceted neuronal heterogeneity

We relied on highly detailed databases (Gouwens et al., 2019; Scala et al., 2019, 2021) to investigate the variation of linear and firing properties in different cell types, according to different factors. Our results relied on both the analysis of central tendencies and population distributions, and highlighted the higher excitability and faster membrane dynamics of PValb cells compared to other commonly defined neuronal types, and was consistent with the qualification of PValb neurons as “Fast-Spiking” cells, while excitatory, Sst, Htr3a and Vip neurons were considered as “Regular-Spiking” (Contreras, 2004). The persistence of these inter cell-types variations when expressing firing properties in linear voltage suggested that these inter cell-types variations were not due to differences in somatic size, but rather to intrinsically different membrane properties.

While the relative properties between cell types were also observable in cells recorded in motor cortex, these membrane properties also appeared to be dependent on the cortical area. By comparing Sst cells recorded in the visual and somatosensory cortex, under similar experimental conditions, we observed that linear and firing properties were different in different cortical areas. Especially, cells recorded in visual cortex appeared to be less excitable compared to cells recorded in somatosensory areas. Interestingly, we also observed that Sst neurons from visual cortex had reduced excitability compared to motor cortex cells. The expression of cellular input in linear voltage did not attenuated the difference of firing properties, suggesting that these differences were not due to variation in cell areas across cortical areas, which corroborates findings from previous studies between visual and somatosensory cortex (Scala et al., 2019) and between primary and secondary motor cortex (Ueta et al., 2014) stating that similar cell types from different cortical regions have distinct properties.

We also benefited from the cell layer annotations in the Allen CTD and Scala 2021 databases. Our results provided different insights on neuronal variability. First, in agreement with previous studies, although these were restricted to pairs of layers (Huggenberger et al., 2009; Prönneke et al., 2015; Scala et al., 2019), or on a distance-related based (Pastoll et al., 2020), our work highlighted that linear and firing properties appeared to vary according across all cortical layers. Secondly, our study revealed that inter-layer variability is a complex properties of cortical neurons, exhibiting both cell-type-dependent and area-specific variations. While some of the variability could be accounted for by the rescaling of current input to cell’s input resistance, other aspects of neuronal network could explain the observed difference within a cell type, as it has been observed for example that cell with common projection targets tend to have similar physiological properties (Hattox and Nelson, 2007).

### 5.3 Impact of neuronal heterogeneity on network properties

Our work highlighted the complex task of describing the neuronal heterogeneity characteristic of neuronal networks, depending on the transcription profiles of cells and their anatomical context, with a diverse variability of inter-cortical and layer-specific properties for different cell types. The precise characterization of neuronal variability represent a crucial step toward understanding of neuronal networks’ properties as neuronal variability is highly implicated in diverse physiological processes. Increasing heterogeneity in neuronal biophysical properties over population of neurons has been reported to confer the population higher coding and information conveying capacities (Shamir and Sompolinsky, 2006; Padmanabhan and Urban, 2010), or participate in human behavioral variability (Findling and Wyart, 2021; Waschke et al., 2021). On the other hand, reduction of neuronal distance to threshold heterogeneity has been observed in *post-mortem* human epileptogenic tissues, suggesting potential pathological role of variability decrease in neuronal network (Rich et al., 2022).

The computational investigation of neuronal heterogeneity has drawn increased attention. To date, these models have relied on single (Shamir and Sompolinsky, 2006; Hutt et al., 2023; Muscinelli et al., 2019), double (excitatory/inhibitory) (Roxin et al., 2011; Muscinelli et al., 2019; Rich et al., 2022) or three population network models (Crockett et al., 2015; Duarte and Morrison, 2019), which, with the exception of (Duarte and Morrison, 2019), have relied on theoretical heterogeneity (e.g. normal distributions with various standard deviations around an assumed mean).

We believe that precisely describing the heterogeneity of cell types across cortical areas and layers is a crucial step toward large-scale neuronal network modeling. This approach allows us to address the emergence of network properties resulting from neuronal variability in a biologically-constrained manner. Our work contributes to a more detailed understanding of the relationships and variability of cellular properties across different common biological factors that constitute key aspects of biologically-constrained network models.

## 6 Supplementary Materials

**Table 6:** Table of descriptive statistics and fitting parameters for figure 3. **IR**: Input Resistance (*G*Ω),**TC**: Time Constant(ms), *V_rest_*: Resting membrane potential (mV), **Gain current** (Hz/pA),**Gain voltage**: (Hz/mV), **Threshold current**: (pA), *V ^Rel^*: Threshold Voltage (mV), *V_θ_*: Threshold Absolute Voltage (mV), **Saturation**: Saturation Frequency (Hz), **SFA**: Spike Frequency Adaptation (no unit)

| Feature | Population | N | Mean | Standard Deviation | Min | $Q_1$ | Median | $Q_3$ | Max | IQR | Distribution Fit |
| --- | --- | --- | --- | --- | --- | --- | --- | --- | --- | --- | --- |
| IR | In Vitro | 1110 | 0.22 | 0.08 | 0.06 | 0.17 | 0.21 | 0.26 | 0.62 | 0.10 | LogNormal(-0.066; 0.267; 0.277) |
|  | In Vivo | 88 | 0.17 | 0.27 | 0.02 | 0.05 | 0.08 | 0.13 | 1.30 | 0.07 | LogNormal(0.019; 1.147; 0.07) |
| TC | In Vitro | 1110 | 20.31 | 8.50 | 5.17 | 13.41 | 19.76 | 25.80 | 69.40 | 12.39 | LogNormal(-6.775; 0.312; 25.807) |
|  | In Vivo | 88 | 17.95 | 11.91 | 0.37 | 10.13 | 14.57 | 21.66 | 69.84 | 11.53 | LogNormal(-3.688; 0.493; 19.092) |
| $V_{rest}$ | In Vitro | 1110 | -72.03 | 5.48 | -85.36 | -75.96 | -72.53 | -68.33 | -54.95 | 7.62 | SkewedGaussian(-78.058; 8.149; 2.402) |
|  | In Vivo | 88 | -79.44 | 9.08 | -117.01 | -85.70 | -79.60 | -73.08 | -62.70 | 12.62 | SkewedGaussian(-69.895; 13.138; -2.221) |
| Gain current | In Vitro | 1018 | 0.31 | 0.26 | 0.03 | 0.17 | 0.23 | 0.35 | 2.76 | 0.17 | LogNormal(-0.002; 0.628; 0.251) |
|  | In Vivo | 86 | 0.14 | 0.09 | 0.02 | 0.09 | 0.12 | 0.15 | 0.57 | 0.06 | LogNormal(-0.071; 0.334; 0.194) |
| Gain voltage | In Vitro | 1018 | 1.53 | 1.75 | 0.13 | 0.89 | 1.19 | 1.67 | 34.72 | 0.78 | LogNormal(-0.016; 0.609; 1.237) |
|  | In Vivo | 86 | 1.71 | 1.52 | 0.11 | 0.83 | 1.39 | 2.12 | 10.99 | 1.29 | LogNormal(-0.338; 0.645; 1.664) |
| Threshold current | In Vitro | 1018 | 104.59 | 74.36 | -16.40 | 53.14 | 86.59 | 136.04 | 552.96 | 82.90 | LogNormal(-28.578; 0.521; 116.184) |
|  | In Vivo | 86 | 226.56 | 120.49 | -14.14 | 142.92 | 235.10 | 321.93 | 562.11 | 179.00 | N(226.558; 119.791) |
| $V_{\theta}^{Rel}$ | In Vitro | 1018 | 18.86 | 8.70 | -6.16 | 12.73 | 18.05 | 24.27 | 77.92 | 11.55 | LogNormal(-29.082; 0.179; 47.18) |
|  | In Vivo | 86 | 27.74 | 39.03 | -12.09 | 11.71 | 17.60 | 24.85 | 199.00 | 13.13 | LogNormal(-48.887; 0.154; 65.179) |
| $V_{\theta}$ | In Vitro | 1018 | -53.33 | 8.24 | -76.33 | -58.65 | -54.27 | -48.62 | 4.05 | 10.03 | SkewedGaussian(-61.722; 11.759; 2.077) |
|  | In Vivo | 86 | -51.78 | 39.91 | -104.66 | -69.95 | -62.30 | -53.93 | 116.13 | 16.02 | LogNormal(-115.121; 0.467; 55.926) |
| Saturation | In Vitro | 159 | 34.32 | 29.59 | 3.69 | 14.70 | 27.69 | 40.87 | 222.93 | 26.17 | LogNormal(0.247; 0.76; 25.587) |
|  | In Vivo | 50 | 34.93 | 18.16 | 4.47 | 22.24 | 34.60 | 47.85 | 79.31 | 25.62 | SkewedGaussian(14.855; 26.954; 2.398) |
| SFA | In Vitro | 1083 | 0.24 | 0.13 | 0.00 | 0.15 | 0.21 | 0.31 | 0.87 | 0.17 | LogNormal(-0.142; 0.331; 0.358) |
|  | In Vivo | 88 | 0.31 | 0.10 | 0.01 | 0.25 | 0.30 | 0.37 | 0.57 | 0.12 | LogNormal(-20.569; 0.005; 20.878) |

**Table 7:** Table of descriptive statistics and fitting parameters for figure 4. **IR**: Input Resistance (*G*Ω),**TC**: Time Constant(ms), *V_rest_*: Resting membrane potential (mV), **Gain current** (Hz/pA),**Gain voltage**: (Hz/mV), **Threshold current**: (pA), *V ^Rel^*: Threshold Voltage (mV), *V_θ_*: Threshold Absolute Voltage (mV), **Saturation**: Saturation Frequency (Hz), **SFA**: Spike Frequency Adaptation (no unit)

| Feature | Population | N | Mean | Standard Deviation | Min | $Q_1$ | Median | $Q_3$ | Max | IQR | Distribution Fit |
| --- | --- | --- | --- | --- | --- | --- | --- | --- | --- | --- | --- |
| IR | 25 C | 136 | 0.24 | 0.15 | 0.11 | 0.18 | 0.22 | 0.26 | 1.83 | 0.08 | LogNormal(0.0; 0.333; 0.227) |
|  | 34 C | 108 | 0.19 | 0.06 | 0.10 | 0.16 | 0.18 | 0.22 | 0.53 | 0.06 | LogNormal(0.0; 0.285; 0.185) |
| TC | 25 C | 136 | 16.19 | 5.95 | 5.61 | 12.27 | 15.04 | 19.85 | 41.43 | 7.58 | LogNormal(0.0; 0.355; 15.204) |
|  | 34 C | 108 | 12.44 | 4.45 | 4.40 | 8.84 | 11.83 | 15.25 | 24.40 | 6.40 | SkewedGaussian(6.645; 7.291; 4.928) |
| $V_{Rest}$ | 25 C | 136 | -60.91 | 7.06 | -77.95 | -66.04 | -61.41 | -56.09 | -39.62 | 9.95 | LogNormal(-142.87; 0.086; 81.661) |
|  | 34 C | 108 | -63.53 | 6.34 | -77.51 | -67.86 | -63.00 | -59.42 | -48.47 | 8.43 | SkewedGaussian(-59.434; 7.526; -0.932) |
| Gain current | 25 C | 135 | 0.37 | 0.14 | 0.02 | 0.27 | 0.34 | 0.42 | 1.01 | 0.16 | SkewedGaussian(0.212; 0.21; 3.041) |
|  | 34 C | 104 | 0.54 | 0.29 | 0.23 | 0.36 | 0.47 | 0.60 | 2.23 | 0.23 | LogNormal(0.0; 0.423; 0.484) |
| Gain voltage | 25 C | 135 | 1.66 | 0.76 | 0.04 | 1.23 | 1.49 | 1.92 | 5.82 | 0.69 | SkewedGaussian(0.871; 1.096; 3.21) |
|  | 34 C | 104 | 2.92 | 1.95 | 0.80 | 2.00 | 2.65 | 3.31 | 17.25 | 1.31 | LogNormal(0.0; 0.46; 2.583) |
| Threshold current | 25 C | 135 | 42.21 | 45.01 | -35.88 | 20.32 | 29.72 | 54.42 | 228.06 | 34.10 | SkewedGaussian(-6.002; 65.845; 3.661) |
|  | 34 C | 104 | 68.03 | 54.77 | -18.08 | 32.69 | 54.97 | 91.52 | 287.22 | 58.82 | SkewedGaussian(4.388; 83.792; 5.274) |
| $V_{\theta}^{Rel}$ | 25 C | 135 | 8.05 | 7.34 | -15.23 | 4.85 | 7.67 | 11.95 | 28.92 | 7.10 | SkewedGaussian(3.041; 8.865; 1.01) |
|  | 34 C | 104 | 11.90 | 7.46 | -3.22 | 7.03 | 11.84 | 16.41 | 37.07 | 9.38 | Gaussian(N(11.898; 7.422)) |
| $V_{\theta}$ | 25 C | 135 | -52.85 | 9.34 | -91.63 | -57.61 | -52.91 | -46.83 | -30.34 | 10.78 | SkewedGaussian(-45.131; 12.09; -1.362) |
|  | 34 C | 104 | -51.57 | 7.58 | -67.35 | -56.50 | -51.42 | -46.30 | -36.25 | 10.20 | LogNormal(-378.767; 0.023; 327.103) |
| Saturation | 25 C | 135 | 55.14 | 28.64 | 2.46 | 38.37 | 50.58 | 65.34 | 201.43 | 26.97 | LogNormal(0.0; 0.528; 48.729) |
|  | 34 C | 105 | 81.62 | 62.86 | 0.00 | 37.55 | 60.94 | 105.66 | 320.42 | 68.11 | SkewedGaussian(9.799; 95.25; 13.61) |
| SFA | 25 C | 136 | 0.28 | 0.13 | 0.04 | 0.17 | 0.28 | 0.35 | 0.64 | 0.18 | Gaussian(N(0.278; 0.129)) |
|  | 34 C | 108 | 0.35 | 0.23 | 0.00 | 0.15 | 0.33 | 0.49 | 0.88 | 0.34 | Gaussian(N(0.351; 0.226)) |

**Table 8:**
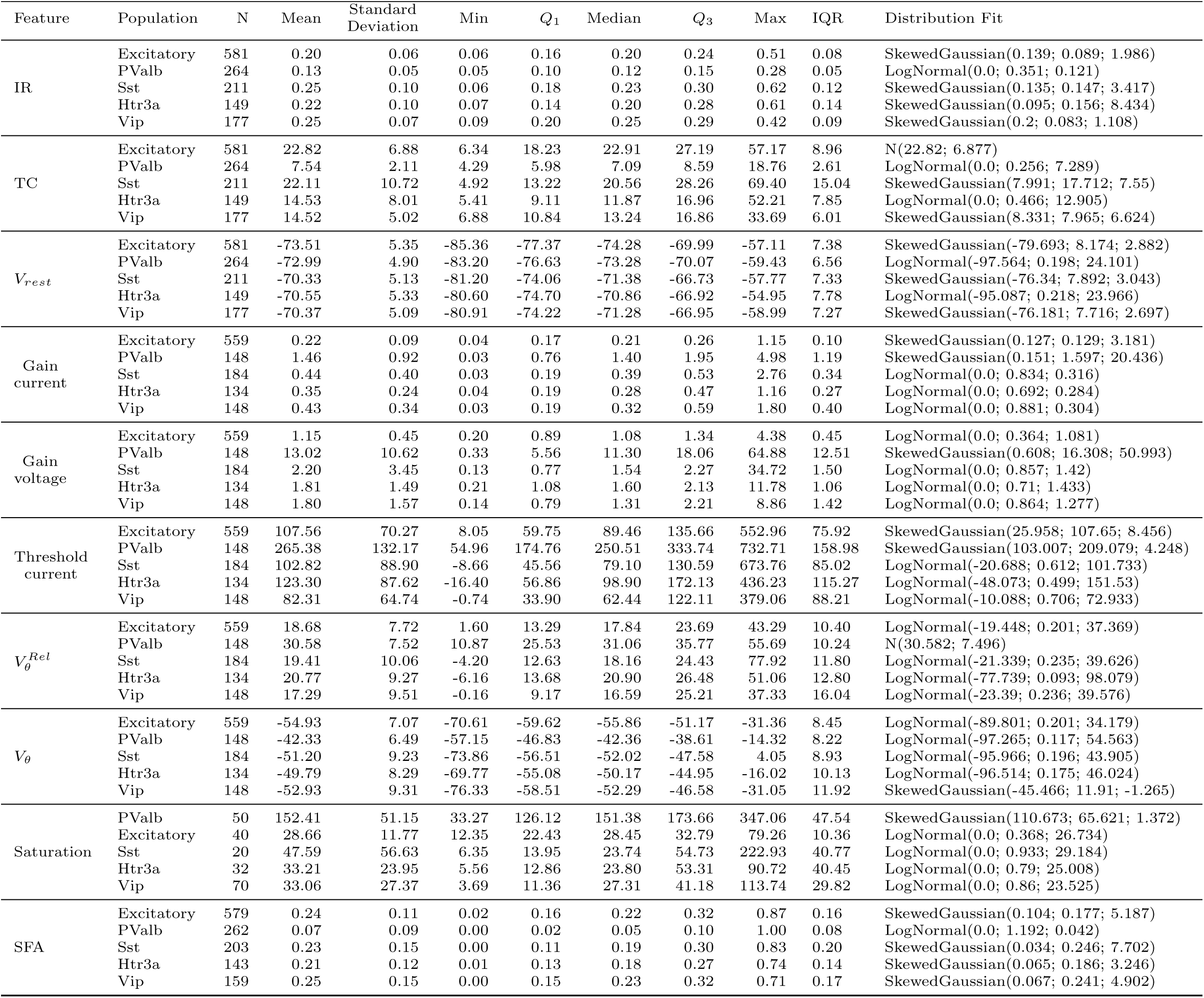
Table of descriptive statistics and fitting parameters for figure 5. **IR**: Input Resistance (*G*Ω),**TC**: Time Constant(ms), *V_rest_*: Resting membrane potential (mV), **Gain current** (Hz/pA),**Gain voltage**: (Hz/mV), **Threshold current**: (pA), *V ^Rel^*: Threshold Voltage (mV), *V_θ_*: Threshold Absolute Voltage (mV), **Saturation**: Saturation Frequency (Hz), **SFA**: Spike Frequency Adaptation (no unit)

**Table 9:** Table of descriptive statistics and fitting parameters for figure 6. IR. : Input Resistance (*G*Ω),**TC**: Time Constant(ms), *V_rest_*: Resting membrane potential (mV), **Gain current** (Hz/pA),Gain voltage: (Hz/mV), **Threshold current**: (pA), *V_θ_*:*^Rel^*: Threshold Voltage (mV), *V***_θ_**: Threshold Absolute Voltage (mV), **Saturation**: Saturation Frequency (Hz), **SFA**: Spike Frequency Adaptation (no unit)

| Feature | Population | N | Mean | Standard Deviation | Min | Q<sub>1</sub> | Median | Q<sub>3</sub> | Max | IQR | Distribution Fit |
| --- | --- | --- | --- | --- | --- | --- | --- | --- | --- | --- | --- |
| IR | PValb | 191 | 0.17 | 0.05 | 0.07 | 0.14 | 0.17 | 0.20 | 0.36 | 0.05 | LogNormal(0.0; 0.251; 0.167) |
|  | Vip | 137 | 0.29 | 0.16 | 0.08 | 0.20 | 0.25 | 0.32 | 1.35 | 0.13 | LogNormal(0.0; 0.403; 0.26) |
|  | Sst | 256 | 0.25 | 0.12 | 0.11 | 0.19 | 0.24 | 0.29 | 1.83 | 0.09 | LogNormal(0.0; 0.314; 0.238) |
|  | Excitatory | 25 | 0.20 | 0.06 | 0.12 | 0.16 | 0.18 | 0.22 | 0.35 | 0.07 | SkewedGaussian(0.125; 0.094; 9.759) |
| TC | PValb | 191 | 9.77 | 2.16 | 5.31 | 8.24 | 9.60 | 11.17 | 18.49 | 2.94 | SkewedGaussian(7.537; 3.104; 2.066) |
|  | Vip | 137 | 11.13 | 5.08 | 3.26 | 7.37 | 10.28 | 13.24 | 32.73 | 5.87 | LogNormal(0.0; 0.441; 10.107) |
|  | Sst | 256 | 16.81 | 6.66 | 5.61 | 12.27 | 15.60 | 20.47 | 42.06 | 8.19 | LogNormal(0.0; 0.386; 15.606) |
|  | Excitatory | 25 | 15.75 | 5.62 | 7.38 | 12.22 | 14.72 | 18.77 | 28.19 | 6.55 | SkewedGaussian(8.928; 8.765; 4.33) |
| $V_{rest}$ | PValb | 191 | -64.13 | 6.65 | -94.98 | -67.93 | -64.58 | -60.05 | -45.26 | 7.88 | SkewedGaussian(-58.991; 8.391; -1.216) |
|  | Vip | 137 | -62.24 | 8.02 | -86.65 | -67.38 | -61.64 | -56.81 | -40.61 | 10.57 | SkewedGaussian(-55.733; 10.3; -1.297) |
|  | Sst | 256 | -60.01 | 7.31 | -78.73 | -65.01 | -60.76 | -55.04 | -39.62 | 9.98 | SkewedGaussian(-65.737; 9.272; 1.224) |
|  | Excitatory | 25 | -64.58 | 6.78 | -75.93 | -70.37 | -64.28 | -59.97 | -50.51 | 10.40 | LogNormal(-97.652; 0.202; 32.411) |
| Gain current | PValb | 159 | 0.49 | 0.28 | 0.08 | 0.35 | 0.43 | 0.52 | 2.23 | 0.17 | LogNormal(0.0; 0.436; 0.439) |
|  | Vip | 136 | 0.50 | 0.27 | 0.05 | 0.30 | 0.49 | 0.63 | 1.84 | 0.33 | SkewedGaussian(0.188; 0.41; 3.369) |
|  | Sst | 254 | 0.36 | 0.15 | 0.02 | 0.27 | 0.33 | 0.42 | 1.04 | 0.15 | SkewedGaussian(0.206; 0.216; 2.912) |
|  | Excitatory | 25 | 0.23 | 0.16 | 0.04 | 0.12 | 0.17 | 0.28 | 0.62 | 0.16 | LogNormal(0.0; 0.711; 0.179) |
| Gain voltage | PValb | 159 | 2.91 | 1.49 | 0.57 | 2.13 | 2.67 | 3.15 | 11.82 | 1.02 | LogNormal(0.0; 0.395; 2.669) |
|  | Vip | 136 | 1.98 | 1.10 | 0.24 | 1.09 | 1.95 | 2.64 | 6.22 | 1.55 | N(1.984; 1.1) |
|  | Sst | 254 | 1.57 | 0.77 | 0.04 | 1.11 | 1.43 | 1.84 | 5.83 | 0.72 | LogNormal(0.0; 0.51; 1.404) |
|  | Excitatory | 25 | 1.14 | 0.74 | 0.23 | 0.63 | 0.81 | 1.49 | 2.76 | 0.86 | LogNormal(0.0; 0.647; 0.936) |
| Threshold current | PValb | 159 | 94.56 | 54.59 | -156.68 | 61.24 | 85.02 | 121.72 | 237.86 | 60.48 | SkewedGaussian(54.684; 67.468; 1.11) |
|  | Vip | 137 | 27.89 | 36.63 | -21.19 | 2.01 | 23.42 | 41.92 | 282.42 | 39.91 | SkewedGaussian(-17.648; 58.357; 15.64) |
|  | Sst | 254 | 40.05 | 43.58 | -35.88 | 17.53 | 28.27 | 52.99 | 242.22 | 35.46 | SkewedGaussian(-7.287; 64.285; 3.963) |
|  | Excitatory | 25 | 59.34 | 35.59 | -13.98 | 44.52 | 61.52 | 72.12 | 131.72 | 27.60 | SkewedGaussian(32.644; 43.917; 1.177) |
| $V_{\theta}^{Rel}$ | PValb | 159 | 14.92 | 6.94 | -21.81 | 10.19 | 15.07 | 19.66 | 31.07 | 9.47 | SkewedGaussian(21.458; 9.516; -1.749) |
|  | Vip | 137 | 5.64 | 6.89 | -12.70 | 0.51 | 5.59 | 10.95 | 22.46 | 10.44 | SkewedGaussian(11.114; 8.785; -1.247) |
|  | Sst | 254 | 8.05 | 7.66 | -16.06 | 4.55 | 7.25 | 12.15 | 32.44 | 7.60 | SkewedGaussian(1.737; 9.915; 1.348) |
|  | Excitatory | 25 | 10.25 | 5.72 | -4.94 | 7.68 | 9.83 | 13.63 | 19.56 | 5.96 | SkewedGaussian(16.31; 8.256; -2.507) |
| $V_{\theta}$ | PValb | 159 | -48.88 | 7.81 | -86.16 | -53.58 | -48.49 | -44.00 | -25.97 | 9.59 | SkewedGaussian(-42.155; 10.288; -1.466) |
|  | Vip | 137 | -56.60 | 8.98 | -79.39 | -63.41 | -56.83 | -50.10 | -31.24 | 13.31 | N(-56.601; 8.949) |
|  | Sst | 254 | -51.93 | 8.84 | -91.63 | -56.86 | -51.92 | -46.16 | -30.34 | 10.70 | SkewedGaussian(-44.345; 11.641; -1.448) |
|  | Excitatory | 25 | -54.33 | 5.91 | -66.22 | -58.02 | -54.03 | -50.81 | -43.21 | 7.21 | SkewedGaussian(-51.586; 6.405; -0.636) |
| Saturation | PValb | 157 | 119.80 | 34.82 | 24.36 | 94.76 | 117.38 | 143.19 | 212.65 | 48.44 | LogNormal(0.0; 0.309; 114.569) |
|  | Vip | 137 | 48.12 | 27.11 | 8.05 | 31.73 | 41.29 | 54.15 | 177.03 | 22.43 | LogNormal(0.0; 0.522; 42.042) |
|  | Sst | 254 | 52.39 | 27.03 | 2.46 | 35.84 | 47.04 | 60.47 | 201.43 | 24.63 | LogNormal(0.0; 0.507; 46.488) |
|  | Excitatory | 25 | 38.61 | 32.02 | 18.80 | 24.91 | 30.36 | 33.70 | 177.76 | 8.79 | LogNormal(0.0; 0.466; 33.126) |
| SFA | PValb | 191 | 0.09 | 0.05 | 0.00 | 0.05 | 0.08 | 0.11 | 0.32 | 0.06 | SkewedGaussian(0.022; 0.082; 4.621) |
|  | Vip | 137 | 0.28 | 0.15 | 0.03 | 0.17 | 0.25 | 0.35 | 0.84 | 0.18 | SkewedGaussian(0.084; 0.246; 6.004) |
|  | Sst | 256 | 0.29 | 0.14 | 0.01 | 0.18 | 0.29 | 0.36 | 0.75 | 0.18 | SkewedGaussian(0.156; 0.19; 1.718) |
|  | Excitatory | 25 | 0.30 | 0.12 | 0.06 | 0.21 | 0.29 | 0.35 | 0.63 | 0.14 | SkewedGaussian(0.181; 0.168; 1.787) |

**Table 10:**
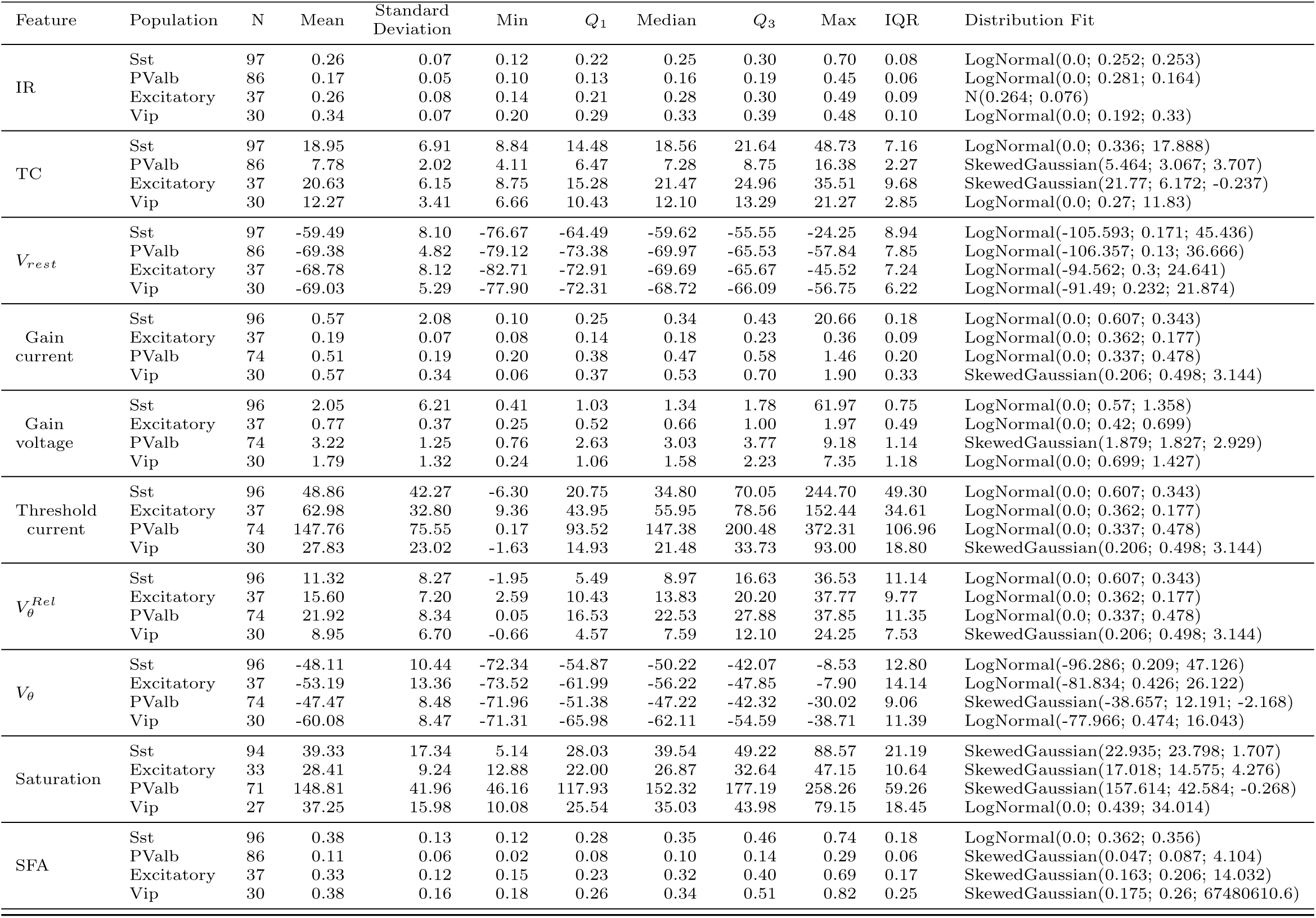
Table of descriptive statistics and fitting parameters for figure 7. IR: Input Resistance (*G*Ω),**TC**: Time Constant(ms), *V_rest_*: Resting membrane potential (mV), **Gain current** (Hz/pA),Gain voltage: (Hz/mV), **Threshold current**: (pA), *V_θ_*:*^Rel^* : Threshold Voltage (mV), *V_θ_*:*^Rel^* : Threshold Absolute Voltage (mV), **Saturation**: Saturation Frequency (Hz), **SFA**: Spike Frequency Adaptation (no unit)

**Table 11:** Table of descriptive statistics and fitting parameters for figure 11. **IR**: Input Resistance (*G*Ω),**TC**: Time Constant(ms), *V_rest_*: Resting membrane potential (mV), **Gain current** (Hz/pA),**Gain voltage**: (Hz/mV), **Threshold current**: (pA), *V ^Rel^*: Threshold Voltage (mV), *V_θ_*: Threshold Absolute Voltage (mV), **Saturation**: Saturation Frequency (Hz), **SFA**: Spike Frequency Adaptation (no unit)

| Feature | Population | N | Mean | Standard Deviation | Min | $Q_1$ | Median | $Q_3$ | Max | IQR | Distribution Fit |
| --- | --- | --- | --- | --- | --- | --- | --- | --- | --- | --- | --- |
| IR | VIS | 122 | 0.26 | 0.09 | 0.10 | 0.20 | 0.24 | 0.31 | 0.51 | 0.12 | SkewedGaussian(0.149; 0.142; 4.015) |
|  | MO | 108 | 0.19 | 0.06 | 0.10 | 0.16 | 0.18 | 0.22 | 0.53 | 0.06 | LogNormal(0.0; 0.285; 0.185) |
| TC | VIS | 122 | 24.11 | 9.94 | 7.53 | 16.75 | 22.76 | 29.85 | 59.51 | 13.10 | LogNormal(0.0; 0.422; 22.13) |
|  | MO | 108 | 12.44 | 4.45 | 4.40 | 8.84 | 11.83 | 15.25 | 24.40 | 6.40 | SkewedGaussian(6.645; 7.291; 4.928) |
| $V_{rest}$ | VIS | 122 | -70.51 | 5.08 | -80.75 | -74.46 | -71.54 | -67.06 | -57.77 | 7.40 | SkewedGaussian(-76.808; 8.076; 4.596) |
|  | MO | 108 | -63.53 | 6.34 | -77.51 | -67.86 | -63.00 | -59.42 | -48.47 | 8.43 | SkewedGaussian(-59.434; 7.526; -0.932) |
| Gain current | VIS | 110 | 0.39 | 0.34 | 0.03 | 0.17 | 0.36 | 0.52 | 2.35 | 0.35 | LogNormal(0.0; 0.827; 0.288) |
|  | MO | 104 | 0.54 | 0.29 | 0.23 | 0.36 | 0.47 | 0.60 | 2.23 | 0.23 | LogNormal(0.0; 0.423; 0.484) |
| Gain voltage | VIS | 110 | 1.69 | 1.79 | 0.13 | 0.68 | 1.34 | 1.90 | 12.17 | 1.22 | LogNormal(0.0; 0.814; 1.207) |
|  | MO | 104 | 2.92 | 1.95 | 0.80 | 2.00 | 2.65 | 3.31 | 17.25 | 1.31 | LogNormal(0.0; 0.46; 2.583) |
| Threshold current | VIS | 110 | 89.45 | 66.11 | -8.66 | 43.16 | 73.82 | 106.93 | 387.06 | 63.77 | SkewedGaussian(14.758; 99.545; 7.936) |
|  | MO | 104 | 68.03 | 54.77 | -18.08 | 32.69 | 54.97 | 91.52 | 287.22 | 58.82 | SkewedGaussian(4.388; 83.792; 5.274) |
| $V_{\theta}^{Rel}$ | VIS | 110 | 19.01 | 10.74 | -4.20 | 12.48 | 17.30 | 23.71 | 77.92 | 11.24 | LogNormal(-17.752; 0.268; 35.428) |
|  | MO | 104 | 11.90 | 7.46 | -3.22 | 7.03 | 11.84 | 16.41 | 37.07 | 9.38 | N(11.898; 7.422) |
| $V_{\theta}$ | VIS | 110 | -51.76 | 10.05 | -72.55 | -57.81 | -52.50 | -48.30 | 4.05 | 9.51 | LogNormal(-85.088; 0.274; 32.064) |
|  | MO | 104 | -51.57 | 7.58 | -67.35 | -56.50 | -51.42 | -46.30 | -36.25 | 10.20 | LogNormal(-378.767; 0.023; 327.103) |
| Saturation | VIS | 8 | 37.84 | 44.32 | 11.40 | 12.53 | 17.51 | 38.05 | 138.84 | 25.52 | N(37.835; 41.461) |
|  | MO | 105 | 81.62 | 62.86 | 0.00 | 37.55 | 60.94 | 105.66 | 320.42 | 68.11 | SkewedGaussian(9.799; 95.25; 13.61) |
| SFA | VIS | 118 | 0.24 | 0.16 | 0.01 | 0.13 | 0.22 | 0.32 | 0.83 | 0.19 | SkewedGaussian(0.04; 0.254; 8.028) |
|  | MO | 108 | 0.35 | 0.23 | 0.00 | 0.15 | 0.33 | 0.49 | 0.88 | 0.34 | N(0.351; 0.226) |

**Table 12:** Table of descriptive statistics and fitting parameters for figure 12. **IR**: Input Resistance (*G*Ω),**TC**: Time Constant(ms), *V_rest_*: Resting membrane potential (mV), **Gain current** (Hz/pA),**Gain voltage**: (Hz/mV), **Threshold current**: (pA), *V ^Rel^*: Threshold Voltage (mV), *V_θ_*: Threshold Absolute Voltage (mV), **Saturation**: Saturation Frequency (Hz), **SFA**: Spike Frequency Adaptation (no unit)

| Feature | Population | N | Mean | Standard Deviation | Min | $Q_1$ | Median | $Q_3$ | Max | IQR | Distribution Fit |
| --- | --- | --- | --- | --- | --- | --- | --- | --- | --- | --- | --- |
| IR | VIS | 80 | 0.27 | 0.08 | 0.12 | 0.22 | 0.26 | 0.30 | 0.70 | 0.08 | LogNormal(0.0; 0.262; 0.257) |
|  | SS | 58 | 0.19 | 0.05 | 0.10 | 0.16 | 0.18 | 0.22 | 0.33 | 0.07 | SkewedGaussian(0.141; 0.067; 2.428) |
| TC | VIS | 80 | 19.32 | 7.21 | 8.84 | 14.70 | 19.02 | 22.05 | 48.73 | 7.35 | LogNormal(0.0; 0.341; 18.193) |
|  | SS | 58 | 11.12 | 3.16 | 4.49 | 9.08 | 10.94 | 12.62 | 21.55 | 3.54 | SkewedGaussian(7.99; 4.43; 1.977) |
| $V_{rest}$ | VIS | 80 | -59.72 | 8.41 | -76.67 | -64.57 | -60.40 | -55.57 | -24.25 | 9.00 | LogNormal(-103.437; 0.186; 42.965) |
|  | SS | 58 | -59.22 | 6.56 | -77.37 | -62.60 | -59.06 | -54.34 | -41.50 | 8.26 | SkewedGaussian(-53.177; 8.883; -1.67) |
| Gain current | VIS | 79 | 0.60 | 2.29 | 0.10 | 0.25 | 0.32 | 0.42 | 20.66 | 0.16 | LogNormal(0.0; 0.621; 0.336) |
|  | SS | 58 | 0.38 | 0.13 | 0.19 | 0.30 | 0.37 | 0.44 | 0.83 | 0.14 | LogNormal(0.0; 0.311; 0.363) |
| Gain voltage | VIS | 79 | 2.09 | 6.84 | 0.46 | 0.98 | 1.28 | 1.64 | 61.97 | 0.66 | LogNormal(0.0; 0.565; 1.306) |
|  | SS | 58 | 2.08 | 0.87 | 1.11 | 1.63 | 1.87 | 2.28 | 6.40 | 0.64 | LogNormal(0.0; 0.315; 1.962) |
| Threshold current | VIS | 79 | 52.33 | 44.92 | -6.30 | 20.55 | 37.30 | 76.08 | 244.70 | 55.53 | SkewedGaussian(-1.721; 70.098; 17.001) |
|  | SS | 58 | 53.37 | 53.22 | -8.40 | 19.58 | 33.71 | 75.27 | 216.20 | 55.69 | SkewedGaussian(-4.674; 78.433; 14.278) |
| $V_{\theta}^{Rel}$ | VIS | 79 | 12.01 | 8.53 | -1.95 | 5.52 | 10.10 | 17.72 | 36.53 | 12.20 | SkewedGaussian(0.914; 13.967; 6.662) |
|  | SS | 58 | 8.69 | 6.95 | -1.40 | 3.98 | 6.22 | 14.00 | 27.23 | 10.01 | LogNormal(-5.247; 0.504; 12.325) |
| $V_{\theta}$ | VIS | 79 | -47.63 | 10.86 | -72.34 | -54.90 | -49.88 | -41.86 | -8.53 | 13.03 | LogNormal(-99.157; 0.203; 50.464) |
|  | SS | 58 | -50.53 | 7.23 | -63.35 | -55.47 | -51.68 | -46.22 | -25.68 | 9.25 | SkewedGaussian(-58.593; 10.786; 2.791) |
| Saturation | VIS | 77 | 39.50 | 16.06 | 5.14 | 28.30 | 37.41 | 49.10 | 88.57 | 20.80 | SkewedGaussian(22.972; 22.972; 2.085) |
|  | SS | 56 | 79.20 | 27.72 | 18.54 | 60.58 | 78.70 | 91.90 | 151.07 | 31.31 | SkewedGaussian(50.993; 39.371; 2.051) |
| SFA | VIS | 79 | 0.40 | 0.13 | 0.18 | 0.30 | 0.36 | 0.49 | 0.74 | 0.18 | SkewedGaussian(0.227; 0.219; 5.708) |
|  | SS | 58 | 0.27 | 0.09 | 0.06 | 0.22 | 0.26 | 0.32 | 0.62 | 0.10 | SkewedGaussian(0.18; 0.127; 1.702) |

**Table 13:**
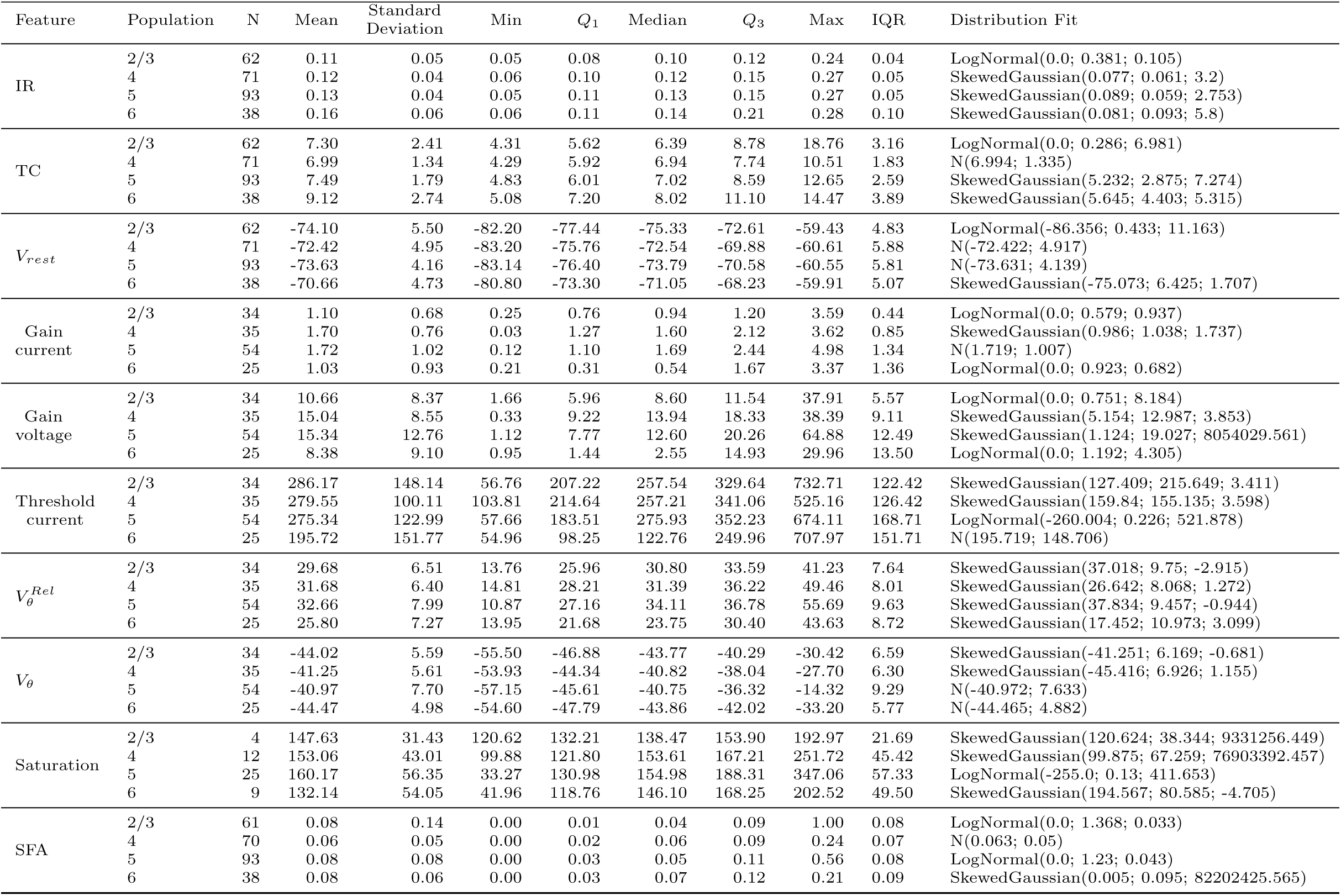
Table of descriptive statistics and fitting parameters for figure 13. IR: Input Resistance (*G*Ω),TC: Time Constant(ms), Vrest: Resting membrane potential (mV), **Gain current** (Hz/pA),Gain voltage: (Hz/mV), **Threshold current**: (pA), *V_θ_*:*^Rel^* : Threshold Voltage (mV), *V_θ_*: Threshold Absolute Voltage (mV), **Saturation**: Saturation Frequency (Hz), **SFA**: Spike Frequency Adaptation (no unit)

**Table 14:**
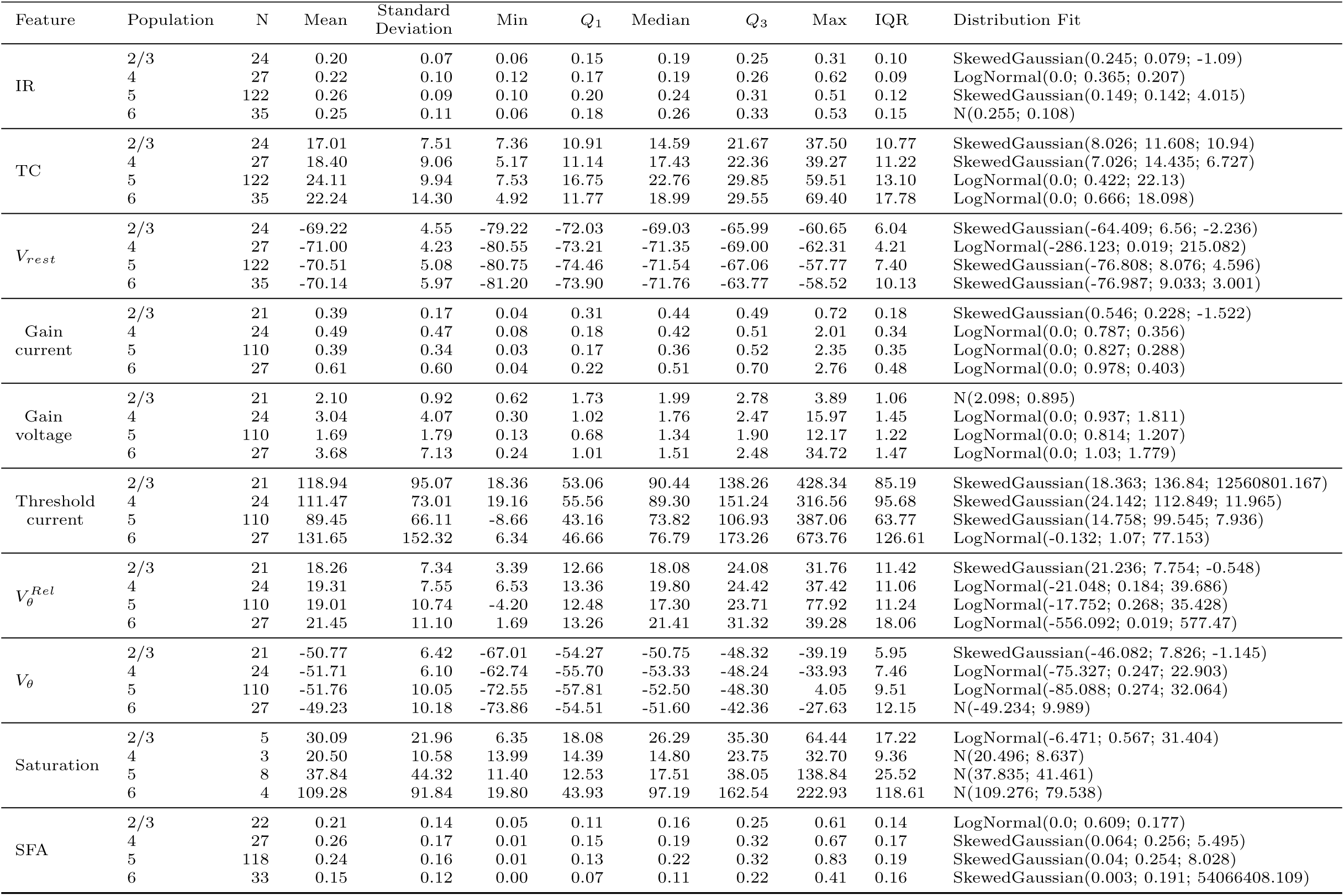
Table of descriptive statistics and fitting parameters for figure 14. IR. : Input Resistance (*G*Ω),TC: Time Constant(ms), Vrest: Resting membrane potential (mV), **Gain current** (Hz/pA),Gain voltage: (Hz/mV), **Threshold current**: (pA), *V_θ_*:*^Rel^* : Threshold Voltage (mV), *V_θ_*: Threshold Absolute Voltage (mV), **Saturation**: Saturation Frequency (Hz), **SFA**: Spike Frequency Adaptation (no unit)

**Table 15:**
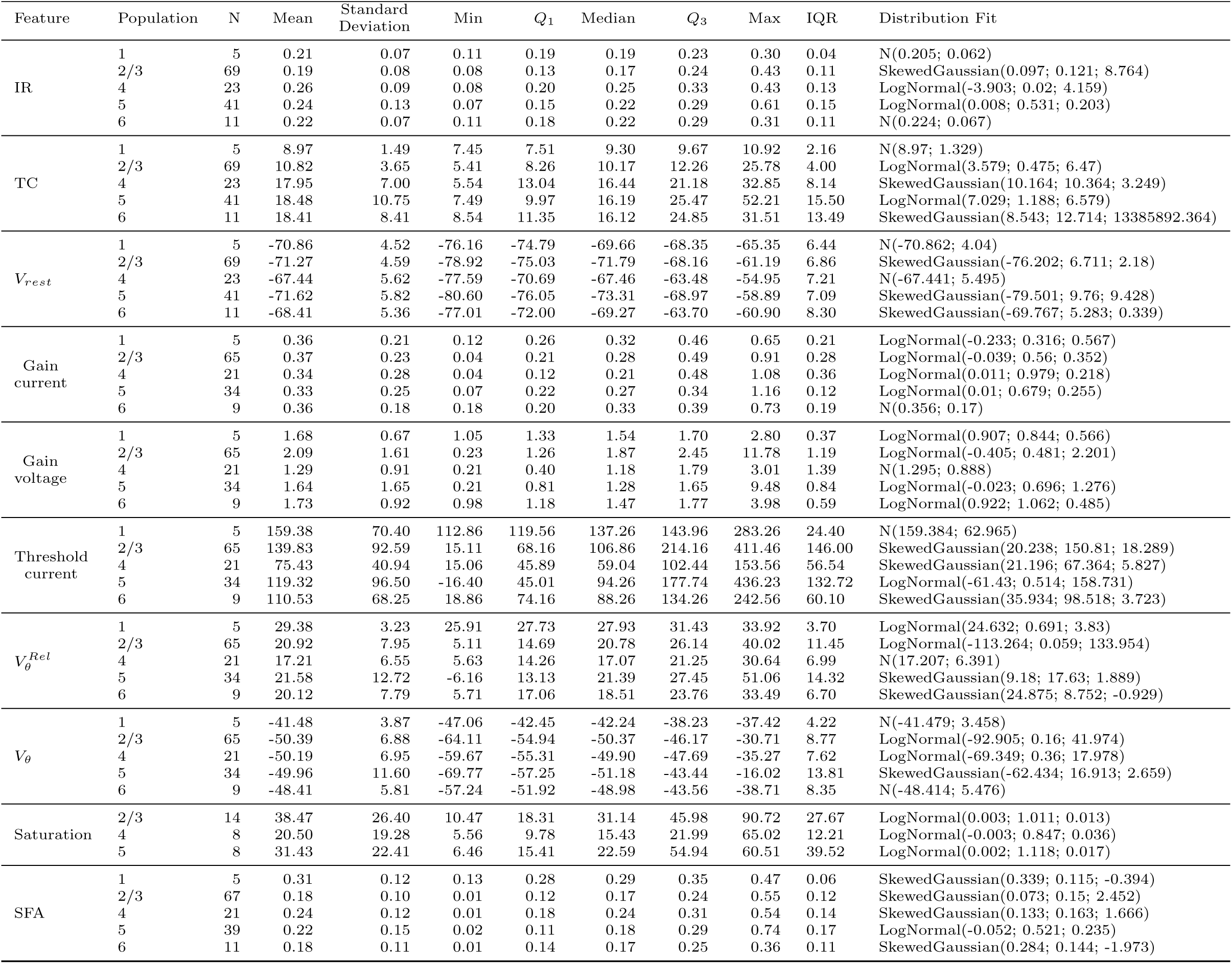
Table of descriptive statistics and fitting parameters for figure 15. **IR**: Input Resistance (*G*Ω),**TC**: Time Constant(ms), *V_rest_*: Resting membrane potential (mV), **Gain current** (Hz/pA),**Gain voltage**: (Hz/mV), **Threshold current**: (pA), *V ^Rel^*: Threshold Voltage (mV), *V_θ_*: Threshold Absolute Voltage (mV), **Saturation**: Saturation Frequency (Hz), **SFA**: Spike Frequency Adaptation (no unit)

**Table 16:**
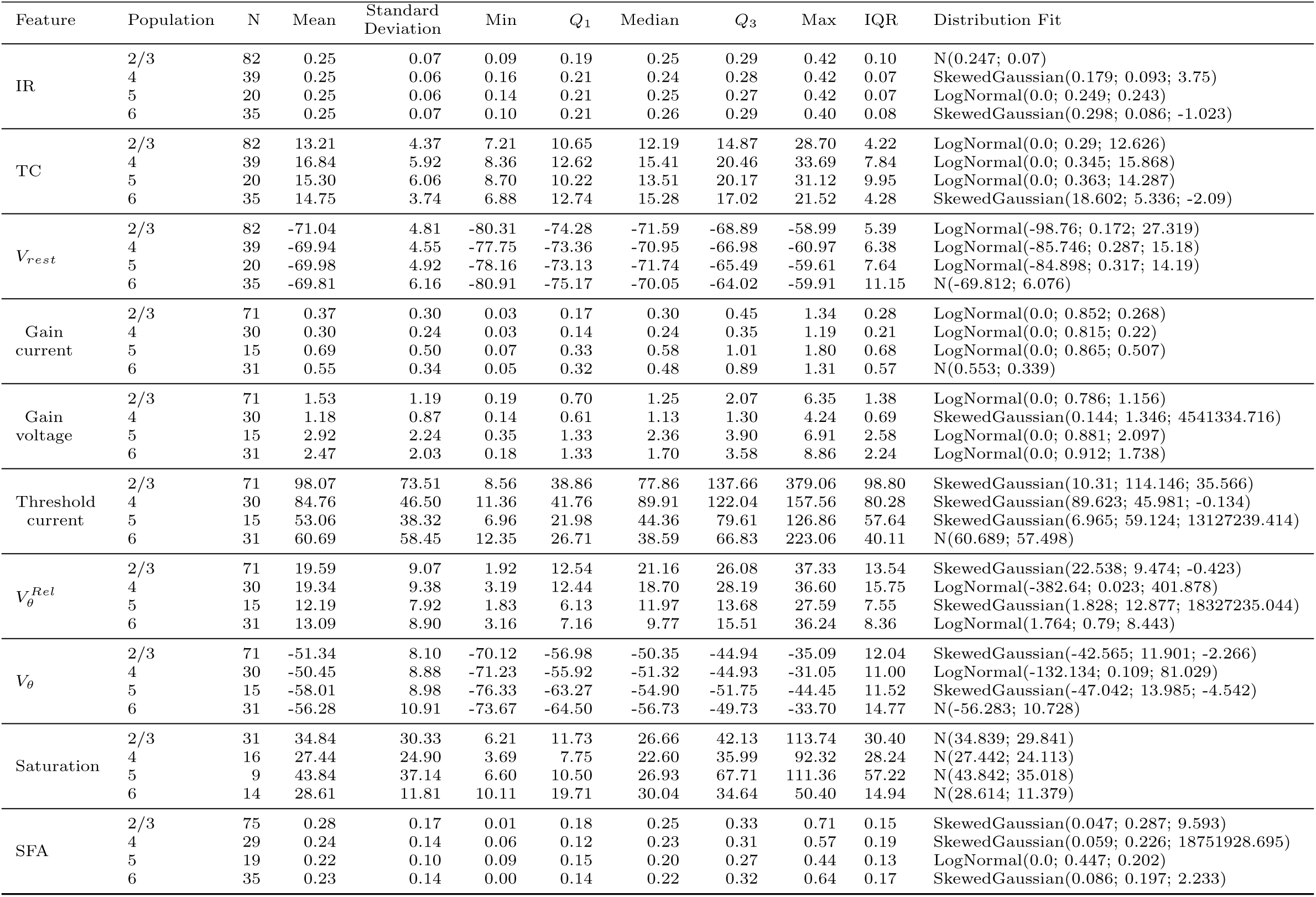
Table of descriptive statistics and fitting parameters for figure 16. IR: Input Resistance (*G*Ω),TC: Time Constant(ms), Vrest: Resting membrane potential (mV), **Gain current** (Hz/pA),Gain voltage: (Hz/mV), **Threshold current**: (pA), *V_θ_*:*^Rel^* : Threshold Voltage (mV), *V_θ_*: Threshold Absolute Voltage (mV), **Saturation**: Saturation Frequency (Hz), **SFA**: Spike Frequency Adaptation (no unit)

**Table 17:**
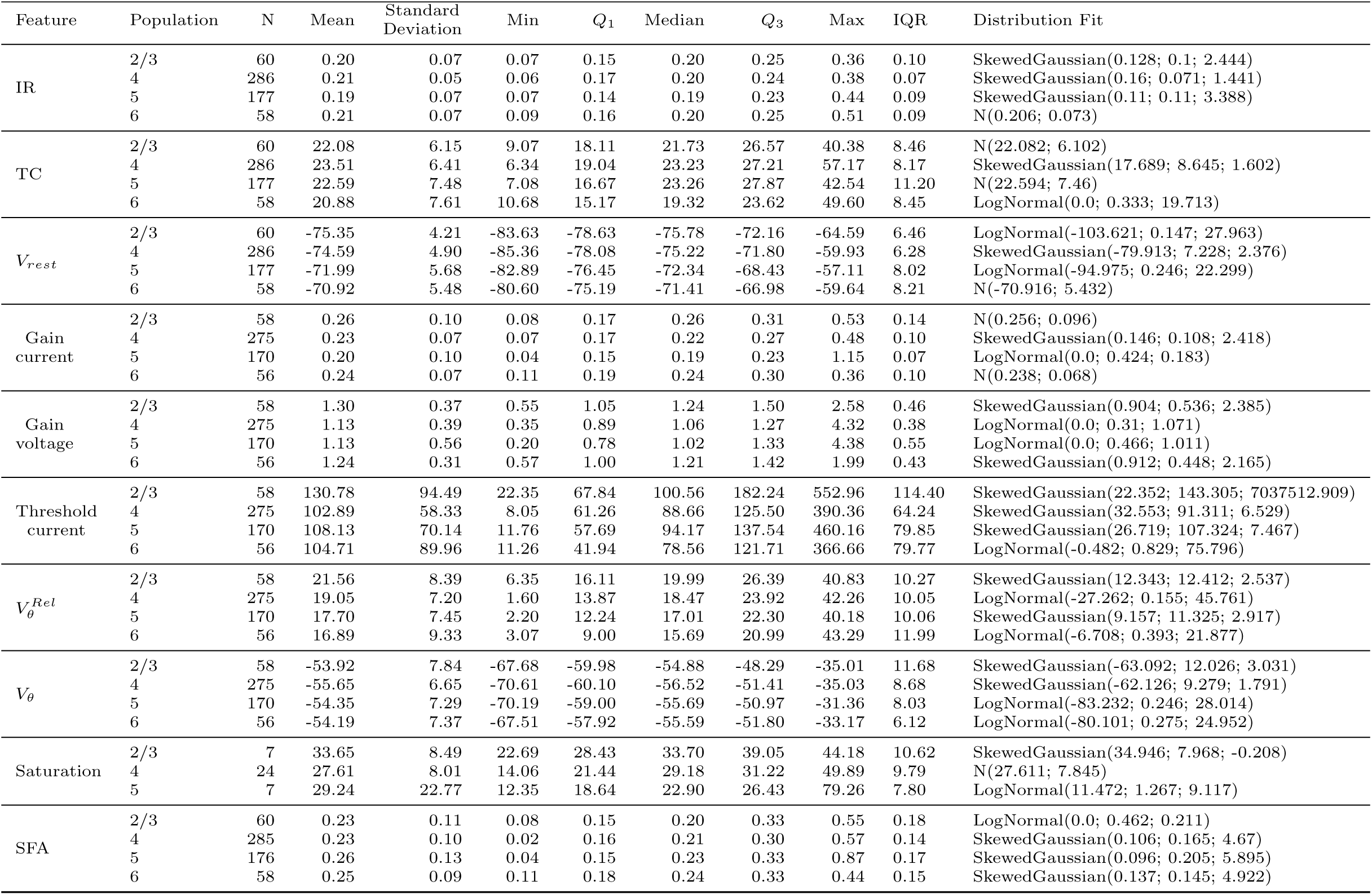
Table of descriptive statistics and fitting parameters for figure 17. IR. : Input Resistance (GΩ),TC: Time Constant(ms), Vrest: Resting membrane potential (mV), **Gain current** (Hz/pA),Gain voltage: (Hz/mV), **Threshold current**: (pA), *V_θ_*:*^Rel^* : Threshold Voltage (mV), *V_θ_*: Threshold Absolute Voltage (mV), **Saturation**: Saturation Frequency (Hz), **SFA**: Spike Frequency Adaptation (no unit)

**Table 18:** Table of descriptive statistics and fitting parameters for figure 18. **IR**: Input Resistance (*G*Ω),**TC**: Time Constant(ms), *V_rest_*: Resting membrane potential (mV), **Gain current** (Hz/pA),**Gain voltage**: (Hz/mV), **Threshold current**: (pA), *V ^Rel^*: Threshold Voltage (mV), *V_θ_*: Threshold Absolute Voltage (mV), **Saturation**: Saturation Frequency (Hz), **SFA**: Spike Frequency Adaptation (no unit)

| Feature | Population | N | Mean | Standard Deviation | Min | Q<sub>1</sub> | Median | Q<sub>3</sub> | Max | IQR | Distribution Fit |
| --- | --- | --- | --- | --- | --- | --- | --- | --- | --- | --- | --- |
| IR | 2/3 | 75 | 0.27 | 0.07 | 0.13 | 0.22 | 0.27 | 0.30 | 0.53 | 0.08 | N(0.266; 0.072) |
|  | 5 | 136 | 0.24 | 0.15 | 0.11 | 0.18 | 0.22 | 0.26 | 1.83 | 0.08 | LogNormal(0.0; 0.333; 0.227) |
|  | 6 | 45 | 0.25 | 0.08 | 0.12 | 0.19 | 0.24 | 0.30 | 0.45 | 0.10 | SkewedGaussian(0.159; 0.119; 3.844) |
| TC | 2/3 | 75 | 19.95 | 7.28 | 8.07 | 15.08 | 18.09 | 24.29 | 42.06 | 9.21 | LogNormal(0.0; 0.347; 18.773) |
|  | 5 | 136 | 16.19 | 5.95 | 5.61 | 12.27 | 15.04 | 19.85 | 41.43 | 7.58 | LogNormal(0.0; 0.355; 15.204) |
|  | 6 | 45 | 13.41 | 5.48 | 5.85 | 9.51 | 11.37 | 17.21 | 26.27 | 7.70 | SkewedGaussian(6.468; 8.806; 11.484) |
| $V_{rest}$ | 2/3 | 75 | -57.56 | 7.38 | -78.73 | -63.12 | -58.53 | -52.13 | -40.22 | 11.00 | LogNormal(-134.062; 0.095; 76.106) |
|  | 5 | 136 | -60.91 | 7.06 | -77.95 | -66.04 | -61.41 | -56.09 | -39.62 | 9.95 | LogNormal(-142.87; 0.086; 81.661) |
|  | 6 | 45 | -61.38 | 7.08 | -77.91 | -64.91 | -61.53 | -56.58 | -45.39 | 8.32 | LogNormal(-200.136; 0.05; 138.634) |
| Gain current | 2/3 | 75 | 0.32 | 0.10 | 0.05 | 0.25 | 0.29 | 0.37 | 0.64 | 0.12 | SkewedGaussian(0.218; 0.14; 2.245) |
|  | 5 | 135 | 0.37 | 0.14 | 0.02 | 0.27 | 0.34 | 0.42 | 1.01 | 0.16 | SkewedGaussian(0.212; 0.21; 3.041) |
|  | 6 | 44 | 0.43 | 0.21 | 0.05 | 0.32 | 0.38 | 0.52 | 1.04 | 0.20 | SkewedGaussian(0.215; 0.295; 2.469) |
| Gain voltage | 2/3 | 75 | 1.25 | 0.42 | 0.37 | 0.92 | 1.21 | 1.56 | 2.29 | 0.64 | SkewedGaussian(0.778; 0.63; 2.688) |
|  | 5 | 135 | 1.66 | 0.76 | 0.04 | 1.23 | 1.49 | 1.92 | 5.82 | 0.69 | SkewedGaussian(0.871; 1.096; 3.21) |
|  | 6 | 44 | 1.81 | 1.05 | 0.23 | 1.23 | 1.49 | 2.10 | 5.83 | 0.87 | LogNormal(0.0; 0.578; 1.553) |
| Threshold current | 2/3 | 75 | 26.98 | 38.67 | -30.48 | 2.06 | 22.02 | 41.92 | 242.22 | 39.86 | SkewedGaussian(-15.771; 57.473; 5.418) |
|  | 5 | 135 | 42.21 | 45.01 | -35.88 | 20.32 | 29.72 | 54.42 | 228.06 | 34.10 | SkewedGaussian(-6.002; 65.845; 3.661) |
|  | 6 | 44 | 55.71 | 41.51 | 9.62 | 23.77 | 46.57 | 65.42 | 162.32 | 41.65 | LogNormal(2.522; 0.792; 39.667) |
| $V_{\theta}^{Rel}$ | 2/3 | 75 | 5.40 | 6.96 | -16.06 | 0.47 | 6.31 | 9.03 | 30.93 | 8.56 | N(5.403; 6.915) |
|  | 5 | 135 | 8.05 | 7.34 | -15.23 | 4.85 | 7.67 | 11.95 | 28.92 | 7.10 | SkewedGaussian(3.041; 8.865; 1.01) |
|  | 6 | 44 | 12.53 | 7.84 | 1.89 | 7.00 | 10.09 | 15.54 | 32.44 | 8.55 | LogNormal(-1.84; 0.533; 12.506) |
| $V_{\theta}$ | 2/3 | 75 | -52.16 | 7.89 | -83.21 | -56.57 | -51.67 | -47.64 | -32.15 | 8.93 | SkewedGaussian(-45.922; 10.012; -1.274) |
|  | 5 | 135 | -52.85 | 9.34 | -91.63 | -57.61 | -52.91 | -46.83 | -30.34 | 10.78 | SkewedGaussian(-45.131; 12.09; -1.362) |
|  | 6 | 44 | -48.75 | 8.26 | -73.91 | -53.22 | -48.15 | -43.20 | -33.91 | 10.03 | SkewedGaussian(-40.375; 11.694; -2.102) |
| Saturation | 2/3 | 75 | 45.10 | 18.46 | 15.58 | 34.95 | 43.54 | 51.30 | 147.60 | 16.36 | LogNormal(-3.29; 0.336; 45.645) |
|  | 5 | 135 | 55.14 | 28.64 | 2.46 | 38.37 | 50.58 | 65.34 | 201.43 | 26.97 | SkewedGaussian(23.886; 42.316; 4.204) |
|  | 6 | 44 | 56.40 | 32.00 | 10.43 | 32.60 | 49.35 | 77.52 | 143.62 | 44.92 | LogNormal(-14.833; 0.445; 64.636) |
| SFA | 2/3 | 75 | 0.35 | 0.13 | 0.02 | 0.28 | 0.33 | 0.40 | 0.72 | 0.12 | SkewedGaussian(0.238; 0.165; 1.45) |
|  | 5 | 136 | 0.28 | 0.13 | 0.04 | 0.17 | 0.28 | 0.35 | 0.64 | 0.18 | N(0.278; 0.129) |
|  | 6 | 45 | 0.22 | 0.15 | 0.01 | 0.12 | 0.21 | 0.29 | 0.75 | 0.17 | SkewedGaussian(0.037; 0.234; 7.183) |

**Table 19:** Table of descriptive statistics and fitting parameters for figure 19. **IR**: Input Resistance (*G*Ω),**TC**: Time Constant(ms), *V_rest_*: Resting membrane potential (mV), **Gain current** (Hz/pA),**Gain voltage**: (Hz/mV), **Threshold current**: (pA), *V ^Rel^*: Threshold Voltage (mV), *V_θ_*: Threshold Absolute Voltage (mV), **Saturation**: Saturation Frequency (Hz), **SFA**: Spike Frequency Adaptation (no unit)

| Feature | Population | N | Mean | Standard Deviation | Min | Q<sub>1</sub> | Median | Q<sub>3</sub> | Max | IQR | Distribution Fit |
| --- | --- | --- | --- | --- | --- | --- | --- | --- | --- | --- | --- |
| IR | 2/3 | 59 | 0.16 | 0.03 | 0.09 | 0.14 | 0.16 | 0.18 | 0.24 | 0.04 | SkewedGaussian(0.179; 0.038; -0.782) |
|  | 5 | 105 | 0.18 | 0.05 | 0.10 | 0.15 | 0.17 | 0.20 | 0.35 | 0.05 | LogNormal(0.0; 0.238; 0.173) |
|  | 6 | 27 | 0.18 | 0.06 | 0.07 | 0.14 | 0.18 | 0.20 | 0.36 | 0.06 | LogNormal(0.0; 0.324; 0.171) |
| TC | 2/3 | 59 | 10.07 | 2.44 | 5.61 | 8.49 | 9.81 | 11.58 | 18.49 | 3.09 | LogNormal(0.0; 0.237; 9.795) |
|  | 5 | 105 | 9.85 | 2.01 | 5.31 | 8.42 | 9.73 | 11.39 | 13.91 | 2.97 | LogNormal(0.0; 0.209; 9.641) |
|  | 6 | 27 | 8.83 | 1.87 | 5.98 | 7.45 | 8.67 | 10.33 | 12.83 | 2.88 | N(8.826; 1.837) |
| $V_{rest}$ | 2/3 | 59 | -63.17 | 5.82 | -77.22 | -67.62 | -63.23 | -59.63 | -50.92 | 7.98 | SkewedGaussian(-68.433; 7.811; 1.559) |
|  | 5 | 105 | -64.01 | 7.00 | -94.98 | -67.52 | -64.57 | -58.85 | -45.26 | 8.68 | SkewedGaussian(-58.013; 9.187; -1.478) |
|  | 6 | 27 | -66.69 | 6.56 | -82.00 | -69.99 | -67.06 | -63.21 | -52.27 | 6.77 | SkewedGaussian(-63.705; 7.097; -0.621) |
| Gain current | 2/3 | 52 | 0.39 | 0.11 | 0.08 | 0.31 | 0.40 | 0.45 | 0.68 | 0.13 | N(0.387; 0.106) |
|  | 5 | 84 | 0.52 | 0.30 | 0.20 | 0.39 | 0.44 | 0.53 | 2.23 | 0.14 | LogNormal(0.0; 0.384; 0.475) |
|  | 6 | 23 | 0.60 | 0.40 | 0.08 | 0.36 | 0.43 | 0.79 | 1.71 | 0.43 | LogNormal(0.0; 0.657; 0.487) |
| Gain voltage | 2/3 | 52 | 2.49 | 0.75 | 0.86 | 2.10 | 2.37 | 2.83 | 5.29 | 0.73 | SkewedGaussian(1.754; 1.048; 2.076) |
|  | 5 | 84 | 3.06 | 1.73 | 1.08 | 2.17 | 2.75 | 3.16 | 11.82 | 0.99 | LogNormal(0.0; 0.392; 2.787) |
|  | 6 | 23 | 3.30 | 1.62 | 0.57 | 2.35 | 2.94 | 4.10 | 8.48 | 1.75 | SkewedGaussian(1.555; 2.354; 3.343) |
| Threshold current | 2/3 | 52 | 100.91 | 58.60 | -17.14 | 60.24 | 87.52 | 131.04 | 237.86 | 70.79 | SkewedGaussian(33.472; 88.968; 3.25) |
|  | 5 | 84 | 91.60 | 44.62 | 13.96 | 61.44 | 82.09 | 112.10 | 216.46 | 50.67 | SkewedGaussian(40.89; 67.368; 3.096) |
|  | 6 | 23 | 91.03 | 76.11 | -156.68 | 45.02 | 97.72 | 140.97 | 222.36 | 95.95 | SkewedGaussian(170.91; 109.188; -2.728) |
| $V_{\theta}^{Rel}$ | 2/3 | 52 | 14.78 | 6.39 | -2.67 | 11.50 | 14.95 | 19.31 | 27.93 | 7.81 | SkewedGaussian(20.524; 8.553; -1.56) |
|  | 5 | 84 | 15.26 | 6.38 | 1.73 | 10.04 | 14.87 | 20.06 | 31.07 | 10.02 | SkewedGaussian(9.592; 8.509; 1.498) |
|  | 6 | 23 | 13.99 | 9.78 | -21.81 | 9.29 | 17.03 | 18.92 | 28.50 | 9.62 | SkewedGaussian(23.885; 13.762; -3.951) |
| $V_{\theta}$ | 2/3 | 52 | -48.34 | 7.30 | -65.26 | -52.51 | -49.99 | -45.10 | -25.97 | 7.41 | SkewedGaussian(-55.646; 10.282; 2.047) |
|  | 5 | 84 | -48.32 | 7.34 | -70.87 | -53.30 | -47.17 | -43.45 | -30.56 | 9.85 | SkewedGaussian(-41.556; 9.953; -1.627) |
|  | 6 | 23 | -52.13 | 9.92 | -86.16 | -55.06 | -49.10 | -45.46 | -41.25 | 9.60 | SkewedGaussian(-40.962; 15.229; -3.589) |
| Saturation | 2/3 | 51 | 119.71 | 34.72 | 24.36 | 94.58 | 125.04 | 145.59 | 192.48 | 51.01 | N(119.712; 34.377) |
|  | 5 | 84 | 122.92 | 34.26 | 65.44 | 96.52 | 119.28 | 145.16 | 212.65 | 48.63 | LogNormal(-0.613; 0.276; 118.957) |
|  | 6 | 22 | 108.05 | 36.28 | 62.66 | 83.24 | 100.15 | 118.96 | 200.95 | 35.72 | SkewedGaussian(62.656; 57.594; 4023916.033) |
| SFA | 2/3 | 59 | 0.07 | 0.06 | 0.00 | 0.04 | 0.06 | 0.08 | 0.32 | 0.04 | SkewedGaussian(0.002; 0.092; 35866938.756) |
|  | 5 | 105 | 0.09 | 0.05 | 0.00 | 0.06 | 0.08 | 0.13 | 0.23 | 0.07 | SkewedGaussian(0.029; 0.079; 4.415) |
|  | 6 | 27 | 0.09 | 0.05 | 0.01 | 0.06 | 0.08 | 0.11 | 0.21 | 0.05 | SkewedGaussian(0.04; 0.07; 3.21) |

**Table 20:**
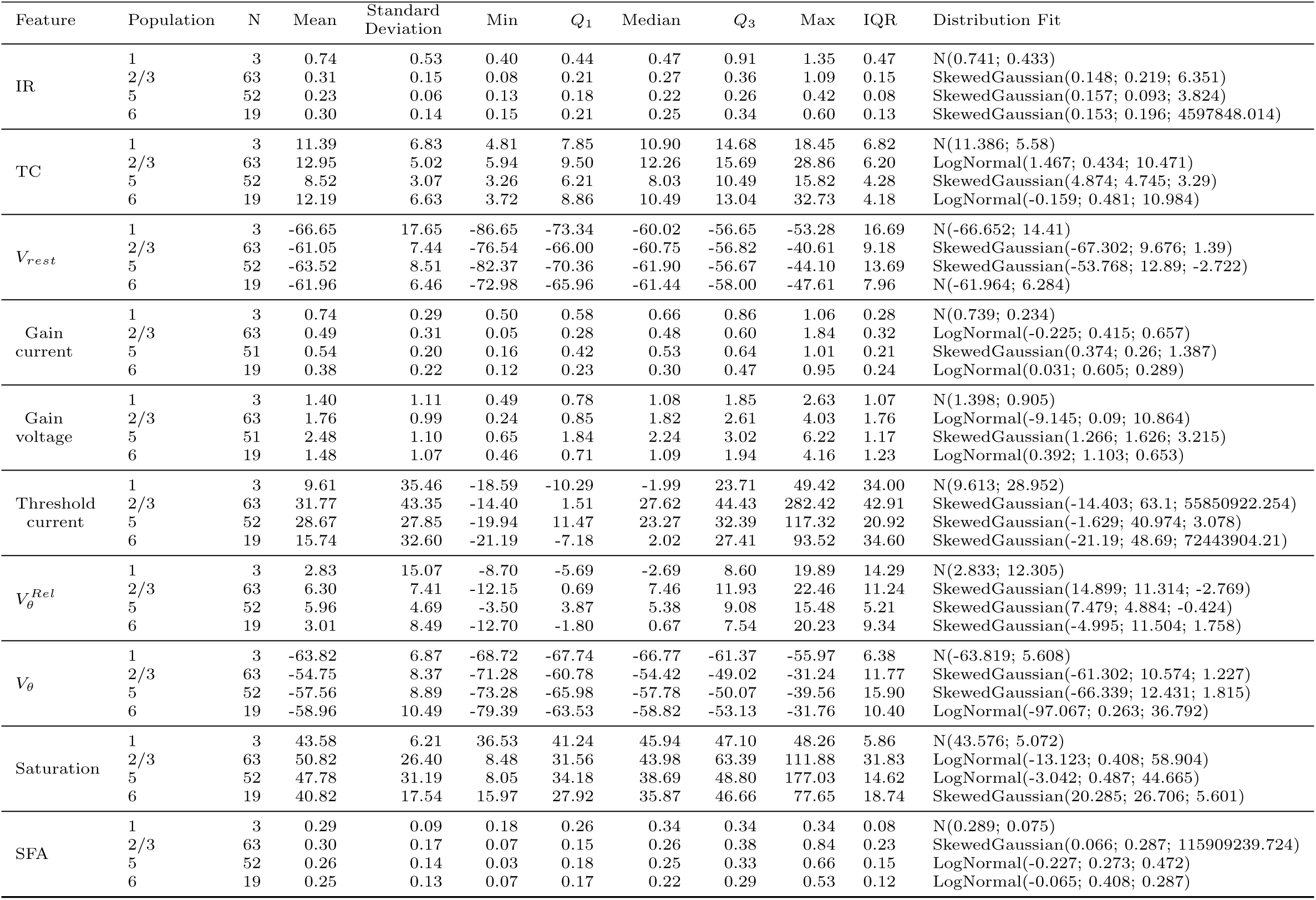
Table of descriptive statistics and fitting parameters for figure 20. **IR**: Input Resistance (*G*Ω),**TC**: Time Constant(ms), *V_rest_*: Resting membrane potential (mV), **Gain current** (Hz/pA),**Gain voltage**: (Hz/mV), **Threshold current**: (pA), *V ^Rel^*: Threshold Voltage (mV), *V_θ_*: Threshold Absolute Voltage (mV), **Saturation**: Saturation Frequency (Hz), **SFA**: Spike Frequency Adaptation (no unit)

**Table 21:** Table of descriptive statistics and fitting parameters for figure 21. **IR**: Input Resistance (*G*Ω),**TC**: Time Constant(ms), *V_rest_*: Resting membrane potential (mV), **Gain current** (Hz/pA),**Gain voltage**: (Hz/mV), **Threshold current**: (pA), *V ^Rel^*: Threshold Voltage (mV), *V_θ_*: Threshold Absolute Voltage (mV), **Saturation**: Saturation Frequency (Hz), **SFA**: Spike Frequency Adaptation (no unit)

| Feature | Population | N | Mean | Standard Deviation | Min | $Q_1$ | Median | $Q_3$ | Max | IQR | Distribution Fit |
| --- | --- | --- | --- | --- | --- | --- | --- | --- | --- | --- | --- |
| IR | 5 | 18 | 0.19 | 0.06 | 0.12 | 0.15 | 0.18 | 0.20 | 0.33 | 0.05 | LogNormal(0.099; 0.61; 0.074) |
|  | 6 | 6 | 0.22 | 0.07 | 0.17 | 0.19 | 0.21 | 0.22 | 0.35 | 0.03 | LogNormal(0.156; 0.923; 0.046) |
| TC | 5 | 18 | 15.93 | 5.63 | 7.73 | 12.32 | 14.48 | 19.06 | 28.19 | 6.74 | LogNormal(2.367; 0.401; 12.527) |
|  | 6 | 6 | 15.38 | 6.58 | 7.38 | 10.84 | 15.46 | 18.30 | 25.45 | 7.46 | N(15.383; 6.004) |
| $V_{rest}$ | 5 | 18 | -64.84 | 6.34 | -75.93 | -69.30 | -65.46 | -60.42 | -53.04 | 8.88 | SkewedGaussian(-70.925; 8.66; 1.788) |
|  | 6 | 6 | -66.13 | 6.41 | -73.26 | -71.42 | -66.24 | -60.42 | -59.41 | 10.99 | N(-66.13; 5.854) |
| Gain current | 5 | 18 | 0.22 | 0.16 | 0.04 | 0.11 | 0.16 | 0.28 | 0.62 | 0.16 | LogNormal(0.007; 0.811; 0.16) |
|  | 6 | 6 | 0.25 | 0.17 | 0.09 | 0.17 | 0.20 | 0.26 | 0.57 | 0.09 | LogNormal(0.048; 0.751; 0.156) |
| Gain voltage | 5 | 18 | 1.17 | 0.75 | 0.23 | 0.53 | 1.02 | 1.59 | 2.76 | 1.05 | SkewedGaussian(0.231; 1.194; 40179446.569) |
|  | 6 | 6 | 1.14 | 0.80 | 0.55 | 0.79 | 0.85 | 1.04 | 2.75 | 0.25 | LogNormal(0.505; 1.153; 0.353) |
| Threshold current | 5 | 18 | 68.26 | 34.14 | -6.90 | 46.14 | 62.12 | 81.42 | 131.72 | 35.27 | SkewedGaussian(39.415; 43.964; 1.447) |
|  | 6 | 6 | 38.45 | 32.54 | -13.98 | 26.34 | 38.47 | 60.79 | 77.22 | 34.45 | N(38.452; 29.708) |
| $V_{\theta}^{Rel}$ | 5 | 18 | 11.56 | 5.13 | -2.30 | 9.22 | 10.93 | 15.11 | 19.56 | 5.89 | SkewedGaussian(16.68; 7.141; -2.115) |
|  | 6 | 6 | 7.04 | 6.72 | -4.94 | 5.66 | 7.76 | 11.73 | 13.63 | 6.06 | N(7.037; 6.132) |
| $V_{\theta}$ | 5 | 18 | -53.28 | 5.02 | -60.97 | -56.59 | -53.14 | -50.31 | -43.21 | 6.28 | N(-53.276; 4.874) |
|  | 6 | 6 | -59.09 | 5.68 | -66.22 | -63.22 | -58.00 | -56.83 | -50.94 | 6.39 | SkewedGaussian(-59.104; 5.181; 0.003) |
| Saturation | 5 | 18 | 39.49 | 36.54 | 18.80 | 26.43 | 29.54 | 33.12 | 177.76 | 6.69 | LogNormal(17.417; 0.938; 12.599) |
|  | 6 | 6 | 36.48 | 19.33 | 22.64 | 24.05 | 27.63 | 41.40 | 72.11 | 17.35 | N(36.476; 17.645) |
| SFA | 5 | 18 | 0.30 | 0.10 | 0.06 | 0.24 | 0.29 | 0.35 | 0.46 | 0.12 | SkewedGaussian(0.399; 0.144; -1.945) |
|  | 6 | 6 | 0.24 | 0.10 | 0.11 | 0.19 | 0.23 | 0.30 | 0.40 | 0.12 | SkewedGaussian(0.13; 0.148; 3.226) |

## References

H. Akoglu. User’s guide to correlation coefficients. Turkish Journal of Emergency Medicine, 18:91–93, 9 2018. ISSN 24522473. doi: 10.1016/j.tjem.2018.08.001.

G. A. Ascoli, L. Alonso-Nanclares, S. A. Anderson, G. Barrionuevo, R. Benavides-Piccione, A. Burkhalter, G. Buzsáki, B. Cauli, J. DeFelipe, E. P. Gardner, J. H. Goldberg, M. Helmstaedter, S. Hestrin, F. Karube, Z. F. Kisvárday, B. Lambolez, D. A. Lewis, O. Marin, H. Markram, A. Munôz, A. Packer, C. C. Petersen, K. S. Rockland, J. Rossier, B. Rudy, P. Somogyi, J. F. Staiger, G. Tamas, A. M. Thomson, M. Toledo-Rodriguez, Y. Wang, D. C. West, and R. Yuste. Petilla terminology: nomenclature of features of gabaergic interneurons of the cerebral cortex. Nature Reviews Neuroscience, 9:557–568, 2008.

J. Benda and A. V. M. Herz. A universal model for spike-frequency adaptation. Neural Computation, 15: 2523–2564, 2003.

B. W. Connors and M. J. Gutnick. Intrinsic firing patterns of diverse neocortical neurons. Trends in neurosciences, 13, 1990.

D. Contreras. Electrophysiological classes of neocortical neurons. Neural Networks, 17:633–646, 2004. ISSN 08936080. doi: 10.1016/j.neunet.2004.04.003.

T. Crockett, N. Wright, S. Thornquist, M. Ariel, and R. Wessel. Turtle dorsal cortex pyramidal neurons comprise two distinct cell types with indistinguishable visual responses. PLoS ONE, 10, 12 2015. ISSN 19326203. doi: 10.1371/journal.pone.0144012.

A. da Silva Lantyer, N. Calcini, A. Bijlsma, K. Kole, M. Emmelkamp, M. Peeters, W. J. Scheenen, F. Zeldenrust, and T. Celikel. A databank for intracellular electrophysiological mapping of the adult somatosensory cortex. GigaScience, 7, 12 2018. ISSN 2047217X. doi: 10.1093/gigascience/giy147.

J. Defelipe. Cortical interneurons: from cajal to 2001. Prog Brain Res., 2002.

J. Defelipe and I. Fariñas. The pyramidal neuron of the cerebral cortex: Morphological and chemical characteristics of the synaptic inputs. Progress in Neurobiology, 39:563, 1992.

A. Destexhe, M. Rudolph, and D. Paré. The high-conductance state of neocortical neurons in vivo. Nature Reviews Neuroscience, 4:739–751, 2003. ISSN 14710048. doi: 10.1038/nrn1198.

R. Duarte and A. Morrison. Leveraging heterogeneity for neural computation with fading memory in layer 2/3 cortical microcircuits. PLoS Computational Biology, 15, 2019. ISSN 15537358. doi: 10.1371/journal.pcbi.1006781.

F. R. Fernandez, B. Rahsepar, and J. A. White. Differences in the electrophysiological properties of mouse somatosensory layer 2/3 neurons in vivo and slice stem from intrinsic sources rather than a network-generated high conductance state. eNeuro, 5, 3 2018. ISSN 23732822. doi: 10.1523/ENEURO.0447-17.2018.

C. Findling and V. Wyart. Computation noise in human learning and decision-making: origin, impact, function, 4 2021. ISSN 23521546.

M. G. Frantz, E. C. Crouse, G. Sokhadze, T. Ikrar, C. Élise Stephany, C. Nguyen, X. Xu, and A. W. McGee. Layer 4 gates plasticity in visual cortex independent of a canonical microcircuit. Current Biology, 30: 2962–2973.e5, 8 2020. ISSN 18790445. doi: 10.1016/j.cub.2020.05.067.

I. Férézou, B. Cauli, E. L. Hill, J. Rossier, E. Hamel, and B. Lambolez. 5-ht 3 receptors mediate serotonergic fast synaptic excitation of neocortical vasoactive intestinal peptide/cholecystokinin interneurons. The journal of Neuroscience, 22, 9 2002.

C. R. Gerfen, R. Paletzki, and N. Heintz. Gensat bac cre-recombinase driver lines to study the functional organization of cerebral cortical and basal ganglia circuits. Neuron, 80:1368–1383, 12 2013. ISSN 10974199. doi: 10.1016/j.neuron.2013.10.016.

P. Ghaderi, H. R. Marateb, and M. S. Safari. Electrophysiological profiling of neocortical neural subtypes: A semi-supervised method applied to in vivo whole-cell patch-clamp data. Frontiers in Neuroscience, 12, 11 2018. ISSN 1662453X. doi: 10.3389/fnins.2018.00823.

N. W. Gouwens, S. A. Sorensen, J. Berg, C. Lee, T. Jarsky, J. Ting, S. M. Sunkin, D. Feng, C. A. Anastassiou, E. Barkan, K. Bickley, N. Blesie, T. Braun, K. Brouner, A. Budzillo, S. Caldejon, T. Casper, D. Castelli, P. Chong, K. Crichton, C. Cuhaciyan, T. L. Daigle, R. Dalley, N. Dee, T. Desta, S. L. Ding, S. Dingman, A. Doperalski, N. Dotson, T. Egdorf, M. Fisher, R. A. de Frates, E. Garren, M. Garwood, A. Gary, N. Gaudreault, K. Godfrey, M. Gorham, H. Gu, C. Habel, K. Hadley, J. Harrington, J. A. Harris, A. Henry, D. J. Hill, S. Josephsen, S. Kebede, L. Kim, M. Kroll, B. Lee, T. Lemon, K. E. Link, X. Liu, B. Long, R. Mann, M. McGraw, S. Mihalas, A. Mukora, G. J. Murphy, L. Ng, K. Ngo, T. N. Nguyen, P. R. Nicovich, A. Oldre, D. Park, S. Parry, J. Perkins, L. Potekhina, D. Reid, M. Robertson, D. Sandman, M. Schroedter, C. Slaughterbeck, G. Soler-Llavina, J. Sulc, A. Szafer, B. Tasic, N. Taskin, C. Teeter, N. Thatra, H. Tung, W. Wakeman, G. Williams, R. Young, Z. Zhou, C. Farrell, H. Peng, M. J. Hawrylycz, E. Lein, L. Ng, A. Arkhipov, A. Bernard, J. W. Phillips, H. Zeng, and C. Koch. Classification of electrophysiological and morphological neuron types in the mouse visual cortex. Nature Neuroscience, 22:1182–1195, 7 2019. ISSN 15461726. doi: 10.1038/s41593-019-0417-0.

N. W. Gouwens, S. A. Sorensen, F. Baftizadeh, A. Budzillo, B. R. Lee, T. Jarsky, L. Alfiler, K. Baker, E. Barkan, K. Berry, D. Bertagnolli, K. Bickley, J. Bomben, T. Braun, K. Brouner, T. Casper, K. Crichton, T. L. Daigle, R. Dalley, R. A. de Frates, N. Dee, T. Desta, S. D. Lee, N. Dotson, T. Egdorf, L. Ellingwood, R. Enstrom, L. Esposito, C. Farrell, D. Feng, O. Fong, R. Gala, C. Gamlin, A. Gary, A. Glandon, J. Goldy, M. Gorham, L. Graybuck, H. Gu, K. Hadley, M. J. Hawrylycz, A. M. Henry, D. J. Hill, M. Hupp, S. Kebede, T. K. Kim, L. Kim, M. Kroll, C. Lee, K. E. Link, M. Mallory, R. Mann, M. Maxwell, M. McGraw, D. McMillen, A. Mukora, L. Ng, L. Ng, K. Ngo, P. R. Nicovich, A. Oldre, D. Park, H. Peng, O. Penn, T. Pham, A. Pom, Z. Popović, L. Potekhina, R. Rajanbabu, S. Ransford, D. Reid, C. Rimorin, M. Robertson, K. Ronellenfitch, A. Ruiz, D. Sandman, K. Smith, J. Sulc, S. M. Sunkin, A. Szafer, M. Tieu, A. Torkelson, J. Trinh, H. Tung, W. Wakeman, K. Ward, G. Williams, Z. Zhou, J. T. Ting, A. Arkhipov, U. Sümbül, E. S. Lein, C. Koch, Z. Yao, B. Tasic, J. Berg, G. J. Murphy, and H. Zeng. Integrated morphoelectric and transcriptomic classification of cortical gabaergic cells. Cell, 183:935–953.e19, 11 2020. ISSN 10974172. doi: 10.1016/j.cell.2020.09.057.

C. Halfmann, T. Rüland, F. Müller, K. Jehasse, and B. M. Kampa. Electrophysiological properties of layer 2/3 pyramidal neurons in the primary visual cortex of a retinitis pigmentosa mouse model (rd10). Frontiers in Cellular Neuroscience, 17, 9 2023. ISSN 16625102. doi: 10.3389/fncel.2023.1258773.

P. M. Harrison, L. Badel, M. J. Wall, and M. J. Richardson. Experimentally verified parameter sets for modelling heterogeneous neocortical pyramidal-cell populations. PLoS Computational Biology, 11, 8 2015. ISSN 15537358. doi: 10.1371/journal.pcbi.1004165.

A. M. Hattox and S. B. Nelson. Layer v neurons in mouse cortex projecting to different targets have distinct physiological properties. Journal of Neurophysiology, 98:3330–3340, 12 2007. ISSN 00223077. doi: 10.1152/jn.00397.2007.

S. H. C. Hendry and E. G. Jones. Gaba neuronal subpopulations in cat primary auditory cortex: colocalization with calcium binding proteins. Brain Research, 543:45–55, 1991.

S. H. C. Hendry, E. G. Jones, P. C. Emson, D. E. M. Lawson, C. W. Heizmann, and P. Streit. Two classes of cortical gaba neurons defined by differential calcium binding protein immunoreactivities. Experimental Brain Research, 76:467–472, 1989.

A. Hodgkin and A. Huxley. Hodgkin and huxley 1939. Nature, 10 1939.

A. L. Hodgkin. The local electric changes associated with repetitive action in a non-medullated axon. J. Physiol., 07:65–73, 1948.

A. L. Hodgkin and A. F. Huxley. A quantitative description of membrane current and its application to conduction and excitation in nerve. J. Physiol, pages 500–544, 1952.

B. A. L. Hodgkin, B. Katz, and M. B. Association. The effect of sodium ions on the electrical activity of the giant axon of the squid from the laboratory of the. J. Physiol. (I949), 8:37–77, 1949.

M. J. V. Hook. Temperature effects on synaptic transmission and neuronal function in the visual thalamus. PLoS ONE, 15, 4 2020. ISSN 19326203. doi: 10.1371/journal.pone.0232451.

L. Huang, P. Ledochowitsch, U. Knoblich, Jrôme Lecoq, G. J. Murphy, R. C. Reid, S. E. de Vries, C. Koch, H. Zeng, M. A. Buice, J. Waters, and L. Li. Relationship between simultaneously recorded spiking activity and fluorescence signal in gcamp6 transgenic mice. eLife, 2021. doi: 10.6080/K02R3PMN. URL 10.6080/K02R3PMN.

S. Huggenberger, M. Vater, and R. A. Deisz. Interlaminar differences of intrinsic properties of pyramidal neurons in the auditory cortex of mice. Cerebral Cortex, 19:1008–1018, 5 2009. ISSN 10473211. doi: 10.1093/cercor/bhn143.

A. Hutt, S. I. Rich, T. A. Valiante, and J. Lefebvre. Intrinsic neural diversity quenches the dynamic volatility of neural networks. 120, 2023. doi: 10.1073/pnas.

M. Janssens, S. Gaillard, J. de Haan, W. de Leeuw, M. Brooke, M. Burke, J. Flores, I. Kruijen, J. Menon, A. Smith, I. Tiebosch, and F. Weijdema. How open science can support the 3rs and improve animal research. Research Ideas and Outcomes, 9, 8 2023. doi: 10.3897/rio.9.e105198.

E. J. Kim, A. L. Juavinett, E. M. Kyubwa, M. W. Jacobs, and E. M. Callaway. Three types of cortical layer 5 neurons that differ in brain-wide connectivity and function. Neuron, 88:1253–1267, 2015. ISSN 10974199. doi: 10.1016/j.neuron.2015.11.002.

A. L. Lamprecht, L. Garcia, M. Kuzak, C. Martinez, R. Arcila, E. M. D. Pico, V. D. D. Angel, S. V. D. Sandt, J. Ison, P. A. Martinez, P. McQuilton, A. Valencia, J. Harrow, F. Psomopoulos, J. L. Gelpi, N. C. Hong, C. Goble, and S. Capella-Gutierrez. Towards fair principles for research software. Data Science, 3: 37–59, 6 2020. ISSN 24518492. doi: 10.3233/DS-190026.

B. R. Lee, A. Budzillo, K. Hadley, J. A. Miller, T. Jarsky, K. Baker, D. Hill, L. Kim, R. Mann, L. Ng, Oldre, R. Rajanbabu, J. Trinh, S. Vargas, T. Braun, R. A. Dalley, N. W. Gouwens, B. E. Kalmbach, T. K. Kim, K. A. Smith, G. Soler-Llavina, S. Sorensen, B. Tasic, J. T. Ting, E. Lein, H. Zeng, G. J. Murphy, and J. Berg. Scaled, high fidelity electrophysiological, morphological, and transcriptomic cell characterization. doi: 10.7554/eLife.

S. H. Lee, J. Hjerling-Leffler, E. Zagha, G. Fishell, and B. Rudy. The largest group of superficial neocortical gabaergic interneurons expresses ionotropic serotonin receptors. Journal of Neuroscience, 30:16796–16808, 12 2010. ISSN 02706474. doi: 10.1523/JNEUROSCI.1869-10.2010.

X. Mao and J. F. Staiger. Multimodal cortical neuronal cell type classification, 5 2024. ISSN 14322013.

H. Markram, M. Toledo-Rodriguez, Y. Wang, A. Gupta, G. Silberberg, and C. Wu. Interneurons of the neocortical inhibitory system, 10 2004. ISSN 1471003X.

G. Milior, M. A. D. Castro, L. P. Sciarria, S. Garofalo, I. Branchi, D. Ragozzino, C. Limatola, and L. Maggi. Electrophysiological properties of ca1 pyramidal neurons along the longitudinal axis of the mouse hippocampus. Scientific Reports, 6, 12 2016. ISSN 20452322. doi: 10.1038/srep38242.

M. Mitrić, A. Seewald, G. Moschetti, P. Sacerdote, F. Ferraguti, K. K. Kummer, and M. Kress. Layer- and subregion-specific electrophysiological and morphological changes of the medial prefrontal cortex in a mouse model of neuropathic pain. Scientific Reports, 9, 12 2019. ISSN 20452322. doi: 10.1038/s41598-019-45677-z.

M. Morales and F. E. Bloom. The 5-ht 3 receptor is present in different subpopulations of gabaergic neurons in the rat telencephalon binding sites have been found consistently in cortex, amygdala, and hippocampus (kilpatrick et al. The journal of neuroscience, 1997.

M. Munz, A. Bharioke, G. Kosche, V. Moreno-Juan, A. Brignall, T. M. Rodrigues, A. Graff-Meyer, T. Ulmer, S. Haeuselmann, D. Pavlinic, N. Ledergerber, B. Gross-Scherf, B. Rózsa, J. Krol, S. Picelli, C. S. Cowan, and B. Roska. Pyramidal neurons form active, transient, multilayered circuits perturbed by autism-associated mutations at the inception of neocortex. Cell, 186:1930–1949.e31, 4 2023. ISSN 10974172. doi: 10.1016/j.cell.2023.03.025.

S. P. Muscinelli, W. Gerstner, and T. Schwalger. How single neuron properties shape chaotic dynamics and signal transmission in random neural networks. PLoS Computational Biology, 15, 6 2019. ISSN 15537358. doi: 10.1371/journal.pcbi.1007122.

A. Nandi, T. Chartrand, W. V. Geit, A. Buchin, Z. Yao, S. Y. Lee, Y. Wei, B. Kalmbach, B. Lee, E. Lein, J. Berg, U. Sümbül, C. Koch, B. Tasic, and C. A. Anastassiou. Single-neuron models linking electrophysiology, morphology, and transcriptomics across cortical cell types. Cell Reports, 40, 8 2022. ISSN 22111247. doi: 10.1016/j.celrep.2022.111176.

O. Ophir, O. Shefi, and O. Lindenbaum. Classifying neuronal cell types based on shared electrophysiological information from humans and mice. Neuroinformatics, 2024. ISSN 15590089. doi: 10.1007/s12021-024-09675-5.

K. Padmanabhan and N. N. Urban. Intrinsic biophysical diversity decorrelates neuronal firing while increasing information content. Nature Neuroscience, 13:1276–1282, 10 2010. ISSN 10976256. doi: 10.1038/nn.2630.

G. Pan, J. M. Yang, X. Y. Hu, and X. M. Li. Postnatal development of the electrophysiological properties of somatostatin interneurons in the anterior cingulate cortex of mice. Scientific Reports, 6, 6 2016. ISSN 20452322. doi: 10.1038/srep28137.

H. Pastoll, D. Garden, I. Papastathopoulos, G. Sürmeli, and M. F. Nolan. Interand intraanimal variation in the integrative properties of stellate cells in the medial entorhinal cortex. eLife, 9, 2 2020. ISSN 2050084X. doi: 10.7554/eLife.52258.

S. P. Peron and F. Gabbiani. Role of spike-frequency adaptation in shaping neuronal response to dynamic stimuli. Biological Cybernetics, 100:505–520, 6 2009. ISSN 03401200. doi: 10.1007/s00422-009-0304-y.

P. C. Petersen, M. Voroslakos, and G. Buzsáki. Brain temperature affects quantitative features of hippocampal sharp wave ripples. Journal of Neurophysiology, 127:1417–1425, 5 2022. ISSN 15221598. doi: 10.1152/jn.00047.2022.

A. Prönneke, B. Scheuer, R. J. Wagener, M. Möck, M. Witte, and J. F. Staiger. Characterizing vip neurons in the barrel cortex of vipcre/tdtomato mice reveals layer-specific differences. Cerebral Cortex, 25:4854–4868, 12 2015. ISSN 14602199. doi: 10.1093/cercor/bhv202.

S. Rich, H. M. Chameh, J. Lefebvre, and T. A. Valiante. Loss of neuronal heterogeneity in epileptogenic human tissue impairs network resilience to sudden changes in synchrony. Cell Reports, 39, 5 2022. ISSN 22111247. doi: 10.1016/j.celrep.2022.110863.

S. H. Richter. Challenging current scientific practice: how a shift in research methodology could reduce animal use, 1 2024. ISSN 15484475.

D. Rodarie, C. Verasztó, Y. Roussel, M. Reimann, D. Keller, S. Ramaswamy, H. Markram, and M. O. Gewaltig. A method to estimate the cellular composition of the mouse brain from heterogeneous datasets. PLoS Computational Biology, 18, 12 2022. ISSN 15537358. doi: 10.1371/journal.pcbi.1010739.

A. Rodríguez-Collado and C. Rueda. Electrophysiological and transcriptomic features reveal a circular taxonomy of cortical neurons. Frontiers in Human Neuroscience, 15, 7 2021. ISSN 16625161. doi: 10.3389/fnhum.2021.684950.

A. Roxin, N. Brune, D. Hansel, G. Mongillo, and C. van Vreeswijk. On the distribution of firing rates in networks of cortical neurons. Journal of Neuroscience, 31:16217–16226, 11 2011. ISSN 02706474. doi: 10.1523/JNEUROSCI.1677-11.2011.

B. Rudy, G. Fishell, S. H. Lee, and J. Hjerling-Leffler. Three groups of interneurons account for nearly 100 *Developmental Neurobiology*, 71:45–61, 1 2011. ISSN 19328451. doi: 10.1002/dneu.20853.

O. Rübel, A. Tritt, R. Ly, B. K. Dichter, S. Ghosh, L. Niu, P. Baker, I. Soltesz, L. Ng, K. Svoboda, L. Frank, and K. E. Bouchard. The neurodata without borders ecosystem for neurophysiological data science. eLife, 11:78362, 2022. doi: 10.7554/eLife.

F. Scala, D. Kobak, S. Shan, Y. Bernaerts, S. Laturnus, C. R. Cadwell, L. Hartmanis, E. Froudarakis, J. R. Castro, Z. H. Tan, S. Papadopoulos, S. S. Patel, R. Sandberg, P. Berens, X. Jiang, and A. S. Tolias. Layer 4 of mouse neocortex differs in cell types and circuit organization between sensory areas. Nature Communications, 10, 12 2019. ISSN 20411723. doi: 10.1038/s41467-019-12058-z.

F. Scala, D. Kobak, M. Bernabucci, Y. Bernaerts, C. R. Cadwell, J. R. Castro, L. Hartmanis, X. Jiang, S. Laturnus, E. Miranda, S. Mulherkar, Z. H. Tan, Z. Yao, H. Zeng, R. Sandberg, P. Berens, and A. S. Tolias. Phenotypic variation of transcriptomic cell types in mouse motor cortex. Nature, 598:144–150, 10 2021. ISSN 14764687. doi: 10.1038/s41586-020-2907-3.

M. Shamir and H. Sompolinsky. Implications of neuronal diversity on population coding. Neural Computation, 18, 2006.

S. Shinomoto, K. Shima, and J. Tanji. Differences in spiking patterns among cortical neurons. Neural Computation, 15:2832–2842, 2003.

S. Shinomoto, H. Kim, T. Shimokawa, N. Matsuno, S. Funahashi, K. Shima, I. Fujita, H. Tamura, T. Doi, K. Kawano, N. Inaba, K. Fukushima, S. Kurkin, K. Kurata, M. Taira, K. I. Tsutsui, H. Komatsu, T. Ogawa, K. Koida, J. Tanji, and K. Toyama. Relating neuronal firing patterns to functional differentiation of cerebral cortex. PLoS Computational Biology, 5, 2009. ISSN 15537358. doi: 10.1371/journal.pcbi.1000433.

N. P. Staff, H.-Y. Jung, T. Thiagarajan, M. Yao, and N. Spruston. Resting and active properties of pyramidal neurons in subiculum and ca1 of rat hippocampus. Journal of Physiology, 2000. URL www.jn.physiology.org.

F. Tang, C. Barbacioru, Y. Wang, E. Nordman, C. Lee, N. Xu, X. Wang, J. Bodeau, B. B. Tuch, A. Siddiqui, K. Lao, and M. A. Surani. mrna-seq whole-transcriptome analysis of a single cell. Nature Methods, 6: 377–382, 2009. ISSN 15487091. doi: 10.1038/nmeth.1315.

B. Tasic, V. Menon, T. N. Nguyen, T. K. Kim, T. Jarsky, Z. Yao, B. Levi, L. T. Gray, S. A. Sorensen, T. Dolbeare, D. Bertagnolli, J. Goldy, N. Shapovalova, S. Parry, C. Lee, K. Smith, A. Bernard, L. Madisen, S. M. Sunkin, M. Hawrylycz, C. Koch, and H. Zeng. Adult mouse cortical cell taxonomy revealed by single cell transcriptomics. Nature Neuroscience, 19:335–346, 1 2016. ISSN 15461726. doi: 10.1038/nn.4216.

B. Tasic, Z. Yao, L. T. Graybuck, K. A. Smith, T. N. Nguyen, D. Bertagnolli, J. Goldy, E. Garren, M. N. Economo, S. Viswanathan, O. Penn, T. Bakken, V. Menon, J. Miller, O. Fong, K. E. Hirokawa, K. Lathia, C. Rimorin, M. Tieu, R. Larsen, T. Casper, E. Barkan, M. Kroll, S. Parry, N. V. Shapovalova, D. Hirschstein, J. Pendergraft, H. A. Sullivan, T. K. Kim, A. Szafer, N. Dee, P. Groblewski, I. Wickersham, A. Cetin, J. A. Harris, B. P. Levi, S. M. Sunkin, L. Madisen, T. L. Daigle, L. Looger, A. Bernard, J. Phillips, E. Lein, M. Hawrylycz, K. Svoboda, A. R. Jones, C. Koch, and H. Zeng. Shared and distinct transcriptomic cell types across neocortical areas. Nature, 563:72–78, 11 2018. ISSN 14764687. doi: 10.1038/s41586-018-0654-5.

C. Teeter, R. Iyer, V. Menon, N. Gouwens, D. Feng, J. Berg, A. Szafer, N. Cain, H. Zeng, M. Hawrylycz, C. Koch, and S. Mihalas. Generalized leaky integrate-and-fire models classify multiple neuron types. Nature Communications, 9, 12 2018. ISSN 20411723. doi: 10.1038/s41467-017-02717-4.

J. L. Teeters, K. Godfrey, R. Young, C. Dang, C. Friedsam, B. Wark, H. Asari, S. Peron, N. Li, A. Peyrache, G. Denisov, J. H. Siegle, S. R. Olsen, C. Martin, M. Chun, S. Tripathy, T. J. Blanche, K. Harris, G. Buzsáki, C. Koch, M. Meister, K. Svoboda, and F. T. Sommer. Neurodata without borders: Creating a common data format for neurophysiology, 11 2015. ISSN 10974199.

R. Tremblay, S. Lee, and B. Rudy. Gabaergic interneurons in the neocortex: From cellular properties to circuits, 7 2016. ISSN 10974199.

Y. Ueta, T. Otsuka, M. Morishima, M. Ushimaru, and Y. Kawaguchi. Multiple layer 5 pyramidal cell subtypes relay cortical feedback from secondary to primary motor areas in rats. Cerebral Cortex, 24:2362–2376, 2014. ISSN 14602199. doi: 10.1093/cercor/bht088.

M. Valero, A. Navas-Olive, L. M. de la Prida, and G. Buzsáki. Inhibitory conductance controls place field dynamics in the hippocampus. Cell Reports, 40, 8 2022. ISSN 22111247. doi: 10.1016/j.celrep.2022.111232.

P. Vitale, A. R. Salgueiro-Pereira, C. A. Lupascu, M. Willem, R. Migliore, M. Migliore, and H. Marie. Analysis of age-dependent alterations in excitability properties of ca1 pyramidal neurons in an appps1 model of alzheimer’s disease. Frontiers in Aging Neuroscience, 13, 6 2021. ISSN 16634365. doi: 10.3389/fnagi.2021.668948.

L. Waschke, N. A. Kloosterman, J. Obleser, and D. D. Garrett. Behavior needs neural variability, 3 2021. ISSN 10974199.

M. D. Wilkinson, M. Dumontier, I. J. Aalbersberg, G. Appleton, M. Axton, A. Baak, N. Blomberg, J. W. Boiten, L. B. da Silva Santos, P. E. Bourne, J. Bouwman, A. J. Brookes, T. Clark, M. Crosas, I. Dillo, O. Dumon, S. Edmunds, C. T. Evelo, R. Finkers, A. Gonzalez-Beltran, A. J. Gray, P. Groth, C. Goble, J. S. Grethe, J. Heringa, P. A. t Hoen, R. Hooft, T. Kuhn, R. Kok, J. Kok, S. J. Lusher, M. E. Martone, A. Mons, A. L. Packer, B. Persson, P. Rocca-Serra, M. Roos, R. van Schaik, S. A. Sansone, E. Schultes, T. Sengstag, T. Slater, G. Strawn, M. A. Swertz, M. Thompson, J. V. D. Lei, E. V. Mulligen, J. Velterop, A. Waagmeester, P. Wittenburg, K. Wolstencroft, J. Zhao, and B. Mons. Comment: The fair guiding principles for scientific data management and stewardship. Scientific Data, 3, 3 2016. ISSN 20524463. doi: 10.1038/sdata.2016.18.

C. G. Williams, H. J. Lee, T. Asatsuma, R. Vento-Tormo, and A. Haque. An introduction to spatial transcriptomics for biomedical research. Genome Medicine, 14, 12 2022. ISSN 1756994X. doi: 10.1186/s13073-022-01075-1.

X. Xu, K. D. Roby, and E. M. Callaway. Immunochemical characterization of inhibitory mouse cortical neurons: Three chemically distinct classes of inhibitory cells. Journal of Comparative Neurology, 518: 389–404, 2 2010. ISSN 00219967. doi: 10.1002/cne.22229.

A. Zeisel, A. B. Moz-Manchado, S. Codeluppi, P. Lannerberg, G. L. Manno, A. Juraus, S. Marques, H. Munguba, L. He, C. Betsholtz, C. Rolny, G. Castelo-Branco, J. Hjerling-Leffler, and S. Linnarsson. Cell types in the mouse cortex and hippocampus revealed by single-cell rna-seq. Scienceexpress, 347: 1138–1142, 3 2015. ISSN 10959203. doi: 10.1126/science.aaa1934.

H. Zeng. What is a cell type and how to define it?, 7 2022. ISSN 10974172.

H. Zeng and J. R. Sanes. Neuronal cell-type classification: Challenges, opportunities and the path forward. Nature Reviews Neuroscience, 18:530–546, 8 2017. ISSN 14710048. doi: 10.1038/nrn.2017.85.

